# Proteogenomics refines the molecular classification of chronic lymphocytic leukemia

**DOI:** 10.1101/2022.03.01.481539

**Authors:** Sophie A. Herbst, Mattias Vesterlund, Alexander J. Helmboldt, Rozbeh Jafari, Ioannis Siavelis, Matthias Stahl, Eva C. Schitter, Nora Liebers, Berit J. Brinkmann, Felix Czernilofsky, Tobias Roider, Peter-Martin Bruch, Murat Iskar, Adam Kittai, Ying Huang, Junyan Lu, Sarah Richter, Georgios Mermelekas, Husen Muhammad Umer, Mareike Knoll, Carolin Kolb, Angela Lenze, Xiaofang Cao, Cecilia Österholm, Linus Wahnschaffe, Carmen Herling, Sebastian Scheinost, Matthias Ganzinger, Larry Mansouri, Katharina Kriegsmann, Mark Kriegsmann, Simon Anders, Marc Zapatka, Giovanni Del Poeta, Antonella Zucchetto, Riccardo Bomben, Valter Gattei, Peter Dreger, Jennifer Woyach, Marco Herling, Carsten Müller-Tidow, Richard Rosenquist, Stephan Stilgenbauer, Thorsten Zenz, Wolfgang Huber, Eugen Tausch, Janne Lehtiö, Sascha Dietrich

## Abstract

Cancer heterogeneity at the proteome level may explain differences in therapy response and prognosis beyond the currently established genomic and transcriptomic based diagnostics. The relevance of proteomics for disease classifications remains to be established in clinically heterogeneous cancer entities such as chronic lymphocytic leukemia (CLL). Here, we characterized the proteome and transcriptome in-depth alongside genetic and *ex-vivo* drug response profiling in a clinically well annotated CLL discovery cohort (n= 68). Unsupervised clustering of the proteome data revealed six subgroups. Five of these proteomic groups were associated with genetic features, while one group was only detectable at the proteome level. This new group was characterized by accelerated disease progression, high spliceosomal protein abundances associated with aberrant splicing, and low B cell receptor signaling protein abundances (ASB-CLL). We developed classifiers to identify ASB-CLL based on its characteristic proteome or splicing signature in two independent cohorts (n= 165, n= 169) and confirmed that ASB-CLL comprises about 20 % of CLL patients. The inferior overall survival observed in ASB-CLL was independent of both *TP53-* and IGHV mutation status. Our multi-omics analysis refines the classification of CLL and highlights the potential of proteomics to improve cancer patient stratification beyond genetic and transcriptomic profiling.

**Single sentence summary:** We performed the largest proteogenomic analysis of CLL, linked proteomic profiles to clinical outcomes, and discovered a new poor outcome subgroup (ASB-CLL).

## Introduction

Chronic lymphocytic leukemia (CLL) is the most common adult leukemia in Western countries. It is characterized by the accumulation of mature B lymphocytes in the peripheral blood, the bone marrow, and lymph nodes. This incurable malignancy has a very heterogeneous clinical course. Some patients can be followed with a “watch and wait” strategy for many years, while others need frequent treatments and have a shorter overall survival ^1^.

The cell of origin of CLL represents an important source of heterogeneity, which is marked by the mutation status of the immunoglobulin heavy variable (IGHV) genes and characteristic methylation profiles ^2–4^. CLL with unmutated IGHV genes (U-CLL) cases progress faster and have worse outcomes than IGHV-mutated CLL (M-CLL) cases ^5^. These differences in clinical behavior between M-CLL and U-CLL are partially determined by differences in responsiveness to B cell receptor (BcR) stimulation ^6^. Inhibition of the BcR pathway has revolutionized the treatment landscape of CLL ^7^ and the prognostic difference between U-CLL and M-CLL is diminished in patients treated with BcR inhibitors ^8^.

Recently, the genetic landscape of CLL has been well characterized ^9^, which partially explains the heterogeneous disease courses. The most frequent recurrent somatic mutations in CLL alter genes encoding for components of a small set of oncogenic pathways^10^. These include the DNA damage response pathway, pathways that receive input from the microenvironment (NOTCH, Toll-like receptor and CD40 signaling), and pathways affecting ribosomal processing ^11^. Additionally, mutations in central splicing components, most frequently in *SF3B1*, are drivers of CLL ^12^. Recurrent structural chromosomal aberrations, including deletions of 13(q14), 11(q22-23), 17(p13), and trisomy 12, are routinely measured before the initiation of treatment and are detected in approximately 80 % of CLL patients. They are associated with diverse biological phenotypes and contrasting clinical behavior ^13^, but the pathophysiology of some of these variants such as trisomy 12 is still insufficiently understood.

A multi-omics analysis integrating the genome, transcriptome, and clinical outcomes could offer a valuable tool to fill this gap and improve the understanding of genetic variants and outcomes in CLL. Recent developments in mass spectrometry have facilitated deep proteomic profiling of multiple tumor specimens in parallel. The integration of proteomics with genomics and other data layers has started to enhance our understanding of selected cancer entities ^14–21^. However, the proteome-centric data integration to clinically relevant endpoints and functional phenotypes has been mostly lacking.

Although the proteome of CLL B cells has been compared to non-malignant normal B cells ^22–24^, a systematic description of the proteomic landscape of CLL integrating genome, transcriptome, and clinical datasets is currently missing. Here, we employed in-depth high-resolution isoelectric focusing liquid chromatography–mass spectrometry (HiRIEF LC-MS) ^25^ based proteomics to connect the proteomic landscape of a clinically well characterized and representative cohort of 68 CLL patients with genome, transcriptome, and drug perturbation profiling. Among the six identified proteome-based subgroups, we discovered a previously unknown subset of CLL patients with poor outcome. This subgroup was characterized by high abundance of spliceosomal proteins. For the first time, we could link spliceosomal protein abundances to aberrant splicing and poor outcome in CLL. We validated the subgroups in further independent cohorts using multiple validation strategies. Most importantly, we took advantage of the higher throughput of data-independent acquisition (DIA) proteomics to confirm our findings from the in-depth HiRIEF dataset. Including all patients used for the different validation strategies, our study contains data from 1503 CLL patients and directly links the observed molecular phenotypes to clinical outcome.

## Results

### Study outline

We assembled a discovery cohort of 68 well characterized CLL patients with a representative distribution of CLL subgroups ^9, 13^ (Fig 1, Fig. S1a, Supplementary table 1). Viably frozen CLL samples were CD19 enriched and split into three aliquots: one for proteomics, one for RNA-sequencing, and one for DNA panel sequencing ^26^. We further characterized functional phenotypes of the same CLL samples by *ex-vivo* drug response profiling using a high-throughput microscopy-based drug screening platform with a panel of 43 drugs, most of them FDA approved or currently in clinical trials (e.g. ibrutinib, venetoclax)^10^.

**Fig. 1:**
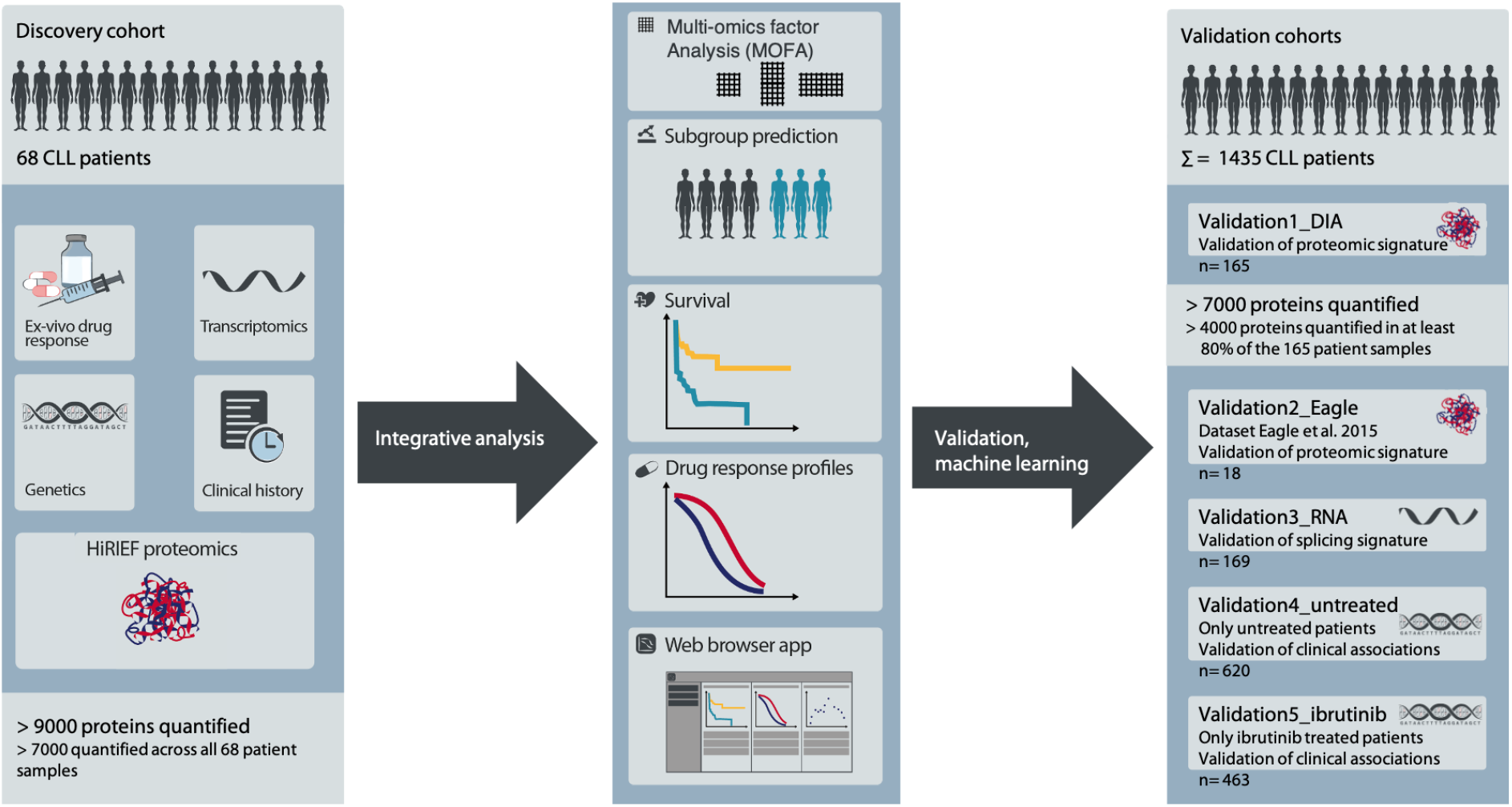
Overview of the study design. In-depth proteomics, transcriptomics, genetic and drug perturbation profiling were performed for a discovery cohort of 68 clinically well characterized CLL patients. An integrative analysis was performed to describe associations between the different molecular layers and to uncover patient subgroups. These were validated in multiple independent cohorts using different validation strategies. To validate proteomic signatures, we performed data-independent acquisition (DIA) proteomics on 165 CLL patient samples (Validation1_DIA) and took advantage of a published cohort of 18 CLL samples (Validation2_Eagle). For the validation of splicing signatures, 169 additional patients were characterized by RNA-sequencing (Validation3_RNA). We linked proteomic signatures with genetic profiles, which allowed us to validate associations of major biological axes in CLL with clinical outcome in 620 untreated (Validation4_untreated) and 463 ibrutinib treated CLL patients (Validation5_ibrutinib). In total this study analyzed data from 1503 CLL patients.

We used multiple validation strategies to confirm associations between genetic features, the proteome, splicing signatures and clinical outcome in independent CLL cohorts (Fig. 1, Supplementary table 1). This included data independent acquisition (DIA) based mass spectrometry profiles of 165 well characterized CLL patient samples (Fig. S1b, Validation1_DIA cohort), 18 CLL proteomes previously published (Validation2_Eagle, ^24^), 169 CLL transcriptomes (Validation3_RNA cohort), and two clinical validation cohorts of 620 untreated CLL patients (Validation4_untreated cohort) and 463 ibrutinib treated CLL patients (Validation5_ibrutinib).

To facilitate further analyses, we set up an open-access, user-friendly web application for others to explore and mine this comprehensive characterization of the relationship between the proteome, recurrent genetic aberrations, and the transcriptome of CLL, and consequences for drug response and clinical outcome (https://www.imbi.uni-heidelberg.de/dietrichlab/CLL_Proteomics/).

### Genotype-molecular phenotype connections with functional consequences

To explore genotype-phenotype relationships, we investigated associations between known recurrent genetic alterations of CLL ^9, 10^, mRNA expression, and protein abundance. We first analyzed protein abundance with respect to recurrent genetic aberrations of CLL and found that trisomy 12 had the strongest impact on differential protein abundance in comparison to other genetic lesions (Fig. 2a). First, we applied a very stringent FDR of 0.1 % and found 54 proteins significantly up- and 13 proteins downregulated in trisomy 12 positive CLL. IGHV mutation status (19 proteins, among them ZAP70) and *SF3B1* mutations (29 proteins) were also associated with multiple differentially abundant proteins. With a less stringent FDR of 5 % we found similar trends (Fig. S1c,d). Next, we analyzed gene expression with respect to recurrent genetic alterations of CLL (FDR 0.1 %, Fig. 2a; FDR 5 % Fig. S1e). Again, trisomy 12 (549 transcripts at FDR 0.1 %) and IGHV mutation status (205 transcripts at FDR 0.1 %) were associated with many significant gene expression changes. In contrast, *SF3B1* mutations were associated with fewer differentially expressed genes than with differentially abundant proteins (14 transcripts at FDR 0.1 %).

**Fig. 2:**
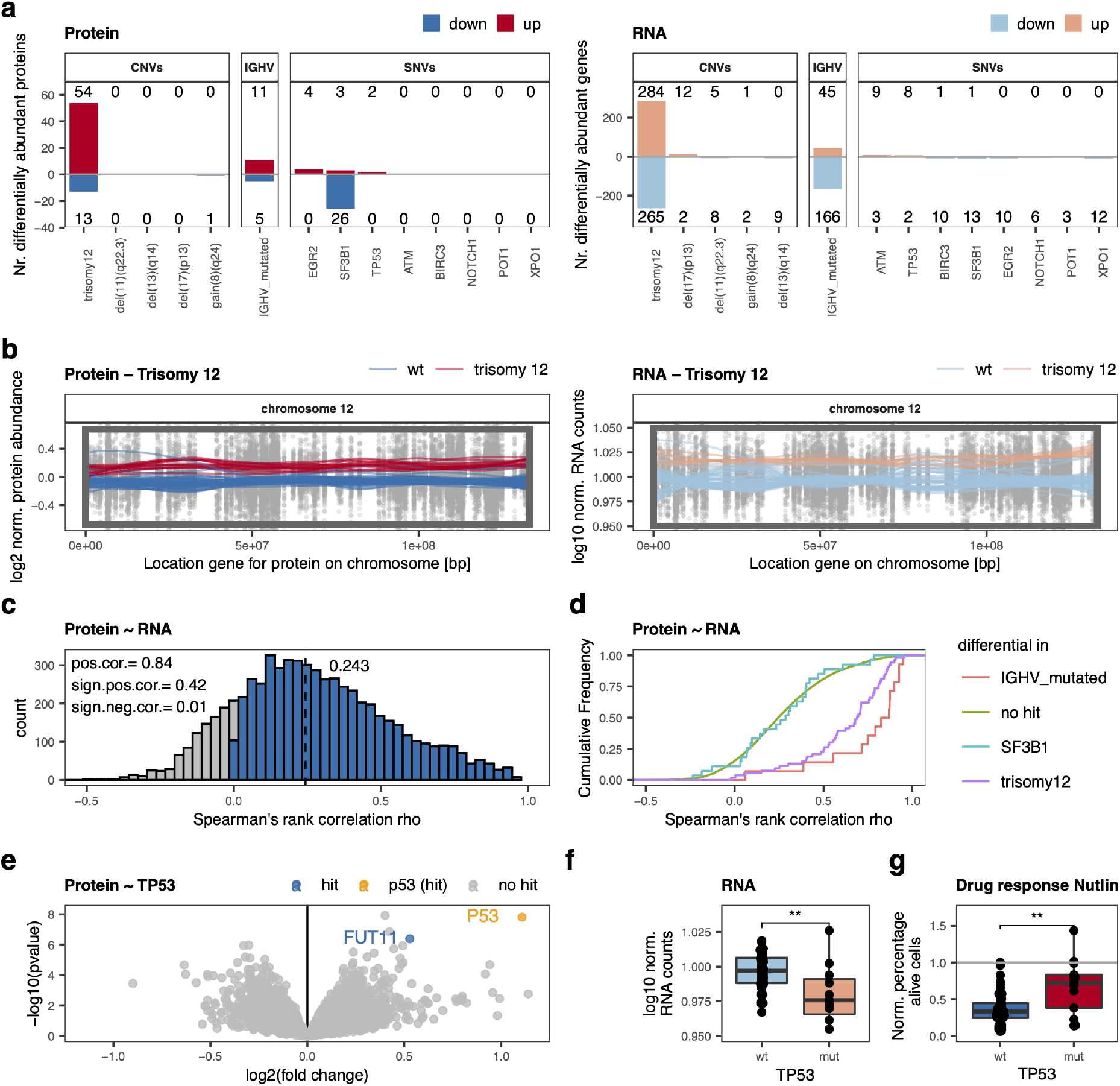
Interplay between genetic alterations, proteomics and transcriptomics. **a,** Number of significantly differentially abundant proteins (left; FDR < 0.1%; |log2FC| >0.5) and differentially expressed genes (right; FDR < 0.1%; |log2FC| >1.5) in relation to recurrent genetic alterations; red/positive numbers = upregulated, blue/negative numbers = downregulated. **b,** Levels of both proteins and transcripts from chromosome 12 were affected by trisomy 12. Normalized protein abundance (left panel) and gene expression levels (right panel) for chromosome 12 are shown. Points represent individual values for protein/gene - patient pairs. Lines are locally weighted scatterplot smoothed values for individual patients with (red) or without (blue) trisomy 12. The box is the region affected by trisomy 12. **c,** Distribution of Spearman’s rank correlations for protein-mRNA pairs. Negative correlations are shown in grey, positive correlations are shown in blue. **d,** Cumulative density distribution of protein-mRNA Spearman’s rank correlations for the proteins significantly differentially abundant in IGHV mutated (red), trisomy 12 (pink) or *SF3B1* mutated (blue) CLL in comparison to all other proteins without these associations (green). **e,** Volcano plot indicating differential proteins in *TP53* mutated in comparison to *TP53* wild type CLL; hit =adjusted p <0.001, |log2FC| >0.5. **f,** *TP53* transcript levels in *TP53* mutated (mut) and wild-type (wt) CLL samples; Wilcoxon signed-rank test p =0.005. **g,** Percentages, normalized to solvent control, of alive cells of *TP53* mutated (mut) and wild-type (wt) CLL samples treated *ex-vivo* with 9 µM nutlin 3a; Wilcoxon signed-rank test, p =0.003.

We further investigated if changes in protein abundance associated with trisomy 12 were related to gene dosage effects. While trisomy 12 increased the abundance of proteins located on chromosome 12 as expected (Fig. 2b), 63 % of the proteins identified as differentially abundant were expressed by genes located on other chromosomes. Similar gene dosage effects affecting mRNA and protein levels were also observed for other important structural aberrations of CLL, e.g. deletion of 13(q14), 11(q22-23), 17(p13), and gain of 8(q24) (Fig. S2a,b). As shown for other cancers, our data confirm in CLL that gene dosage effects translate into transcriptomic as well as proteomic changes ^14, 15, 17, 27^.

Even though alterations like trisomy 12 or presence of somatic hypermutations translated into changes in levels of both mRNA and protein, overall correlation between protein abundance and corresponding mRNA levels was low (Fig. 2c; median correlation= 0.243). This is comparable to gene ∼ protein correlations reported for other cancers, suggesting wide post-translational regulation of cancer proteomes ^14, 15, 17^. Out of all quantified genes and corresponding proteins, 42% showed significant, positive correlations (BH adjusted p <0.05). Differentially abundant proteins associated with trisomy 12 or IGHV mutation status exhibited higher mRNA ∼ protein correlations than unassociated proteins (Fig. 2d; median correlation trisomy 12 associated proteins= 0.69; IGHV associated proteins= 0.86). This was not the case for differentially abundant proteins associated with *SF3B1* mutations (Fig. 2d; median correlation= 0.29). These findings further demonstrate a direct link between protein abundance changes and gene expression changes in trisomy 12 and IGHV-mutated CLL, while the tumorigenic effect of *SF3B1* mutations is caused by post-transcriptional mechanisms.

Although *TP53*, *ATM,* and *XPO1* mutations were associated with relatively few differentially abundant proteins, we detected specific and biologically-relevant protein abundance changes. p53 was the most upregulated protein in *TP53*-mutated compared to wild-type samples (Fig. 2e). It is well-established that not only loss-of-function but also gain-of-function mutations in *TP53* can contribute to tumor progression ^28^. Our results are consistent with the finding that these tumors accumulate high levels of mutant p53, contributing to gain-of-function properties ^29^. In contrast, *TP53* transcripts were significantly downregulated in *TP53*-mutated CLL samples (Fig. 2f), indicating that post-transcriptional mechanisms are responsible for the accumulation of mutant p53 in CLL, as suggested for other blood cancers^29^. The *ex-vivo* drug response screen revealed that *TP53-*mutated CLL samples responded worse to chemotherapy and the MDM2 inhibitor nutlin 3a than *TP53* wild-type samples (Fig. 2g), as expected ^29^. This example illustrates how multi-omics profiling can be used to trace the effect of a somatic mutation on the transcriptome, the proteome, and finally on the functional consequences for drug response.

We further found that protein levels of the tumor suppressor ATM and the nuclear transport protein XPO1 were lower in mutated than in wild-type samples (Fig. S2c,d). Both associations were only observed on protein but not on transcript level (Fig. S2c,d).

Together, our data shows that the CLL proteome not only mirrors many changes that are present on mRNA level, but also uncovers biological relationships not apparent from the transcriptome.

### Integrative analysis reveals proteomics factor associated with outcome

We performed an unsupervised multi-omics factor analysis (MOFA ^30^) to obtain an integrative view of covariations across multiple datasets. MOFA revealed eleven latent factors (LF) explaining at least 1.5 % of variation each (Fig. 3a). Only LF1, LF2 and LF9 were significantly associated with time to next treatment (TTNT, Fig. 3b). LF1 and LF2 were active in all data layers (proteome, RNA and genetics) and were mainly driven by trisomy 12 and IGHV mutation status in the genetics data (Fig. 3c). The proteins loaded onto LF1 and LF2 were not only significantly associated with TTNT in the discovery cohort (Fig. 3d), but also showed a significant association with overall survival in our DIA proteomics validation cohort (Validation1_DIA; Fig. S3). Interestingly, LF9 was nearly exclusively active in the proteomics data and was apart from LF1 and LF2 the only factor significantly associated with clinical outcome. Hence, MOFA revealed that proteomics profiling uncovers clinically relevant biology that was not detected with the other data layers.

**Fig. 3:**
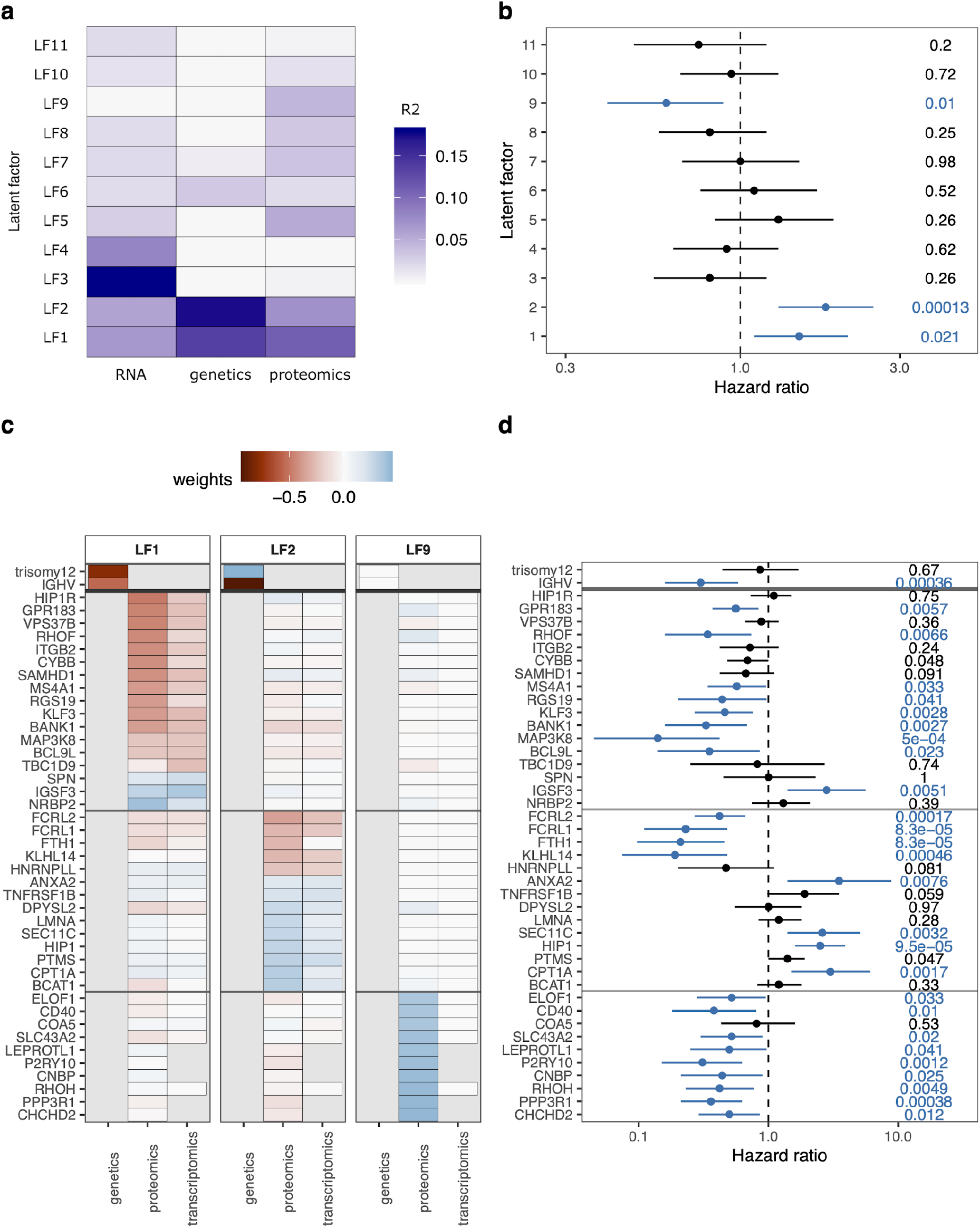
Multi-omics factor analysis (MOFA) of proteogenomics dataset. **a,** MOFA of proteome, transcriptome, and genome dataset identified 11 latent factors (LF) each explaining at least 1.5 % of variation. Explained variances per factor and dataset are color-coded. **b,** Hazard ratios from Cox regression of LFs with time to next treatment (TTNT). P-values are shown on the right. LF1, LF2 and LF9 were significantly (FDR <10 %, blue) associated with TTNT. Mean and 95 % confidence intervals are shown. **c,** Genes, transcripts, and proteins with the strongest weights loaded onto LF1, LF2 and LF9. Weights were scaled between genetics (divided by two), proteomics and transcriptomics (times ten) to achieve similar ranges. **d,** Hazard ratios from Cox regression for TTNT with genes and proteins with strong weights for LF1, LF2 and LF9. P-values are shown on the right. Significant associations (p <0.05) are colored in blue. Mean and 95 % confidence intervals are shown.

### Proteomics-based stratification of CLL identifies six distinct subgroups

We discovered associations of the proteomics layer and clinical outcome (TTNT), which could not be found on the transcript or on the genetic level. Therefore, we decided to explore the proteomics layer in depth. To describe similarities and differences between protein profiles of patients, we performed consensus clustering of the protein dataset. This revealed six proteomics groups (PG; Fig. 4a, S4a,b). T-distributed stochastic neighbor embedding (t-SNE) and principal component analysis supported the partition into these subgroups (Fig. S4c,d).

**Fig. 4:**
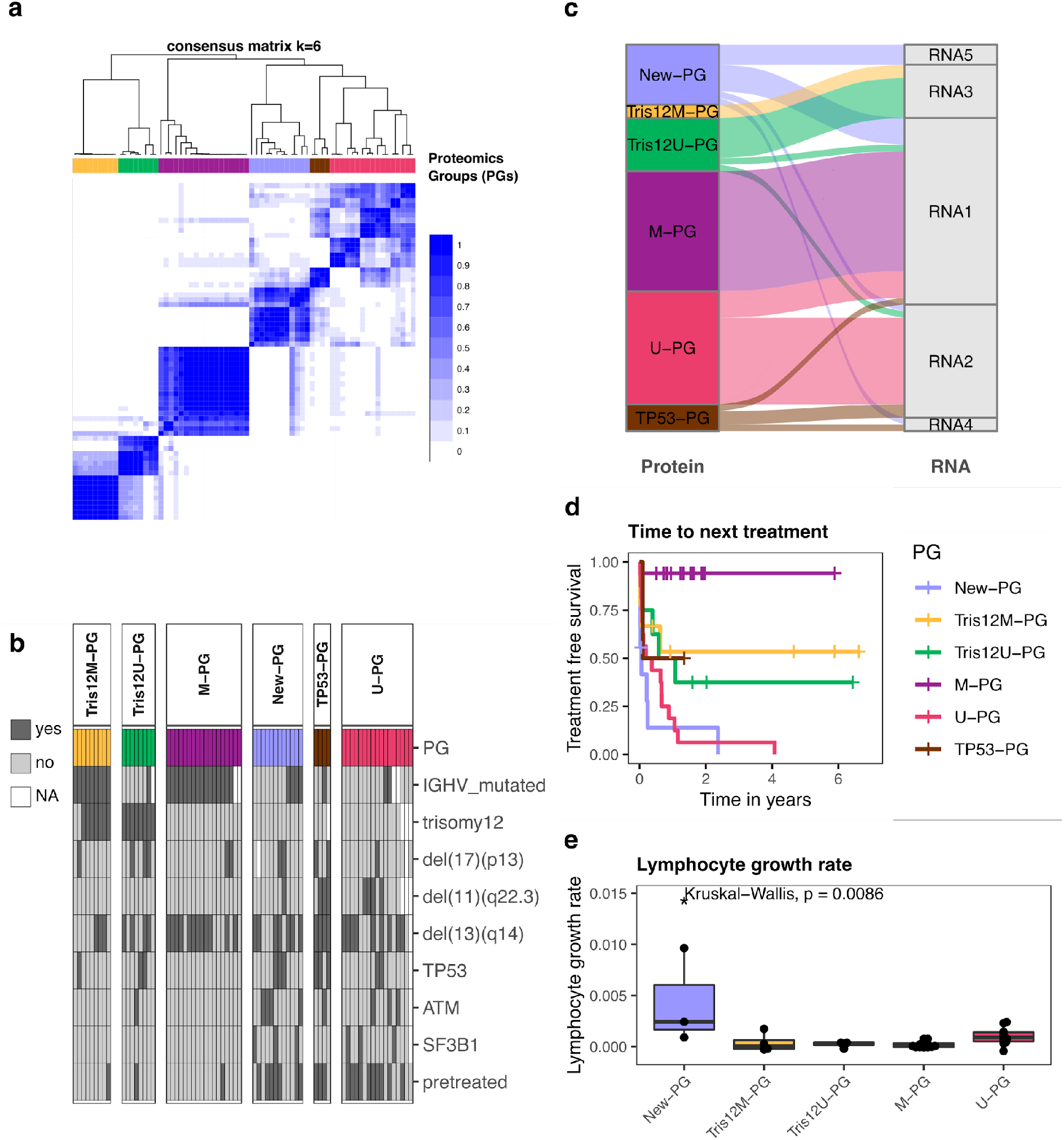
Consensus clustering of proteomics profiles partitioned CLL into six subgroups. **a,** Consensus clustering partitioned CLL samples into six proteomics groups (PGs). The color scale indicates which proportion of iterations patients clustered together. **b,** Annotation of PGs with genetic alterations and pretreatment status. **c,** Comparison of proteomic-based with transcriptomic-based consensus clustering. **d,** The PGs had significantly different times to next treatment (TTNT; log-rank test, p <0.0001). **e,** Lymphocyte growth rates for Tris12M-PG, Tris12U-PG, M-PG, U-PG, and New-PG. No data for TP53-PG was available.

Next, we analyzed if any genetic alterations were associated with the different PGs. We found that four PGs represented trisomy 12/M-CLL (Tris12M-PG, n =9), trisomy12/U-CLL (Tris12U-PG, n =8), M-CLL (M-PG, n =18), and U-CLL (U-PG, n =17, Fig. 4b; Fisher’s exact test, FDR <10 %). A fifth subgroup, TP53-PG, comprised only 4 patient samples, but three of the four samples harbored a *TP53* mutation (Fig. 4b; Fisher’s exact test, FDR < 10%). In accordance with the discovery of LF9, which was exclusively active in the proteomic layer, we additionally found a new proteomics based subgroup, which in contrast to all other subgroups did not show any association with known recurrent genetic alterations (New-PG). Only trisomy 12 was depleted in the new PG.

To relate these proteomics-based groups to the transcriptome, we performed consensus clustering of corresponding transcriptome data. The transcriptomic-based subgroups only partially overlapped with the proteomic groups (Fig. 4c): There was correspondence between RNA groups 1-3 and the proteomics groups Tris12U-PG, Tris12M-PG, M-PG and U-PG, but the new subgroup without a genetic annotation (New-PG) and TP53-PG (*TP53*) were split across multiple transcriptomic subgroups.

We further investigated drug response profiles of all PGs. Although effect sizes differed, the groups responded to the majority of CLL relevant drugs *ex-vivo*. Only the *TP53*-mutated group TP53-PG exerted poor overall response to many drugs including chemotherapeutic agents (Fig. S4e,f).

The proteomic-based grouping separated patients with different TTNT, demonstrating the clinical relevance of the subgroups (Fig. 4d). Although the new PG was not enriched for high risk factors such as mutated *TP53* and unmutated IGHV genes, it had the shortest TTNT (Fig. 4d) and the fastest *in-vivo* lymphocyte doubling time of all PGs, which indicated an increased proliferative capacity of these tumors (Kruskal-Wallis test, p =0.009, Fig. 4e). In contrast, M-PG (M-CLL, no trisomy 12) had a significantly longer TTNT than all other subgroups as expected.

Taken together, unsupervised clustering of relative protein abundances partitioned CLL patients into six biologically and clinically relevant groups, of which five could be explained by known genetic characteristics while a new subgroup with poor outcome was only identifiable based on proteomics.

### Dysregulated cellular processes of the new poor outcome proteomics group ASB-CLL

The new poor outcome subgroup comprised approximately 20% of our discovery cohort and could only be identified at the protein level. To describe which pathways and processes were altered in this group, we analyzed differential protein abundances between the new and other PGs. Enrichment analysis identified BcR signaling proteins such as BTK, PLCG2 and PIK3CD among the most down regulated proteins in the New-PG (Fig. 5a, S5a,b, Data SI2). Surprisingly, downregulation of these central BcR signaling components in the New-PG was independent of the IGHV mutation status, despite the latter being an important surrogate for BcR activity in CLL ^31^. We therefore measured the *ex-vivo* response of the samples to ibrutinib in a co-culture model of stromal cells and CLL cells, which suggested that the New-PG responded less to the BcR inhibitor ibrutinib (Fig. S5c, Ibrutinib: 40 nM, Wilcoxon signed-rank test p =0.01).

**Fig. 5:**
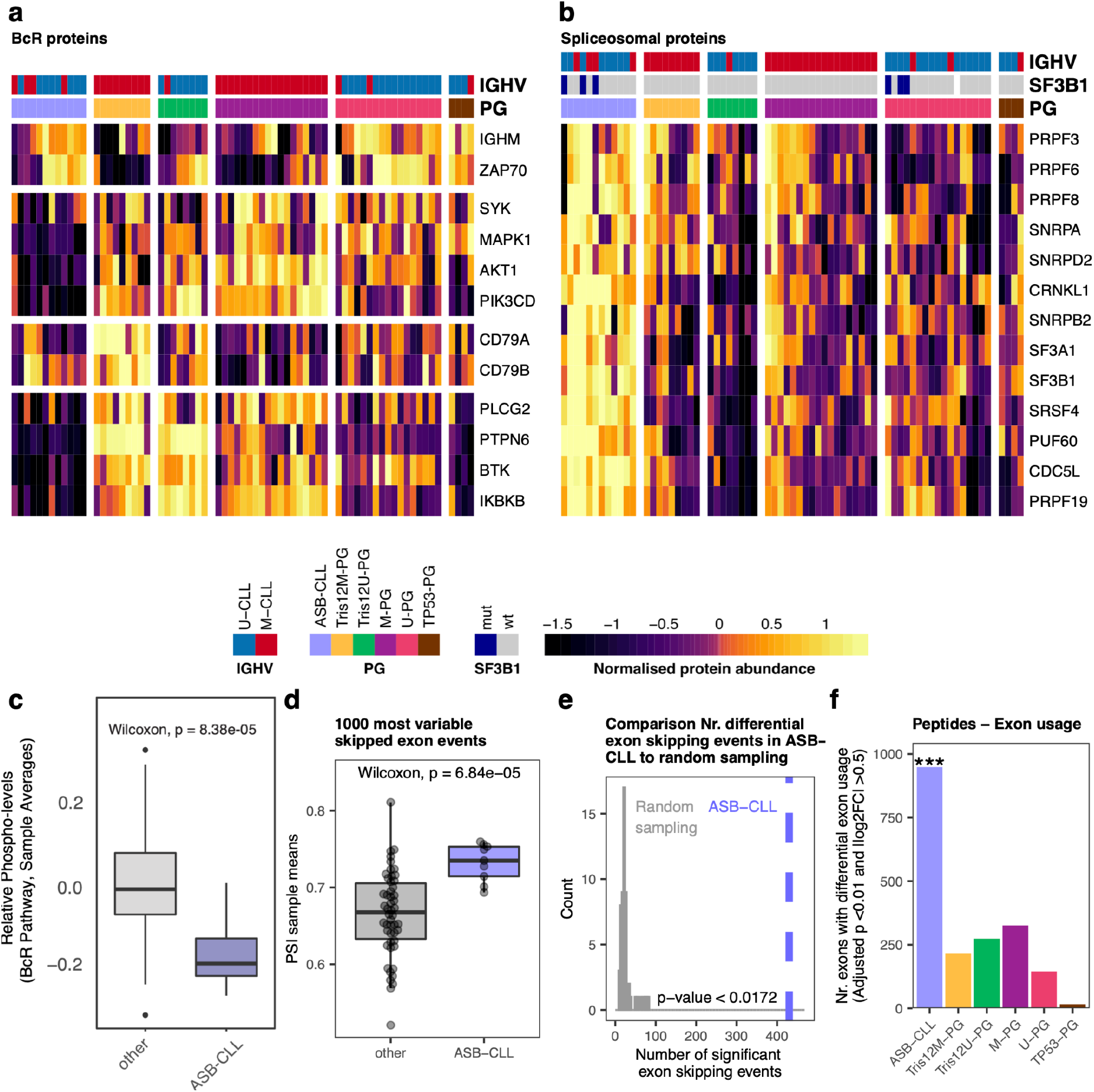
Characterization of the new proteomics group ASB-CLL. **a,** Heatmaps of scaled log2 protein abundances for selected BcR proteins and **b,** spliceosome components. Patients were grouped according to PG and the proteins were clustered hierarchically. Based on these profiles the group “New-PG” was renamed to “ASB-CLL” (**A**ltered **S**pliceosome, low **B**cR signaling proteins **CLL**) **c,** Mean percent spliced-in (PSI) value per patient calculated from the 1000 most variable exon skipping events across all patients of the discovery cohort. ASB-CLL patients were compared to all other patients. **d,** Number of significant differential exon skipping events between ASB-CLL and all other groups for the actual PG assignment (blue line) and random PG permutations (grey distribution)**. e,** Peptide-based differential exon usage per PG (FDR 1 %, |logFC| >0.5).

The New-PG was further characterized by upregulation of enzymes involved in the degradation of branched chain amino acids (BCAA; Fig. S5d,e) and downregulation of proteasomal proteins (Fig. S5f,g). Most importantly, proteins associated with the spliceosome were most significantly upregulated in the New-PG (Fig. 5b, S5h,i, Data SI2). We therefore named this new group ASB-CLL (**A**ltered **S**pliceosome, low **B**cR signaling proteins **CLL**). We and others observed very low correlation between protein- and corresponding mRNA levels of components of the spliceosome ^14, 17^ (median rho = -0.015; Fig. S6a), which might explain why ASB-CLL was not detectable at the transcriptome level.

Splice factor mutations (e.g. *SF3B1* mutations) are recurrent in CLL ^10^; however, mutations of *SF3B1* were neither enriched nor depleted in ASB-CLL (BH adjusted p = 0.39). SF3B1 protein levels were not different between *SF3B1* mutated and wild-type CLL cases, but were significantly higher in ASB-CLL than in other subgroups (Fig. S6c). It has recently been reported that the main tumorigenic effect of *SF3B1* mutations is mediated by inclusion of a poison-exon in the tumor suppressor gene *BRD9* followed by downregulation of BRD9 ^32^. We could confirm mis-splicing (Fig. S6d) and downregulation (Fig. S6e) of BRD9 in *SF3B1*-mutated cancers, but neither mis-splicing nor downregulation of BRD9 was detected for ASB-CLL (Fig. S6f,g). For ten out of the twelve ASB-CLL patients we further performed whole-exome sequencing of tumor and matched normal controls, to explore whether any additional somatic mutations in spliceosomal proteins could be found for this subgroup. Except for a *DDX5* mutation in a *SF3B1* mutated patient we could not detect any other splice factor mutations (Fig. S6h). Together these results suggest that the increased spliceosomal protein abundance detected in ASB-CLL is independent of mutations in the spliceosome.

To further characterize if upregulation of the spliceosome in ASB-CLL had any functional consequences, we analyzed alternative splicing on mRNA and peptide level for this group. We started with an unbiased approach and annotated all possible alternative splicing events across all patients using the transcriptome data and rMATS ^33^. Within each category (skipped exons, 3’ and 5’ alternative splice site usage, retained introns, mutually exclusive exons) we calculated the per-patient mean percent spliced-in (PSI) values from the 1000 alternative splicing events that showed the highest variability across all patients (Fig. S6i). Comparisons between ASB-CLL and all other subgroups revealed a distinct exon usage profile for ASB-CLL, with fewer skipped exons, and more 3’ and 5’ alternative splice site usage (Fig. 5c, S6i). In total, we detected 427 exon skipping events with a significantly distinct alternative splicing pattern of ASB-CLL versus all other proteomics subgroups (FDR <1 %, absolute difference of groupwise mean PSI values >0.1). These exon skipping events were significantly more frequent in ASB-CLL than expected by chance (permutation test, p <0.0172, Fig. 5d). Significantly different exon usage by ASB-CLL was also detected on peptide level (Fig. 5e, Data SI3).

Thus, ASB-CLL was characterized by low abundance of BcR pathway components, lower phosphorylation levels of BcR components and altered spliceosome function.

### Validation of ASB-CLL in independent cohorts

We confirmed the existence and prognostic relevance of the ASB-CLL PG in additional cohorts of CLL patients (Fig. 6a): We validated the proteomics signature associated with ASB-CLL by performing proteomics using data independent acquisition (DIA) based mass spectrometry analysis of 165 patients (Validation1_DIA). We also confirmed the proteomics signature in an independent, previously published cohort (Validation2_Eagle)^24^. We further validated the splicing signature characteristic for ASB-CLL in an independent cohort of 169 CLL patients (Validation3_RNA cohort).

**Fig. 6:**
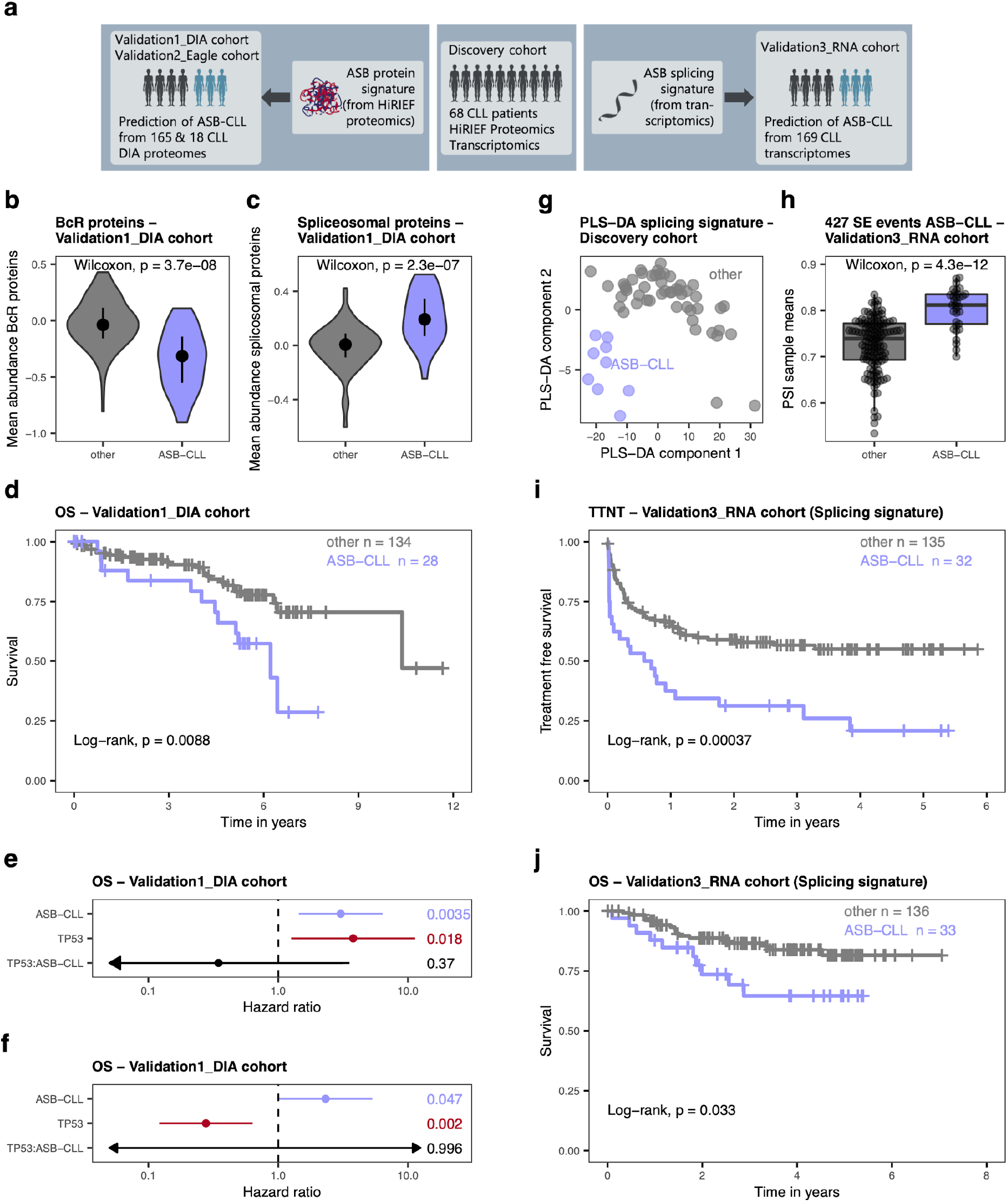
Validation of the new proteomics group ASB-CLL. **a,** Strategy for validating the existence of ASB-CLL in independent cohorts. A classifier was trained from the proteomics dataset of the discovery cohort and applied to a DIA proteomics validation dataset comprising 165 CLL patients (Validation1_DIA cohort; left part of figure) and 18 CLL patients from a published cohort (Validation2_Eagle)^24^. Using the transcriptomics dataset of the discovery cohort, a classifier was trained on the splicing signature of ASB-CLL and applied to a validation dataset comprising the transcriptomes of 169 CLL patients (Validation3_RNA cohort). **b,** Violin plot of BcR protein abundances comparing the subgroup identified as ASB-CLL in the Validation1_DIA dataset to all other patients. **c,** Violin plot of spliceosomal protein abundances comparing the subgroup identified as ASB-CLL in the Validation1_DIA dataset to all other patients. **d,** Overall survival (OS) of Validation1_DIA cohort, divided into ASB-CLL and all other patients. **e,** Hazard ratios from Cox regression of OS in a model including *TP53* aberrations (del(17)(p13) or *TP53* mutations), ASB-CLL, and their interaction (TP53:ASB-CLL). P-values are shown on the right. Both *TP53* aberrations as well as ASB-CLL independently predicted OS, while the interaction of the two factors did not significantly contribute. Mean and 95 % confidence intervals are shown. Arrows indicate that the confidence interval extends beyond the shown length. **f,** Hazard ratios from Cox regression of OS in a model including IGHV mutation status, ASB-CLL, and their interaction (IGHV:ASB-CLL). P-values are shown on the right. Both IGHV mutational status and ASB-CLL independently predicted OS, while the interaction of the two factors did not significantly contribute. Mean and 95 % confidence intervals are shown. Arrows indicate that the confidence interval extends beyond the shown length. **g,** Patient samples of the discovery cohort plotted in the plane of the first two components computed by the partial least squares discriminant analysis (PLS-DA) of the discovery cohort. A clear separation of ASB-CLL samples is visible. The predictive performance of the classifier was assessed using leave-one-out cross-validation with the area under the ROC curve being estimated as 0.97. **h,** Mean PSI value per patient of the Validation3_RNA cohort based on the 427 significant differential exon skipping events identified in the discovery cohort. **i,** Time to next treatment (TTNT) of patients from the Validation3_RNA cohort, classified as ASB-CLL or other based on the characteristic splicing signature of ASB-CLL. ASB-CLL patients had significantly faster disease progression (log-rank test, p =0.00037). **j,** OS of patients from the Validation3_RNA cohort, classified as ASB-CLL or other based on the characteristic splicing signature of ASB-CLL. ASB-CLL patients had significantly shorter overall survival (log-rank test, p =0.033).

To robustly detect ASB-CLL in DIA proteomics, we trained a k-top scoring pairs (kTSP) classifier on all BcR and spliceosomal proteins in the HiRIEF dataset. We applied the classifier to the Validation1_DIA cohort and identified 28 ASB-CLL patients (10 untreated, 15 pretreated, 3 unknown), corresponding to 17 % of patients. This proportion of ASB-CLL was similar to the training dataset. As expected, BcR proteins were down- and spliceosomal proteins were upregulated in ASB-CLL (Fig. 6b,c; Wilcoxon signed-rank test p =p <3.7×10^-8^, p <2.3×10^-7^). Additionally, ASB-CLL was characterized by upregulation of BCAA proteins and downregulation of proteasomal proteins, which is in accordance with the findings observed in the discovery cohort (Fig. S7a,b; Wilcoxon signed-rank test p =0.002 and p <2.2×10^-6^). Most importantly, ASB-CLL patients had significantly worse overall survival (log-rank test, p <0.0088; Fig. 6d) and this was independent of the risk associated with *TP53* aberrations (del(17)(p13) or *TP53* mutations) and unmutated IGHV genes (Fig. 6e,f). The poor overall survival of ASB-CLL was confirmed in the subgroup of samples obtained from treatment naive patients (log-rank test, p =0.034; Fig. S7c). Application of the kTSP classifier to an independently published cohort by Eagle and colleagues (Validation2_Eagle) ^24^ comprising nine U-CLL and nine M-CLL patients identified a subgroup of five patients with a genetic and proteomics profile similar to ASB-CLL (Fig S7d-g).

Additionally, we used an independent cohort of patients (Validation3_RNA) for which paired-end RNA-sequencing was performed to validate the splicing signature detected for ASB-CLL. A partial least squares discriminant analysis (PLS-DA) classifier was trained on the discovery cohort samples using the previously identified 427 significantly differential exon skipping events as predictors (Fig. 6g). Predictive performance of the PLS-DA model was assessed by leave-one-out cross-validation, which gave an estimate of 0.97 for the area under the classifier’s ROC curve. The PLS-DA model was then applied to the Validation3_RNA cohort to classify each of its 169 patients as either belonging to or not belonging to ASB-CLL. As observed in the training cohort, significantly larger mean PSI values of exon skipping events were detected for patients identified as belonging to ASB-CLL than for other patient groups (two-sided Wilcoxon rank sum test, p = 4.3e-12, Fig. 6h). The ASB-CLL samples in the Validation3_RNA cohort exhibited the same splicing pattern as in the discovery cohort and also demonstrated a significantly shorter TTNT (log-rank test, p <0.00037, Fig. 6i) and overall survival (log-rank test, p <0.033, Fig. 6j).

### Cell of origin influences the effect of trisomy 12 on the proteome and clinical outcome

Plotting of the latent factors 1 and 2 from our MOFA resulted in the separation of four distinct groups (Tris12M-PG, Tris12U-PG, M-PG, U-PG), which were strongly associated with IGHV mutation status and trisomy 12 (Fig. 7a,b). Initially, we aimed to understand the functional consequences of this interaction for the proteome and subsequently used the association of these groups with trisomy 12 and IGHV mutation status to investigate survival differences between these subgroups in three independent data sets (Fig. 7a).

**Fig. 7:**
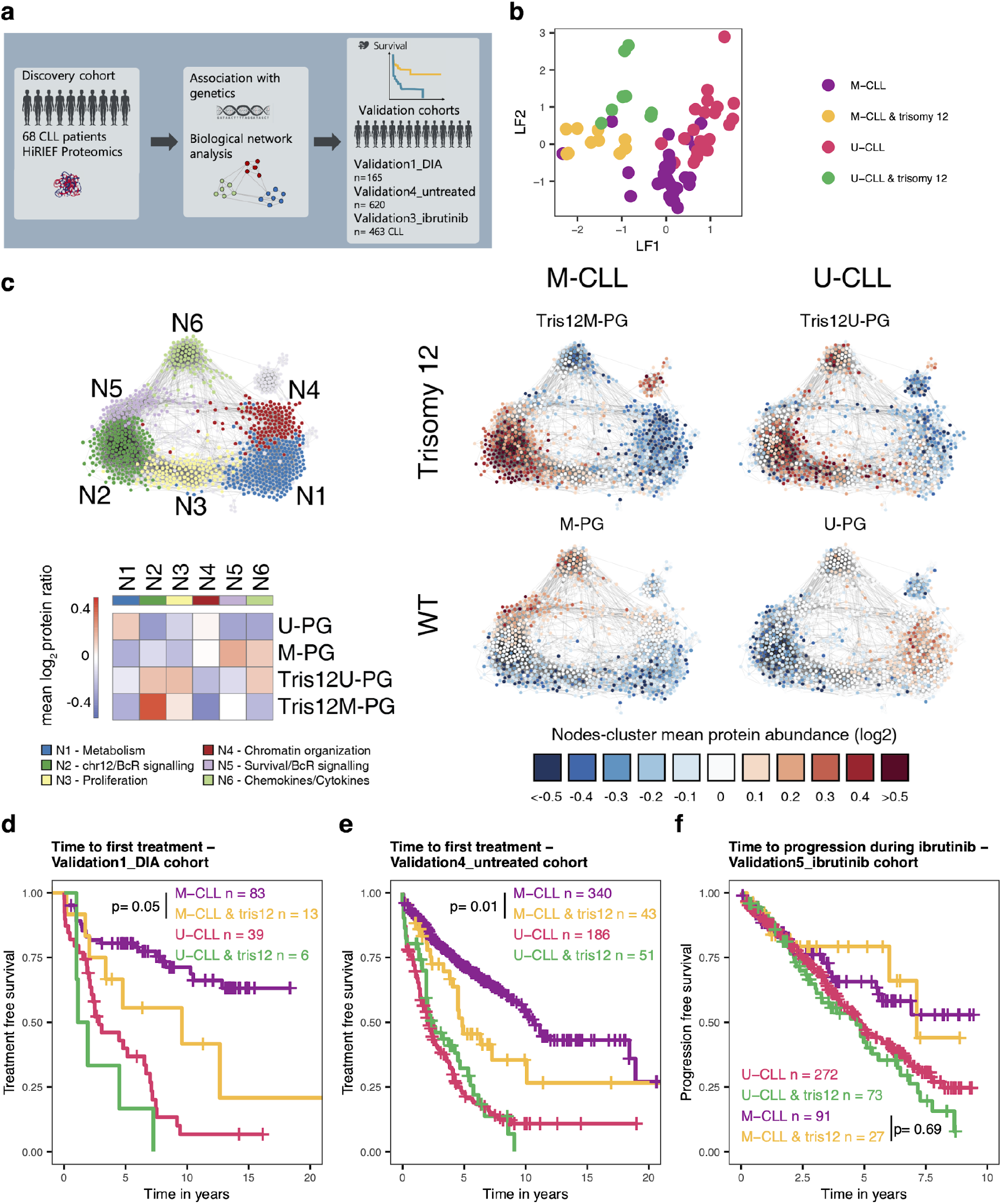
Relevance of the cell of origin for the influence of trisomy 12 on the CLL proteome and clinical outcome. **a,** Validation strategy for Tris12M-PG, Tris12U-PG, M-PG, and U-PG. **b,** Plot of loadings on LF1 and LF2 of individual patients. LF1 and LF2 could classify patients into 4 groups according to IGHV status and trisomy 12. **c,** CLL protein correlation network analysis of HiRIEF dataset based on 1047 high-variance proteins. Protein groups were defined and color-coded based on modularity clustering (N1-N6) and enrichments detailed in Fig. S8. A Heatmap of log2 mean relative protein abundances for all PGs for each modularity cluster (N1-N6) and node-cluster mean protein abundances for Tris12M-PG, Tris12U-PG, M-PG, and U-PG are shown. **d,** Time to first treatment of patients from Validation1_DIA cohort, stratified into groups by IGHV mutation status and trisomy 12 (tris12). Trisomy 12 M-CLL patients had significantly faster disease progression than M-CLL patients without trisomy 12 (log-rank test, p= 0.05). **e,** Time to first treatment of untreated patients from Validation4_untreated cohort, stratified into groups by IGHV mutation status and trisomy 12 (tris12). Trisomy 12 M-CLL patients had significantly faster disease progression than M-CLL patients without trisomy 12 (log-rank test, p= 0.01). **f,** Time to progression of patients uniformly treated with ibrutinib (Validation5_ibrutinib), stratified into groups by IGHV mutation status and trisomy 12 (tris12). Trisomy 12 M-CLL patients did not have significantly faster disease progression than M-CLL patients without trisomy 12 (log-rank test, p= 0.69).

To understand the global influence of IGHV status and trisomy 12 on the proteome, we performed a correlation network analysis^15^ of all samples in the discovery cohort independently of the subgroups to identify the most important biological modules of the CLL proteome (Fig. 7c). This revealed six biological modules (N1-N6) that were affected in all screened CLL patient samples. Next, we performed enrichment analysis for these modules and quantified the importance of each module in each PG (Fig. 7c, S8a). All trends were confirmed by the networks with relative protein abundances from the DIA proteomics data set of the Validation1_DIA cohort (n=165 CLL patients, Fig. S8b). As expected, the two trisomy 12 subgroups Tris12M-PG and Tris12U-PG displayed elevated levels of proteins located on chromosome 12 and proteins involved in BcR signaling (N2). We further assessed functional consequences of this upregulation for *ex-vivo* drug response and found that Tris12M-PG and Tris12U-PG were significantly more sensitive to BcR pathway inhibitors (e.g. ibrutinib, idelalisib) measured in an *ex-vivo* co-culture model of stroma- and CLL cells (Fig. S8c).

Despite these similarities, there were also biologically relevant differences between Tris12M-PG and Tris12U-PG depending on the IGHV mutation background. The module containing proteins involved in chemokine and cytokine signaling (N6), was significantly more abundant in Tris12U-PG. By adjusting for U-CLL/M-CLL differences in cases without trisomy 12 we noted that U-CLL cases, in the context of trisomy 12, exhibited increased levels of MAPK signaling proteins while M-CLL cases had increased levels of cell adhesion molecules (Fig. S8d). We could also identify differences in the BcR signaling pathway between M-CLL and U-CLL, specific to the context of trisomy 12; with U-CLL, trisomy 12 cases exhibiting a further increase of the intracellular components of the pathway (e.g. NFKB1, MAPK13), and M-CLL, trisomy 12 exhibiting an increase of membrane bound components (e.g. CD21, CD79A/B, Fig. S8e).

Occurrence of trisomy 12 in M-CLL was associated with low abundance of proteins in N5, which have been shown to be related to cell survival ^34^ (Fig. S8a). In contrast, occurrence of trisomy 12 in U-CLL did not change protein levels in this module.

We further sought to investigate the impact of trisomy 12 on the natural course of M-CLL and U-CLL independently of any treatment context using time to first treatment (TTFT) as the endpoint. Trisomy 12 did not alter TTFT in U-CLL patients, but significantly decreased TTFT within the M-CLL patients (Validation1_DIA; Fig. 7d; log-rank test, p= 0.05). This could be confirmed in a cohort of 620 untreated CLL patients (Validation4_untreated; Fig. 7e; log-rank test, p= 0.01). The faster disease progression of trisomy 12 M-CLL cases compared to M-CLL without trisomy 12 was also observed in the subset of 437 cases devoid of 11(q22-23) deletion and *TP53* disruption (p=0.0016), as well as in a Rai 0 cohort of 409 cases (p=0.0021; data not shown).

Interestingly, a faster progression of Tris12M-PG patients was not observed in a cohort of 463 CLL patients uniformly treated with ibrutinib (Validation5_ibrutinib; Fig. 7f; log-rank test, p= 0.69). However, patients with trisomy 12 had a better response to ibrutinib *in-vivo*^35^ and *ex-vivo* (Fig. S8c). Therefore, we hypothesized that the better response rate of trisomy 12 patients to BcR inhibitors balances out the faster progression observed in untreated Tris12M-PG patients.

These proteogenomics findings exemplify how the influence of a genetic alteration on the proteome varies depending on the cellular background in which it occurs, and how this interaction could translate into different clinical outcomes.

## Discussion

Cancer proteogenomic studies integrate proteomic and genomic data to gain understanding of the impact of genetic alterations on the proteome, both on particular proteins and proteome-wide effects. As proteins are considered the main effectors of many cellular processes this approach has recently been used to improve the understanding of the drivers of selected cancer entities ^14–20^.

We applied in-depth mass spectrometry based proteomics, using HiRIEF fractionation and data-dependent acquisition, to thoroughly characterize the disease biology of CLL, followed by data-independent acquisition proteomics to validate previously unknown cancer subgroups. Our approach demonstrates how these proteomics methods can complement each other to combine the benefits of deep and high-throughput proteomic profiling. This is especially relevant for association studies with clinical outcome, which require large cohort sizes to provide sufficient statistical power.

This study was conducted using cryopreserved cells that may vary from fresh, never frozen cells. For future development of clinical proteomics workflows, the impact of sample handling will need to be addressed further in order to ensure robust implementation. However, in this study, we demonstrated that both data-independent acquisition and data-dependent acquisition LC-MS/MS can be used to stratify patients based on proteome subtypes using cryopreserved samples.

Our work comprehensively illustrates that proteomic profiling of cancer cells has the ability to improve the understanding of clinically relevant disease biology beyond transcriptomic and genomic profiling. We linked proteomic and transcriptomic profiles to important recurrent genetic aberrations in CLL. This revealed, for instance, that RNA and protein levels were disconnected for important CLL drivers like *TP53* and *XPO1* mutations, while the biologically meaningful information was primarily contained in the proteome.

We utilized a method to simultaneously detect covariances across multiple omics layers and discovered a proteomic profile of CLL without a corresponding genetic or transcriptomic profile, which was strongly associated with clinical outcome. So far, few studies have managed to associate proteomic profiles with clinical outcome ^21, 36, 37^. Our results demonstrate that this poor prognostic signature is stable across patient cohorts and can be determined by different techniques. These findings highlight that proteomic profiling of cancer can improve molecular stratification of cancer patients.

This new biological axis of CLL discovered by proteomics could be identified in approximately 20 % of CLL patients. This subgroup was named ASB-CLL and was characterized by a high proliferative capacity and poor overall survival. We found it to be independent of conventional risk factors such as *TP53* aberrations ^38^ and IGHV mutation status ^5^. These findings illustrate the added value of proteomics for a better understanding of the clinically relevant disease biology of CLL. Our example using CLL as a model disease entity shows that clinical implementation of mass spectrometry based screening, using the data from this study as a template, can aid not only in identifying patients at particular risk for aggressive disease but also in tailoring treatment and clinical monitoring accordingly.

ASB-CLL was characterized by low abundances of major BcR signaling proteins and high abundances of components of the spliceosome. This directly translated into an altered splicing pattern with increased exon-inclusion and was independent of mutations in *SF3B1*^32^. These results describe for the first time that deregulated splicing in CLL occurs as a result of splicing factor upregulation rather than splicing factor mutation. Future mechanistic studies need to clarify how spliceosomal protein abundance is regulated in this subgroup of CLL.

Trisomy 12 and IGHV status explained four of six subgroups with distinct proteomic profiles, suggesting that the interaction of these genetic factors drives the biological differences between these subgroups. While the IGHV mutation status is known to influence the strength of BcR signaling and response to BcR inhibitors ^39^, the biology of trisomy 12 is still insufficiently understood. Our results improve the current understanding of trisomy 12 and demonstrate that trisomy 12 significantly increases the abundance of BcR signaling proteins and BcR activity. In accordance with a previous study we showed that treatment naive M-CLL patients without trisomy 12 had an indolent disease course while M-CLL patients with trisomy 12 had a faster disease progression^40^. The accelerated progression dynamics of trisomy 12 positive M-CLL was not observed in ibrutinib treated CLL patients, which implies that inhibition of trisomy 12 mediated BcR activation compensated for this disadvantage. Future clinical studies should explore how to best exploit this new insight and whether trisomy 12 M-CLL patients benefit from a BcR inhibitor treatment algorithm designed for U-CLL. These results exemplify the importance of genetic marker combinatorics for gene expression, protein abundance, drug response and disease progression.

Our integrative multi-omics analysis of CLL provides the first comprehensive overview of the interplay between genetic alterations, the transcriptome, and the proteome, along with functional consequences for drug response and clinical outcome. The detailed analysis of our dataset improves our understanding of the biological heterogeneity of CLL and provides molecular phenotype-based subtypes that will improve patient stratification and personalized treatments. Through our web application we provide this comprehensive dataset as a valuable and easily accessible resource to the research community (https://www.imbi.uni-heidelberg.de/dietrichlab/CLL_Proteomics/).

## Methods

### Experimental procedures

#### Study design

The purpose of this study was to investigate the relationships between the proteome, transcriptome, and genetic aberrations to clinical parameters (notably TTNT) and drug sensitivity in CLL. The proteome of CLL cells from 68 patient samples was analyzed by HiRIEF-LC-MS/MS, the transcriptome of 59 patients by RNA-sequencing and mutations of 68 samples by DNA panel sequencing. To minimize the introduction of biases all of these datasets were acquired from the same aliquot of cells. Drug sensitivity was assessed by microscopy based *ex-vivo* drug screening and clinical parameters (TTNT) were determined from patient records.

The results obtained from this discovery cohort were validated in five independent CLL cohorts.

#### Discovery cohort: Patient samples

Written consent was obtained from patients according to the declaration of Helsinki. Leukemia cells were isolated from blood using Ficoll density gradient centrifugation. Cells were viably frozen and kept on liquid nitrogen until use. 45 CLL patients were untreated at sample collection and 23 had received prior treatment with immuno-chemotherapy (n=17) or with novel drugs (n=6).

#### Discovery cohort: Sample preparation for proteomics, RNA, and panel sequencing

Cells were thawed, allowed to recover in RPMI medium (Thermo Fisher Scientific) containing 10 % human serum (Sigma Aldrich) for 3h and filtered through a 40 µm cell strainer for removal of dead cells from the thawing process. Viability was assessed with Trypan Blue. Only samples with a viability above 90% were included. Tumor cells were collected by Magnetic-activated cell sorting (MACS) using CD19 beads (Miltenyi Biotec). Samples were split into aliquots for proteomics analysis (1×10^7^ cells), RNA sequencing (5×10^6^ -1×10^7^ cells), and panel sequencing (5×10^6^ cells). Pellets for proteomics analysis were washed twice with PBS and snap frozen in liquid nitrogen. DNA for panel sequencing was extracted using the DNeasy Blood & Tissue Kit (Qiagen). For RNA sequencing RNA was isolated using QIAzol Lysis Reagent (Qiagen), QIAshredder (Qiagen), and the RNeasy Mini Kit (Qiagen).

#### IGHV mutation status analysis

RNA was isolated and cDNA synthesized using High-capacity cDNA Reverse Transcription Kit (Thermo Fisher Scientific). PCR reactions and analyses were performed as previously described with minor modifications ^41^ (see supplementary methods for additional details).

#### DNA copy-number variants

DNA copy numbers were assessed using Illumina CytoSNP-12 and HumanOmni2.5-8 microarrays and read out using the iScan array scanner. Fluorescence in situ hybridization (FISH) analysis was performed for del(11)(q22.3), del(17)(p13), del(13)(q14), trisomy 12, gain(8)(q24), and gain(14)(q32). Only alterations present and absent in at least three patients were considered for analyses.

#### Discovery cohort: HiRIEF proteomics

Cell pellets were lysed by 4 % SDS lysis buffer and prepared for mass spectrometry analysis using a modified version of the SP3 protein clean up and digestion protocol ^42^. Peptides were labelled with TMT10-plex reagent according to the manufacturer’s protocol (Thermo Fisher Scientific) and separated by immobilized pH gradient - isoelectric focusing (IPG-IEF) on 3–10 strips as described previously ^25^. Extracted peptide fractions from the IPG-IEF were separated using an online 3000 RSLCnano system coupled to a Thermo Fisher Scientific Q Exactive-HF. MSGF+ and Percolator in the Galaxy platform were used to match MS spectra to the Ensembl92 human protein database ^43^. For acetylation and methylation variable modifications, we performed two separate searches; one where acetylation was allowed on lysines and one where monomethylation was allowed on lysines and arginines (see supplementary methods for additional details). The mass spectrometry proteomics data have been deposited to the ProteomeXchange Consortium via the PRIDE partner repository with the dataset identifier PXD017453.

#### Discovery cohort: Panel sequencing

For gene mutation analysis we designed a customized Illumina™ TruSeq Custom Amplicon (TSCA) panel with two independent primer sets for redundant coverage ^26^. Mutations in the genes A*TM, BIRC3, EGR2, FBXW7, MYD88, NFKBIE, POT1, TP53, BRAF, NOTCH1, RPS15, SF3B1,* and *XPO1* were covered. Library preparation was performed using TruSeq Custom Amplicon Assay Kit v1.5 and sequenced on an Illumina MiSeq flowcell. For analysis a custom bioinformatics pipeline was used. See supplementary methods for additional details.

#### Discovery cohort: mRNA Sequencing

Stranded mRNA sequencing, using a TruSeq Stranded Total RNA Library Preparation Kit, was performed on a Illumina NextSeq 500. Reads were aligned to GRCh 37.75/hg 19 using STAR (v2.6.0c; ^44^) and counted with htseq-count ^45^. Library size normalization, variance stabilizing transformation and differential expression calling were performed using DESeq2^46^.

#### Discovery cohort: *Ex-vivo* drug sensitivity screen

Drug response profiles were obtained for 68 leukemia samples and 43 drugs (Selleckchem) in 3 concentrations (Data SI4). Cells were thawed and seeded in DMEM medium (Thermo Fisher Scientific) containing 10 % human serum (Sigma Aldrich), 1 % penicillin/streptomycin (Thermo Fisher Scientific) and 1 % glutamine (Thermo Fisher Scientific) at 2×10^4^ cells/well into a CellCarrier-384 Ultra Microplate (Perkin Elmer). Cells were incubated at 37 °C in a humidified atmosphere and 10 % CO_2_ for 3 days.

Cells were stained with 4 µg/ml Hoechst 33342 (Thermo Fisher Scientific). Images were taken using an Opera Phenix High Content Screening System (Perkin Elmer) and processed with Harmony (Perkin Elmer). Cells were segmented and the nuclear area was calculated. Based on a threshold of 23.8 µM^2^ nuclear area cells were classified into alive and dead. Percentage of alive cells was calculated and normalized by dividing through the mean percentage of alive cells across all solvent (DMSO) controls.

#### Validation1_DIA cohort: DIA proteomics

For DIA based proteomics cell pellets were digested and cleaned as described above. Unlabeled peptides from individual samples were separated using an online 3000 RSLCnano system coupled to a Thermo Fisher Scientific Q Exactive-HF. Data independent acquisition (DIA) was employed using a variable window strategy. Spectronaut was used to analyze the spectral files using the Direct-DIA option and files were searched against the ENSEMBL database (see supplementary methods for additional details). In total 203 samples were analyzed by DIA, 36 from the original discovery cohort and 167 validation samples, The mass spectrometry proteomics data have been deposited to the ProteomeXchange Consortium via the PRIDE partner repository with the dataset identifier PXD024544.

#### Validation2_Eagle cohort

For an additional validation of the existence of ASB-CLL in an external cohort, we took advantage of the cohort of 18 CLL patient samples published by Eagle et al. ^24^. Presence of trisomy 12 was estimated by calculating the mean abundance of proteins located on chromosome 12 and defining the 20% of patients with highest abundances as harboring trisomy 12.

#### Validation3_RNA cohort: RNA Sequencing

To validate the splicing signature of ASB-CLL we took advantage of a cohort of 169 CLL patients for who paired end sequencing, using an Illumina TruSeq RNA sample preparation kit v2, was performed on an Illumina HiSeq2000 with 300 bp insert size ^47^. No overlap between the samples of the discovery cohort, the Validation1_DIA cohort or the Validation3_RNA cohort existed. The data is available through the European Genome-Phenome Archive (EGA) under accession number EGAS00001001746.

#### Validation4_untreated cohort

For the evaluation of the effect of the interaction between trisomy 12 and IGHV mutation status in an untreated context, we took advantage of the CLL cohort published by Tissino et al. in 2020 ^48^. All the cases were from a from a single institution, i.e. the Hematology Unit of the University of Tor Vergata in Rome, and analysed at the CRO in Aviano for IGHV gene status, FISH categories (del(17p), del(11q), trisomy 12, and del(13q)) and *TP53* mutations. This comprised in total 620 patients. Time to first treatment was used as the clinical endpoint.

#### Validation5_ibrutinib cohort

For the evaluation of the effect of the interaction between trisomy 12 and IGHV mutation status in an ibrutinib treated context we took advantage of a retrospective study cohort of 620 CLL patients homogeneously treated with ibrutinib at the Ohio State university.

### Statistical analyses

All statistical analyses were performed in R. Statistical tests were performed as indicated in the text and figures. Wilcoxon signed-rank tests were always two-sided. Boxplots are defined as follows: center line, median; box limits, upper and lower quartiles; whiskers, 1.5x interquartile range. Code used is available at github.

#### Analysis of differential proteins and mRNA

Differential protein abundance between samples with different genetic alterations was assessed with limma ^49^ and DEqMS ^50^ (differential proteins defined as adjusted p <0.001 and |log2FC| >0.5) (github code: “limmaProteomics”). Differential gene expression was performed on the raw count values using DESeq2 ^46^ (differential genes defined as adjusted p <0.001 and |log2FC| >1.5; github code: “RNASeq”).

For samples with known trisomy 12 and IGHV mutation status (n_samples_ = 59), we estimated the proteome-wide (n_proteins_ = 7311) relative effect of trisomy 12 in the context of IGHV-status by using the following design matrix in DeqMS: ∼ IGHV+Trisomy 12+IGHV:Trisomy, and extracted results with respect to the interaction coefficient. Enrichment analysis was performed in the sorted by fold-change gene vector using fgsea ^51^ R package and MSigDB KEGG canonical pathways ^52^.

For overlapping gene symbols and samples in transcriptomics and proteomics data (n = 6310, m = 50), we estimated mRNA – protein correlation for each gene i using the first coefficient in the following model:

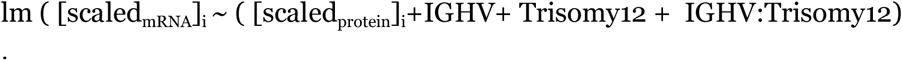

Significant correlated pathways were identified by two-sided t-tests between the mRNA-protein coefficients of leading edge genes (>=5) versus all genes at a Benjamini-Hochberg FDR level of 0.01.

### Protein-mRNA correlation

For each protein-mRNA pair the Spearman correlation was calculated. Cumulative distribution functions of the correlation coefficients were compared using a two-sided Kolmogorov-Smirnov test (github code: “RNASeq”).

### Multi-omics factor analysis

The multi-omics factor analysis was performed on genetics, transcriptomics, and proteomics datasets using the MOFA R package ^30^. Only the 2000 proteins with the highest variance were used. The MOFA model was calculated 10 times and the model with the highest evidence lower bound (ELBO) was chosen. The convergence threshold was set to 0.01. Only factors which explained at least 1.5 % of variance were kept for further analysis (github code: “perfom_MOFA_analysis”).

### *In vivo* lymphocyte growth rate

Patients who had lymphocyte counts available for less than 4 timepoints between the sample collection date and the time of the next treatment and patients currently in treatment were excluded, leaving 35 patients. Lymphocyte growth rates were calculated by fitting a linear model to the log10 transformed lymphocyte counts versus the period of time.

### Analysis of outcome

#### Time-to-next-treatment

Time to next treatment (TTNT) was calculated from the date of sample collection to subsequent treatment initiation. Patients without treatment initiation during observation time and patients who died before treatment initiation were censored at the latest follow-up contact. Proportional hazards regression (Cox regression) was used to explore the potential impact of protein abundances on TTNT using the R package survival. To draw Kaplan-Meier curves the survminer package was used (github code: “CoxRegr”).

#### Time-to-first-treatment

Time to first treatment was calculated from the date of diagnosis to subsequent treatment initiation. Patients without treatment initiation during observation time and patients who died before treatment initiation were censored at the latest follow-up contact. To draw Kaplan-Meier curves the survminer package was used.

#### Overall survival

Overall survival (OS) was calculated from the date of sample collection to the date of death. Patients who did not die within the observation period were censored at the latest follow-up contact. Proportional hazards regression (Cox regression) was used to explore the potential impact of protein abundances on OS using the R package survival. To draw Kaplan-Meier curves the survminer package was used.

TTNT or OS were used as the primary metrics for outcome analysis for groups identified by proteomic or transcriptomic data. For groups stratified from genetic alterations (e.g. IGHV status and Trisomy 12) TTFT or PFS was used.

### Dimensionality reduction and consensus clustering

Consensus clustering on proteins or transcripts was performed using the ConsensusClusterPlus package ^53^ (github code: “ConsensusClustering”). The optimal number of clusters was determined based on cluster stabilities (Data SI5). This supported clustering of proteomic data into 4-6 clusters. As the increase from 5 to 6 clusters led to the subdivision of trisomy 12 patients into U-CLL and M-CLL, indicating biological meaningfulness of the clusters, 6 clusters were chosen. mRNA level clustering into less than 5 groups was not justifiable from the relative change in area under the CDF curve (Data SI6). Clustering into more than 5 subgroups led to splitting off of individual patients. Therefore, 5 clusters were chosen as optimal.

Dimensionality reduction was done by T-distributed stochastic neighbor embedding (t-SNE; Rtsne package), principal component analysis (stats package), and hierarchical clustering (pheatmap package) (github code: “DimensionReduction”).

### Analysis of differential splicing

#### RNA level

For the alternative splicing analysis of the discovery cohort, 59 single-read stranded total RNA-seq samples were processed. First, RNA-seq reads were aligned to the human reference genome (build hs37d5, based on NCBI GRCh37, hg19) using STAR (v2.5.2a) with standard parameters. Aligned RNA-seq data were then subjected to quality control by RNASeQC (v1.1.8) to ensure the integrity of the transcriptome dataset. The same procedure was applied to the Validation3_RNA cohort of 169 paired end RNA-seq samples.

Our alternative splicing analyses largely rely on rMATS, which we employed to identify exon skipping events, mutually exclusive exons, alternative 5’ and 3’ splice sites, and intron retention in individual CLL patient RNA-seq samples. The rMATS tool was also used to quantify alternative splicing activity by calculating estimates for the percent spliced-in (PSI) values associated with each event in a given sample. The entirety of PSI values for a specific sample will be referred to as the sample’s splicing profile in the following and can be seen as characterizing alternative splicing activity in the corresponding CLL patient.

In the case of the discovery cohort, where samples were independently assigned a proteomics group label, rMATS was additionally used to perform a statistical analysis of differences in alternative splicing patterns between ASB-CLL patients and the remaining samples. Based on the results of such a differential alternative splicing analysis within the skipped exon category, we selected the most interesting events in terms of statistical significance (BH-adjusted p-value smaller than 1%) and effect size (absolute difference of groupwise mean PSI values larger than 0.1) for further consideration. Additionally, we excluded events with too low a coverage in read counts (average number of raw inclusion counts across all samples not larger than 20). A total of 430 exon skipping events satisfied all three constraints, 427 of which were also detected in the Validation3_RNA cohort. Reading rMATS output into R and selecting events was done with the help of the maser package.

In order to assess the probability that the observed number of significant exon skipping events was purely due to chance, we performed a permutation test. Specifically, we uniformly drew nine samples from the total list of 59 CLL samples in the discovery cohort without replacement and labelled them as “ASB-CLL”. The remaining 50 samples were labelled as “other”. We then ran rMATS on these randomly labelled samples using the same parameters as in the original analysis and again determined the number N_e_ of significant exon skipping events. This process was repeated for multiple such label permutations, thus generating an estimate for the null distribution of N_e_. The tested null hypothesis is that there is no differential splicing between ASB-CLL and the other proteomics groups.

Apart from the supervised selection of exon skipping events described further above, we performed unsupervised exploratory analysis for each alternative splicing event category in the discovery cohort. We did so by using the 1000 most variable events across all patients in each category. For each patient sample, we then calculated the mean PSI value based on those 1000 events. In order to quantify whether the mean PSI values from ASB-CLL samples tend to be different from those in all other PGs, we performed a two-sided Wilcoxon rank-sum test. The outcome of that test gave us an idea about overall alternative splicing differences in the given category.

In order to capture the characteristic splicing profile of ASB-CLL samples, we used the caret package to train a partial least squares discriminant analysis (PLS-DA) model on the discovery cohort using the PSI values from the 427 selected exon skipping events as predictors. All predictors were centered and scaled prior to training. The PLS-DA classifier has one tuning parameter, namely the number N_c_ of components in the dimensionally reduced space. We determined the optimal value for N_c_ via leave-one-out cross-validation using the area under the ROC curve as the measure of predictive performance to be optimized.

#### Peptide level

Peptide sequences were stripped of modifications and merged by the median followed by mapping to genomic coordinates using the *Splicevista.py* function ^54^ and assigned to Ensembl exon IDs (GRCh38, v92). Each peptide was assigned to exon(s) –in case of splice-junction– based on the initial master protein assignment, but later distributed to multiple proteins supported by the exon(s). Splice-junction-covering peptides were treated as single instances to accommodate the unique quantitative profile deriving from both putative assignments. As such, exons here denote unique coding units, but we follow the exon term for simplification.

Differential splicing was investigated using the limma diffsplice *R* function ^49^. Specifically, for exons quantified in more than two TMT sets (18 samples), we estimated log fold changes of ‘this-cluster-vs-the-rest’ contrasts via moderated t-tests of ‘this-exon-vs-the-rest’ comparisons. Exon-level significance was determined by t-test (FDR corrected p-value <0.01 and |log2FC| >0.5). Enriched groups for significant hits were identified using one-sided Fisher’s exact tests as described above (github code: “CLL_exon_centric_diffSpiced_diffQuant”).

### Gene ontology and KEGG gene set enrichment analyses

Enrichment analysis was performed against KEGG genesets ^55^ using GSEA ^52^. For GO term enrichment ^56^ of proteins loaded onto latent factor 9 the MOFA ^30^ internal function *runEnrichmentAnalysis* was used. The required GO terms were downloaded from MSigDB ^57^ (v7.0) and the FDR cut-off was set to >5 %.

### Protein-protein correlation

For the protein core complex analysis protein-protein Pearson correlations were calculated for CORUM complex members and converted into a pairwise interaction matrix as previously described^15^.

For the generation of the protein correlation network proteins with a high standard deviation were selected and pairwise Pearson correlations were calculated (github code: “ppi_network”). The resulting network was visualized in Gephi 0.9.2 and nodes were filtered using a kcore setting of 3. Modularity clustering of the nodes was carried out with a resolution of 0.8. Annotation of the modularity clusters was done by first extracting all proteins belonging to a cluster. Next, any protein in the full overlap dataset (n =7313) with a Pearson correlation above 0.7 to any of the cluster members was included in the target gene set for that cluster. Enrichment was carried out against the MSigDB and the R packages msigdbr and ClusterProfiler were used to calculate enrichments (github code: “enrichment_of_network-msigdb”). See supplementary methods for additional details.

### Validation of ASB-CLL using DIA based proteomics

To validate the existence of ASB-CLL we analysed 167 new patient samples from Sweden and Germany using DIA based proteomics (Validation1_DIA cohort) as well as 36 samples from the original discovery cohort. Four samples were excluded after quality control assessment due to either poor correlation to the in-depth data (2 samples, discovery cohort) or a low number of identifications (2 samples, Validation1_DIA). We trained a *k*-TSP based classifier on the relative MS2-level quantifications of all identified proteins from the KEGG “Spliceosome” and “B-cell receptor signaling” pathways which were identified by at least 3 spectral matches in the training set. After optimization 8 *k*-TSP pairs were chosen. 36 of the original 68 samples analyzed by HiRIEF were run on DIA and samples with a correlation between the DIA and HiRIEF data of >0.4 were used as a training set. Repeated Monte carlo cross validation was used to optimize both the number of pairs and to identify the pairs which best separated ASB-CLL.

For additional validation of ASB-CLL the published proteomics dataset by Eagle et al. ^24^ was used (Validation2_Eagle cohort). The *k*-TSP classifier trained above was used on the Eagle et al. dataset. One TSP-pair had to be excluded due to lack of overlap and the classification threshold was lowered by 1 as a consequence. Mean protein abundances for the genes in the KEGG gene sets “Spliceosome”, “B-cell receptor signalling”, “proteasome”, and “Valine, leucine and isoleucine degradation” were calculated and compared between the ASB-CLL cluster and all other patients.

### Validation of Tris12M-PG, Tris12U-PG, M-PG, and U-PG

For the confirmation of the clinical relevance of Tris12M-PG, Tris12U-PG, M-PG, and U-PG multiple validation cohorts were used. Patients from the Validation1_DIA cohort (n= 165), the Validation4_untreated cohort (n= 620), only comprising untreated CLL samples, and the Validation5_ibrutinib cohort (n= 463), only comprising ibrutinib treated samples, were split into four groups based on IGHV status and trisomy 12. This was possible because Tris12M-PG, Tris12U-PG, M-PG, and U-PG strongly associated with these two genetic alterations. To assess the native disease progression differences in time to first treatment between the four groups were assessed in the Validation1_DIA and the Validation4_untreated cohorts using Kaplan-Meier curves and log-rank test. To test differences between the groups in the context of treatment with ibrutinib time to progression was evaluated in the Validation5_ibrutinib cohort using Kaplan-Meier curves and log-rank test.

## Supporting information

SI1 Proteomics table

SI2 GSEA Results

SI3 Altered Exon usage ASB-CLL

SI4 Screened drugs

SI5 Consensus cluster proteomics

SI6 Consensus cluster mRNA

## Acknowledgements

The authors would like to acknowledge support from Clinical Proteomics Mass Spectrometry at Karolinska University Hospital and Science for Life Laboratory for providing assistance in mass spectrometry sample preparation and data analysis. We would also like to thank Christiane Zgorzelski for technical assistance. We thank Nils Kurzawa, Simon Raffel, Steffanie Göllner, Konrad Herbst and Peter Lichter for helpful discussions. We thank the EMBL genomics core facility for technical support. The authors gratefully acknowledge the data storage service SDS@hd supported by the Ministry of Science, Research and the Arts Baden-Württemberg (MWK), BioMS and SciLifeLab proteogenomics core facility and the German Research Foundation (DFG) through grant INST 35/1314-1 FUGG; as well as grant support from the Swedish Research Council (R.J: 2017-01653; J.L: 2019-04830; R.R: 2020-05917), the Swedish Cancer Society (J.L: CAN2017/685; R.R: 19 0425 Pj), the Swedish Foundation for Strategic Research (RIF14-0046; SB16-0058), Knut and Alice Wallenberg Foundation (R.R.: 2016.0373) the Stockholm County Council (ALF, J.L.: 20160265; R.R.: 20190472), the Erling-Persson Family Foundation, the Cancer Society in Stockholm (J.L:141243 R.J; 174182) and Swedish Childhood Cancer Foundation (R.J: TJ2016-0035; PR2019-0025 and PR2016-0019; M.S.: TJ2019-0023; J.L: PR2016-0059 and PR2019-0071). St.St. and E.T. were supported by DFG grants of subprojects B1 and B2 of SFB1074. VG was supported by the Associazione Italiana Ricerca Cancro (AIRC), Investigator Grant IG-21687, Milan, Italy. The Progetto Ricerca Finalizzata supported VG (PE-2016-02362756) and AZ (RF-2018-12365790), Ministero della Salute, Rome, Italy.

## Data availability

The mass spectrometry proteomics data have been deposited to the ProteomeXchange Consortium via the PRIDE partner repository with the dataset identifiers PXD017453 (Discovery set) and PXD024544 (DIA Validation set). The RNAseq data for the Validation3_RNA cohort is available through the European Genome-Phenome Archive through accession number EGAS00001001746. The data can be easily explored through our web application: https://www.imbi.uni-heidelberg.de/dietrichlab/CLL_Proteomics/

## Code availability

The code not accessing patient-sensitive information has been deposited on github: https://github.com/DietrichLab/Proteogenomics_and_drug_response_CLL

## Supplementary table

**Supplementary table 1:** Characteristics of the patients in the different cohorts used in this study. Number of patients with a specific characteristic are shown. The percentages in brackets represent the number of patients with this specific feature out of all patients for which this information was available. Validation2_Eagle is the dataset published by Eagle et al. in 2015 ^24^. The type of treatment regiments was available to us for the discovery cohort for which 17 patients had been treated with chemo-immunotherapy, while 6 patients had received treatment with novel agents (rituximab + idelalisib or ibrutinib). Specific treatment information was not available for Validation1_DIA, Validation2_Eagle and Validation3_RNA. Validation4_untreated had not received prior treatment. The Validation5_ibrutinib cohort had uniformly been treated with ibrutinib.

|  | Discovery | Validation1_<br>DIA | Validation2_<br>Eagle | Validation3_<br>RNA | Validation4_<br>untreated | Validation5_<br>ibrutinib |
| --- | --- | --- | --- | --- | --- | --- |
| n | 68 (100) | 165 (100) | 18 (100) | 169 (100) | 620 (100) | 463 (100) |
| median follow-up in months | OS: 19, TTNT: 19 | OS: 64, TTNT: 27 | - | OS: 46, TTNT: 37 | TTFT: 83 | PFS: 71 |
| IGHV mutated | 33 (52) | 106 (66) | 9 (50) | 93 (56) | 383 (62) | 118 (25) |
| Trisomy 12 | 21 (27) | 20 (13) | - | 18 (11) | 94 (15) | 100 (22) |
| del17p | 13 (17) | 5 (3) | - | 13 (8) | 56 (9) | - |
| TP53 mutated | 13 (16) | 14 (9) | - | 23 (14) | 80 (14) | - |
| Female | 29 (43) | 59 (36) | 9 (50) | 61 (36) | 258 (42) | - |
| Treatment at sample collection |  |  |  |  |  |  |
| untreated | 45 (66) | 111 (70) | 13 (72) | 85 (54) | 620 (100) | 0 (0) |
| treated | 23 (34) | 47 (30) | 5 (28) | 72 (46) | 0 (0) | 463 (100) |

## Supplementary figures

**Supplementary figure 1:**
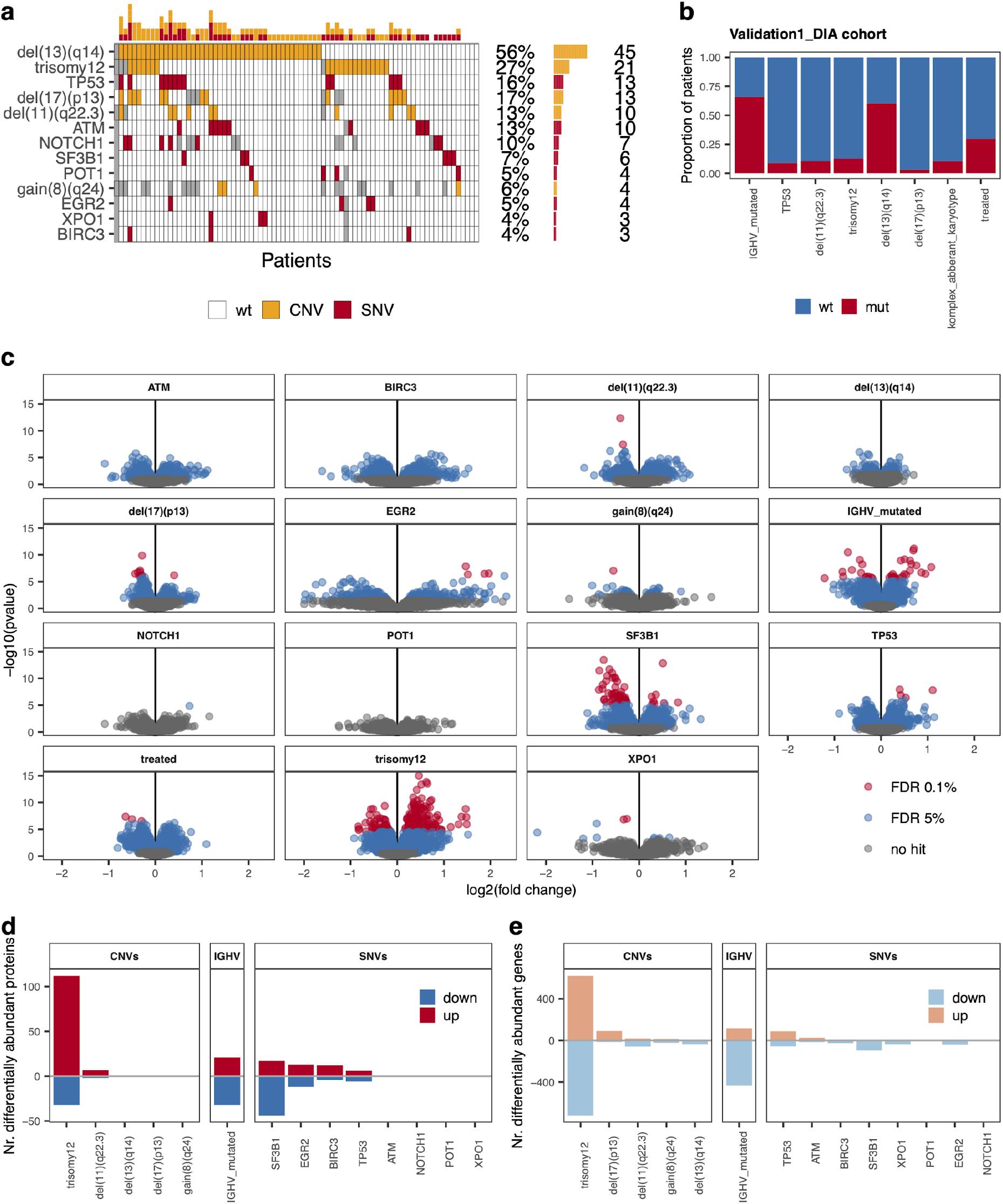
**a,** Overview of the single nucleotide (red, SNV) and copy number variants (yellow, CNV) in the discovery cohort of 68 CLL patients. Grey areas indicate missing values. **b,** Distributions of IGHV, treatment status, the copy number alterations trisomy 12, del(11)(q22.3), del(13)(q14) and del(17)(p13), and the presence of *TP53* mutations in the 165 CLL patients of the Validation1_DIA cohort. **c,** Analysis of differential protein abundances for recurrent SNVs, CNVs, IGHV and treatment status using limma. FDR rates of 0.1 % and 5 % are color-coded. **d,** Number of significantly differentially abundant proteins (FDR 5%; |log2FC| >0.5) and **e,** differentially expressed genes (FDR 5%; |log2FC| >1.5) in relation to recurrent genetic alterations; red/positive numbers =upregulated, blue/negative numbers =downregulated.

**Supplementary figure 2:**
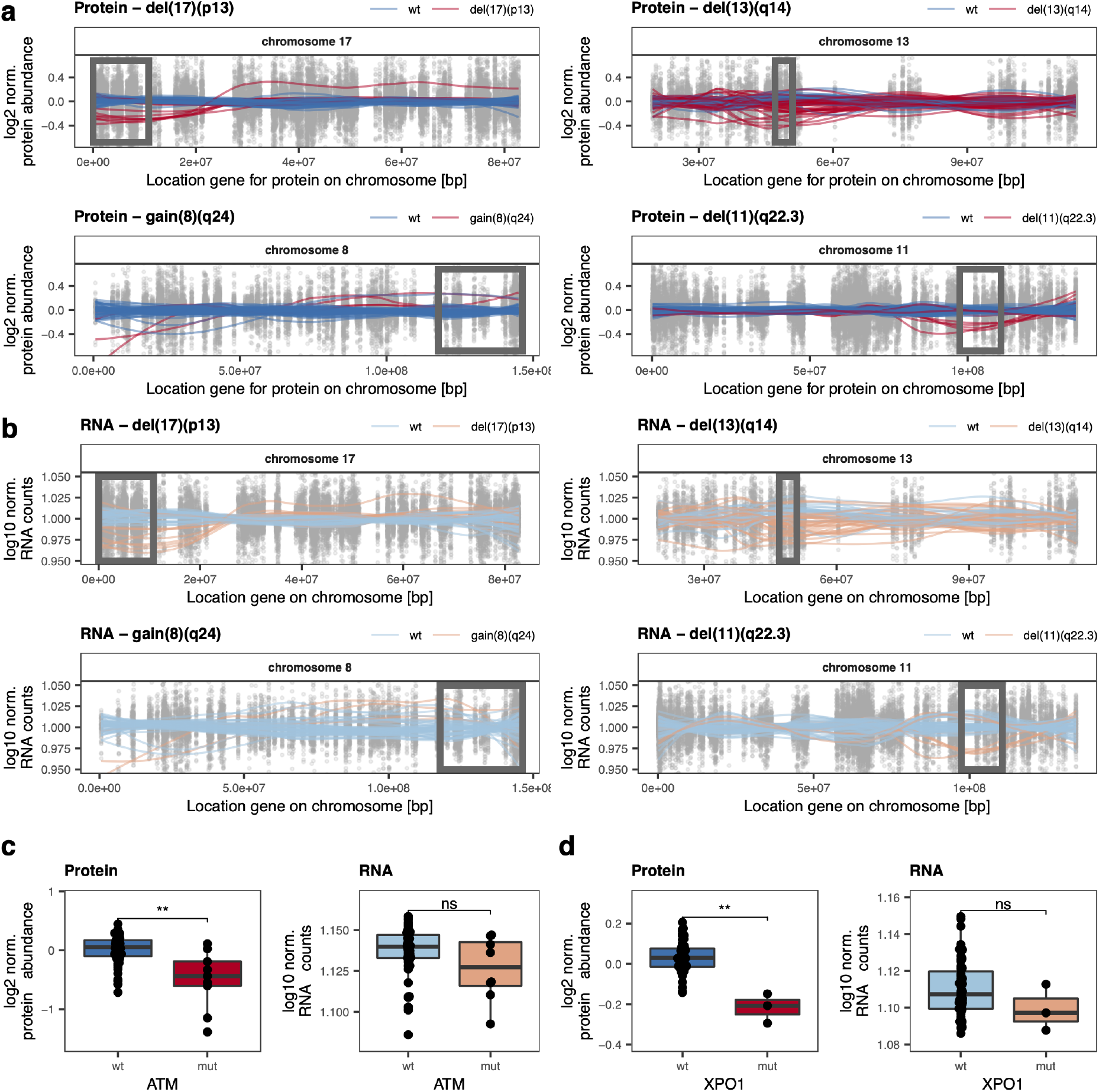
**a,** Effect of different copy-number variations on protein abundances. Normalized protein abundance for the chromosomes affected by the alterations are shown. Points represent individual values for protein - patient pairs. Lines are locally weighted scatterplot smoothed values for individual patients with (red) or without (blue) the alteration. The box is the region affected by the alteration. **b,** Effect of different copy-number variations on gene expression. Normalized gene expression levels for the chromosomes affected by the alterations are shown. Points represent individual values for gene - patient pairs. Lines are locally weighted scatterplot smoothed values for individual patients with (red) or without (blue) the alteration. The box is the region affected by the alteration. **c,** ATM protein (** p =0.001) and transcript levels (not significant p =0.17) in *ATM* mutated (mut) and wild-type (wt) CLL samples; Wilcoxon signed-rank test. **d,** XPO1 protein (** p =0.004) and transcript levels (not significant p =0.18) in *XPO1* mutated (mut) and wild-type (wt) CLL samples; Wilcoxon signed-rank test.

**Supplementary figure 3:**
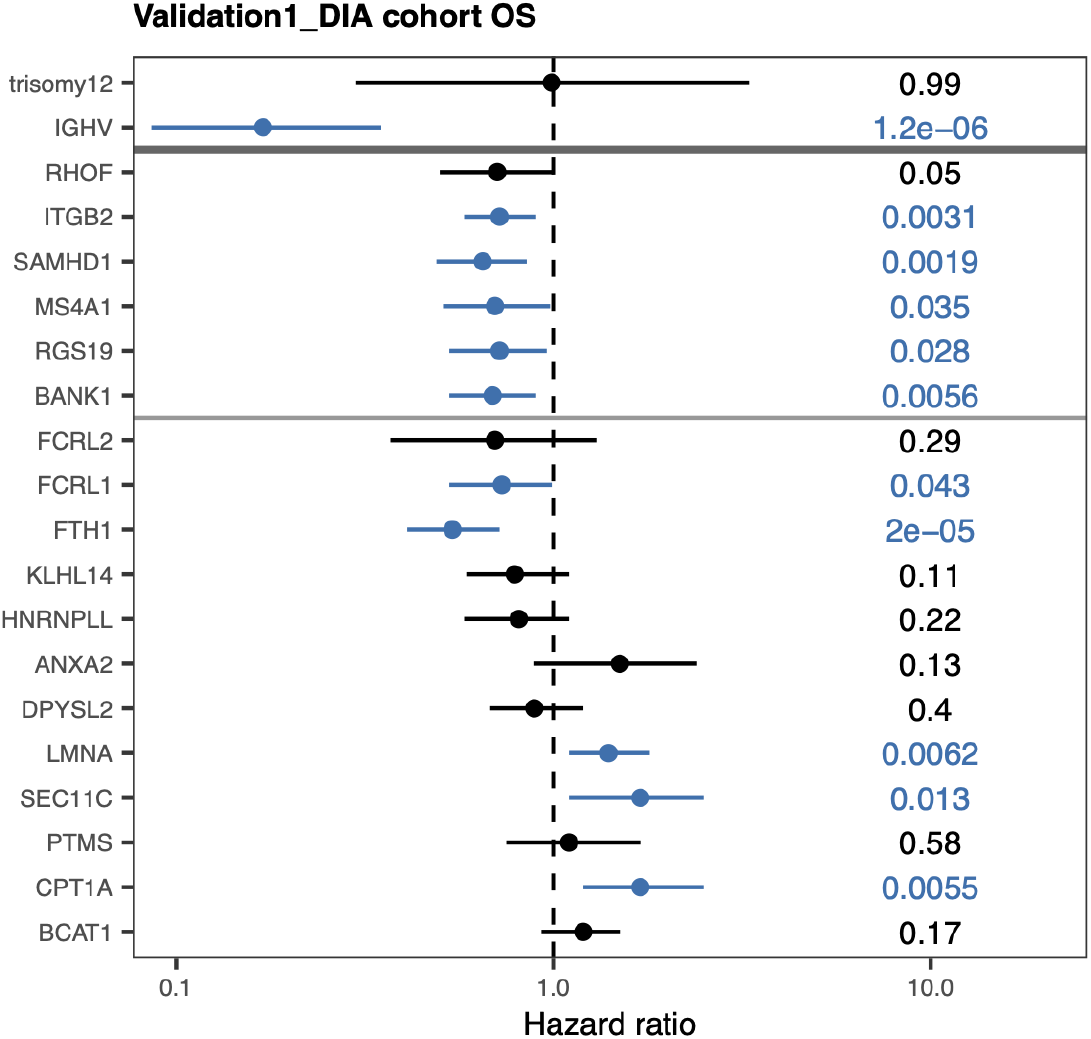
Hazard ratios from Cox regression for overall survival (OS) with genes and proteins in Validation1_DIA dataset with strong weights for LF1 and LF2 in the discovery cohort. P-values are shown on the right. Significant associations (p <0.05) are colored in blue. Mean and 95 % confidence intervals are shown. Out of the proteins with strongest weights on LF9, none were detected in the DIA dataset.

**Supplementary figure 4:**
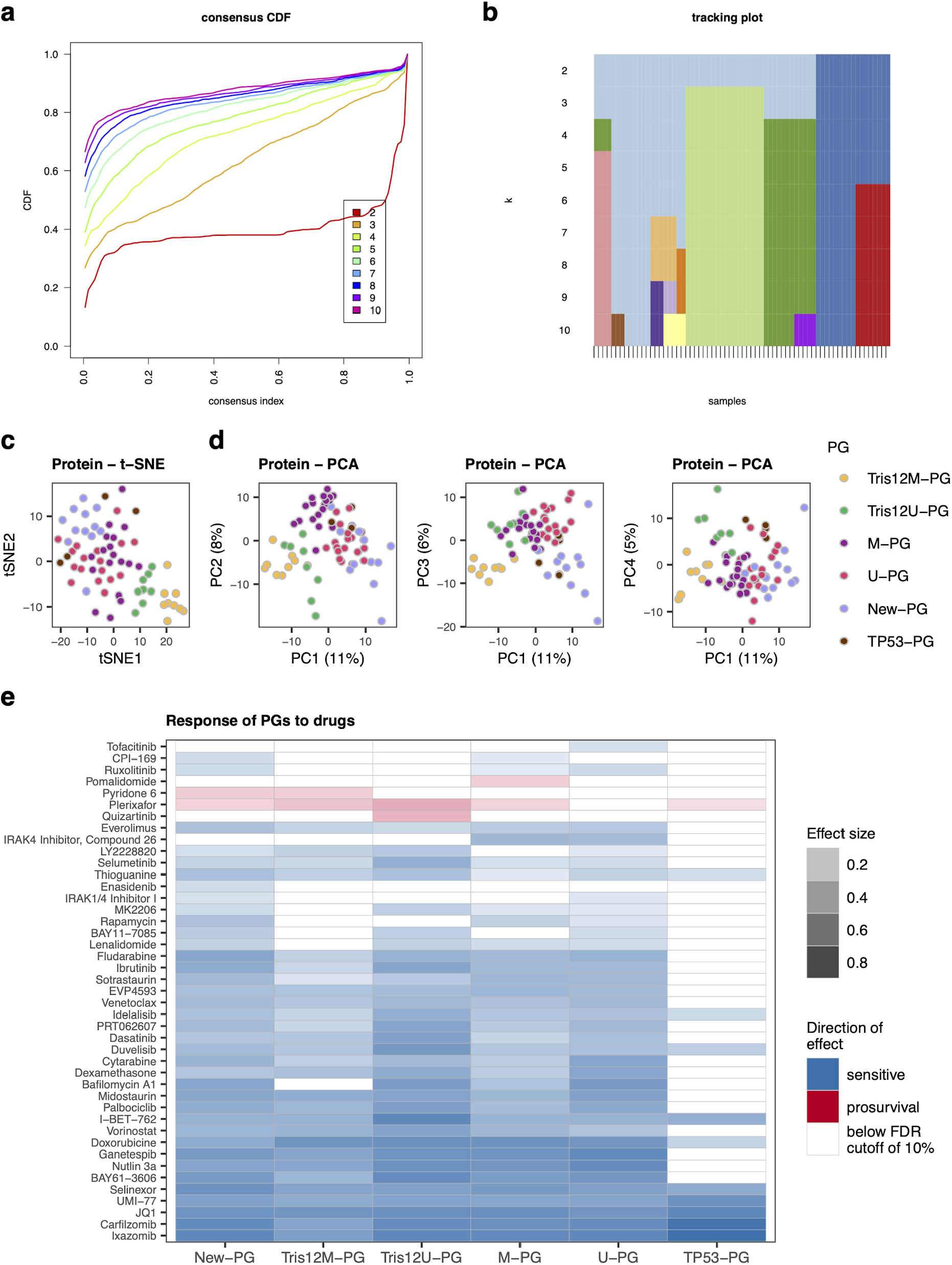
**a,** Cumulative distribution function as produced by the ConsensusClusterPlus package on the proteomics dataset for a number of up to ten clusters. **b,** Tracking plot of clusters as produced by the ConsensusClusterPlus package on the proteomics dataset for a number of up to ten clusters. **c,** t-SNE of proteomics data color coded by Proteomics Groups (PG). **d,** Principal component analysis of proteomics data color coded by PG. **e,** Responses of Proteomics Groups (PG) to the individual drugs tested in the *ex-vivo* drug sensitivity screen. Significance of effects was tested by two sided t-tests against the viability of cells in the control condition.

**Supplementary figure 5:**
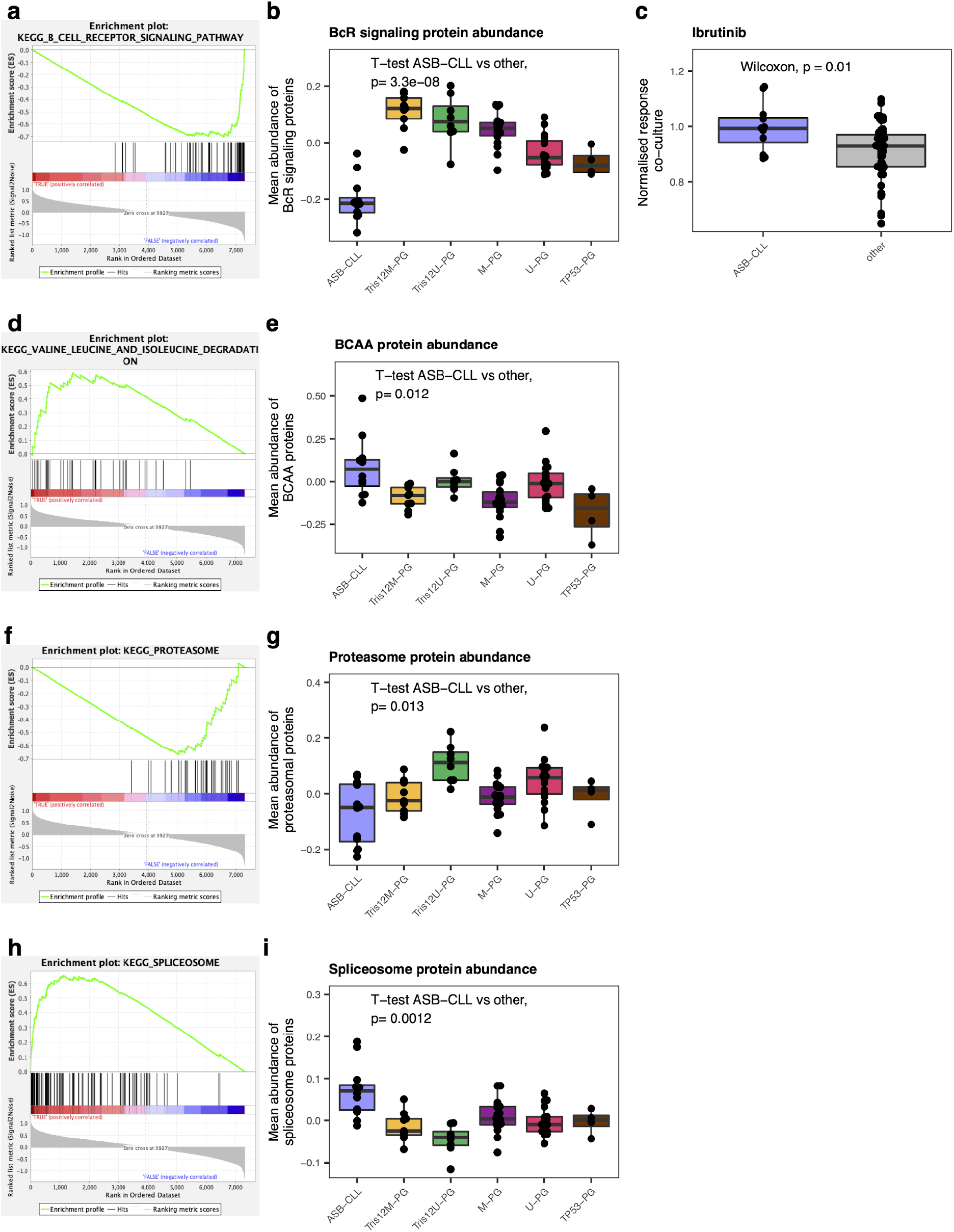
**a,** Enrichment plot for the KEGG pathway “B-cell receptor signaling” for differentially abundant proteins in ASB-CLL. **b,** Mean protein abundance of proteins in KEGG pathway “B-cell receptor signaling” across PGs. **c,** Percentages, normalized to solvent control, of alive cells CLL samples of the discovery cohort in co-culture with the human bone marrow stroma cell line HS-5, treated *ex-vivo* with ibrutinib (40 nM). The comparisons of ASB-CLL samples with the other PGs samples are shown; Wilcoxon signed-rank test. **d,** Enrichment plot for the KEGG pathway “Valine, leucine and isoleucine degradation” for differentially abundant proteins in ASB-CLL. **e,** Mean protein abundance of proteins in KEGG pathway “Valine, leucine and isoleucine degradation” (here termed branched chain amino acid (BCAA) degradation) across PGs. **f,** Enrichment plot for the KEGG pathway “Proteasome” for differentially abundant proteins in ASB-CLL. **g,** Mean protein abundance of proteins in KEGG pathway “Proteasome” across PGs. **h,** Enrichment plot for the KEGG pathway “Spliceosome” for differentially abundant proteins in ASB-CLL. **i,** Mean protein abundance of proteins in KEGG pathway “Spliceosome” across PGs.

**Supplementary figure 6:**
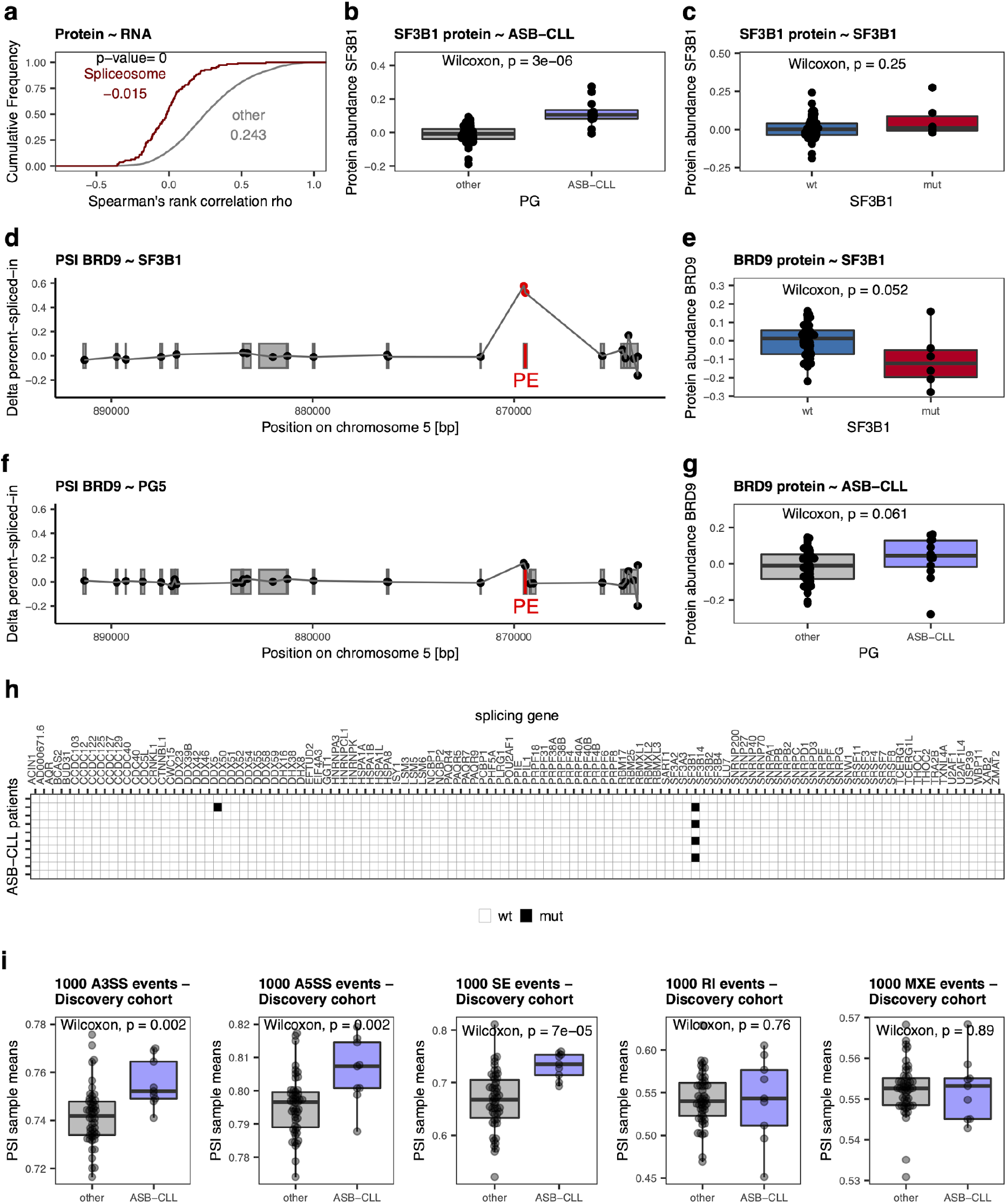
**a,** Cumulative density distribution of protein-mRNA Spearman’s rank correlations for KEGG components of the spliceosome (red) in comparison to all other proteins (gray). A two-sided Kolmogorov-Smirnov test was used to determine the p-value. **b,** SF3B1 log2 protein abundances in ASB-CLL vs. all other groups. **c,** SF3B1 log2 protein abundances in *SF3B1* mutated (mut) vs. wild-type (wt) samples. SF3B1 protein levels were independent of *SF3B1* mutations. **d,** *SF3B1* mutated CLL showed an increase in the percent-spliced-in (PSI) value of the poison exon (PE) in BRD9^32^. **e,** BRD9 log2 protein abundances in *SF3B1* mutated vs. wt CLL. **f,** ASB-CLL did not show altered PSI value of the poison exon in BRD9. **g,** BRD9 log2 protein abundances in ASB-CLL vs. all other groups. **h,** Mutations in genes relevant for splicing in ten ASB-CLL patients, as detected by whole exon sequencing. **i,** Mean PSI value per patient calculated from the 1000 most variable 3’ alternative splice site (A3SS), 5’ alternative splice site (A5SS), skipped exon (SE), retained introns (RI), and mutually exclusive exon (MXE) events across all patients of the discovery cohort. ASB-CLL patients are compared to non-ASB-CLL patients.

**Supplementary figure 7:**
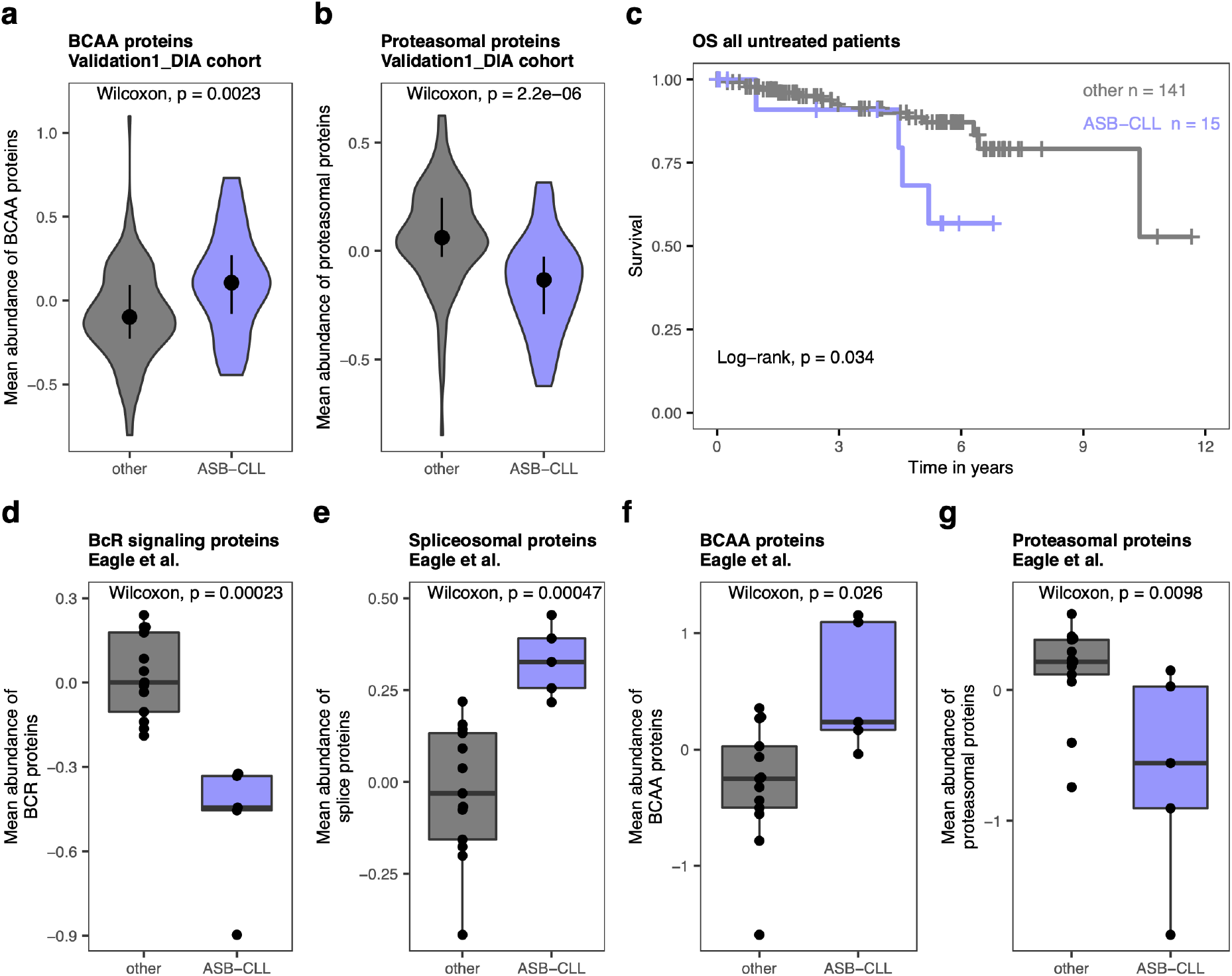
**a,** Violin plot of abundances of proteins involved in branched chain amino acid (BCAA) degradation comparing the subgroup identified as ASB-CLL in the Validation1_DIA dataset to all other patients. **b,** Violin plot of proteasomal protein abundances comparing the subgroup identified as ASB-CLL in the Validation1_DIA dataset to all other patients. **c,** Overall survival (OS) of untreated patients in the discovery and Validation1_DIA cohorts, divided into ASB-CLL and all other patients. **d,** Boxplot of B cell receptor (BcR) signaling protein abundances in the subgroup identified as ASB-CLL in the Validation2_Eagle cohort, in comparison to all other patients. **e,** Boxplot of spliceosomal protein abundances in the subgroup identified as ASB-CLL in the Validation2_Eagle cohort, in comparison to all other patients. **f,** Boxplot of abundances of proteins involved in branched chain amino acid (BCAA) degradation comparing the subgroup identified as ASB-CLL in the Validation2_Eagle dataset to all other patients. **g,** Boxplot of proteasomal protein abundances comparing the subgroup identified as ASB-CLL in the Validation2_Eagle dataset to all other patients.

**Supplementary figure 8:**
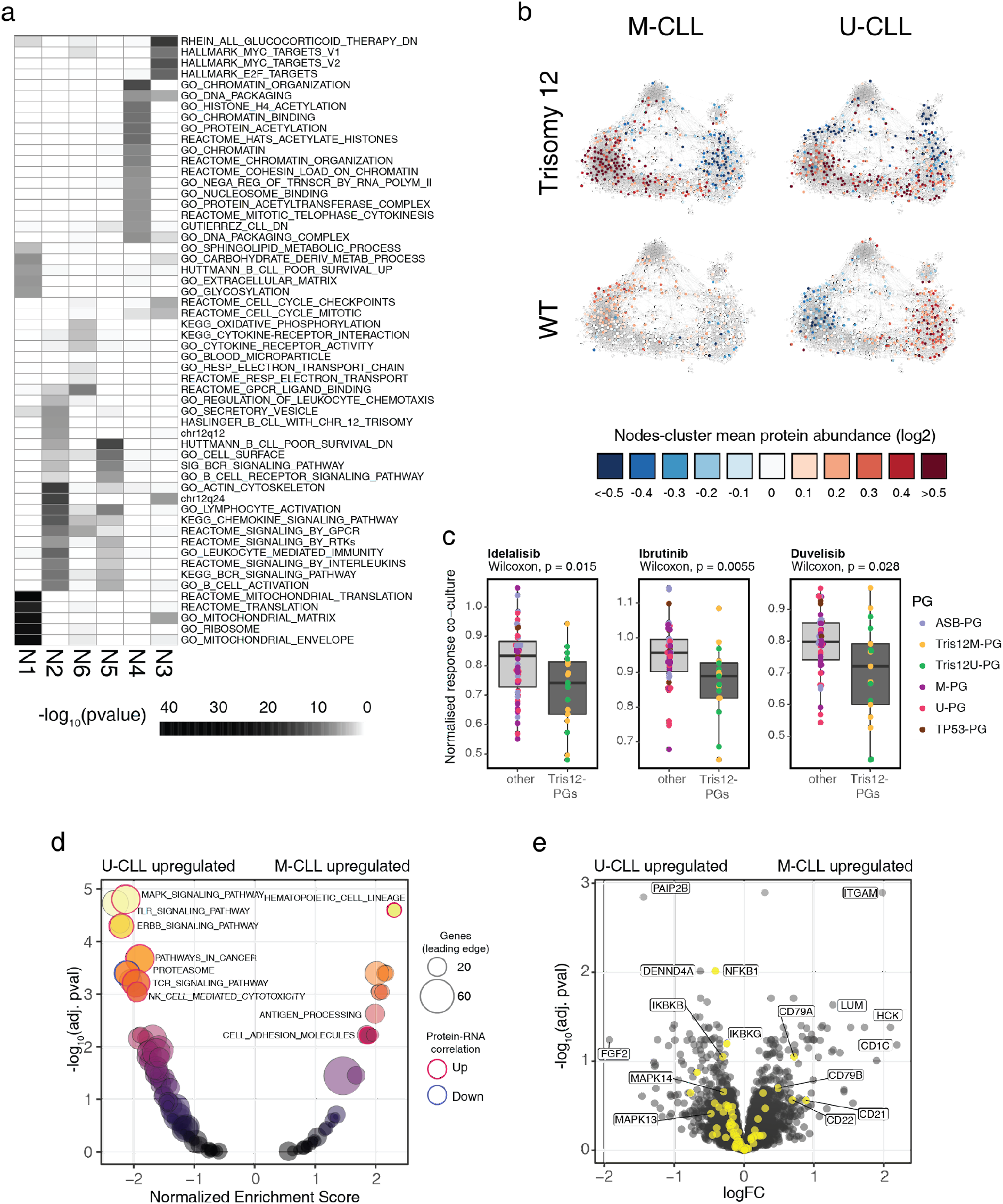
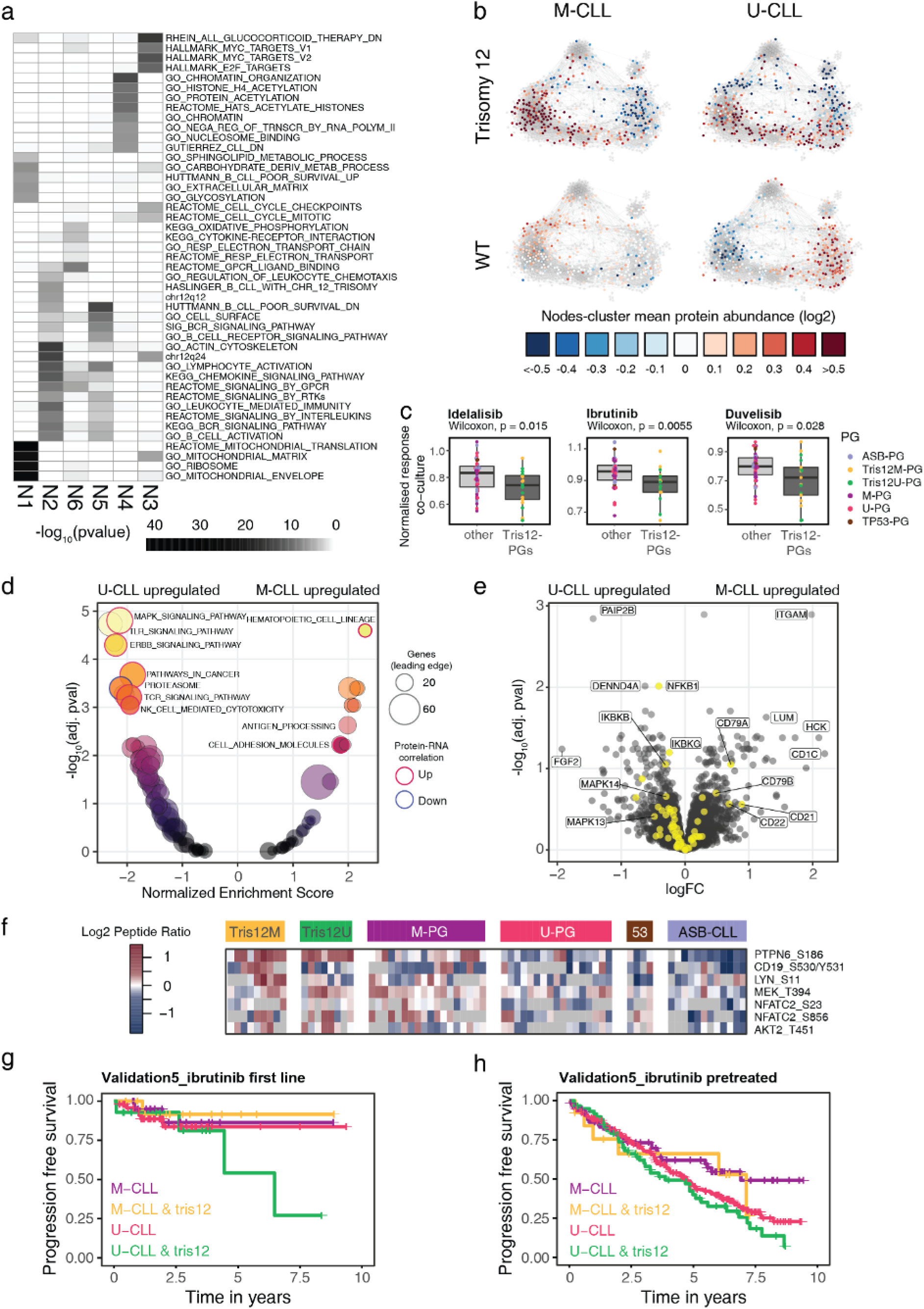
**a**, Heatmap of p-values for selected enrichment terms for the modularity defined clusters N1-N6 in Fig. 7c. **b**, Visualization of the log2 mean, relative protein levels from the Validation1_DIA dataset for patients grouped by IGHV status and trisomy 12 status. Proteins not found in the DIA-dataset are colored grey. (M-CLL, tris12+ n= 14; M-CLL, WT, n= 89; U-CLL, tris12+, n= 6; U-CLL, WT, n= 46), **c,** Percentages, normalized to solvent control, of alive cells CLL samples of the discovery cohort in co-culture with the human bone marrow stroma cell line HS-5, treated *ex-vivo* with idelalisib (9 µM), ibrutinib (40 nM) or duvelisib (4.5 µM). The comparisons of Tris12M-PG & Tris12U-PG samples with the other PGs samples are shown; Wilcoxon signed-rank test. **d,** GSEA analysis results using the KEGG database for protein level differences between U-CLL and M-CLL in the context of trisomy 12. Data was adjusted for differences between U-CLL and M-CLL in cases with disomy 12 (WT). Leading edge genesets with significantly different protein-mRNA correlations are highlighted (red circle = significantly higher correlation, blue circle = Significantly lower correlation) **e,** Volcano plot of DeqMS analysis results of protein level differences between U-CLL and M-CLL in the context of trisomy 12. Data was adjusted for differences between U-CLL and M-CLL in cases with disomy 12 (WT). Genes from the B cell receptor signaling pathway are highlighted in yellow.**g,**

## Supplementary Methods

### IGHV status analysis

RNA was isolated from 1×10^7^PBMCs using TRIZOL reagent (Thermo Fisher Scientific) according to manufacturer’s instructions. cDNA was synthesized from 2 µg RNA using High-capacity cDNA Reverse Transcription Kit (Thermo Fisher Scientific) according to manufacturer’s instructions. PCR reactions as well as the analyses were performed as previously described with minor modifications ^41^. For PCR reactions AmpliTaq Gold DNA polymerase (Thermo Fisher Scientific) with 0.2 µM of each primer and 0.2 mM of each dNTP was used. VH1-, VH3- and VH4-segments were amplified in single reactions whereas primers for VH2, VH3-21, VH5 as well as VH6-segments were run in a multiplex PCR reaction as described ^41^. PCR program was as follows: initial denaturation at 94 °C for 2 minutes, followed by 40 cycles of denaturation (94 °C, 20 seconds), annealing (52 °C, 10 seconds) and elongation (72 °C, 30 seconds) and a final elongation step of 2 minutes at 72 °C. PCR products were sent for Sanger Sequencing (GATC Biotech) using the appropriate forward and the JH-1 reverse primer for the sequencing reaction. In the multiplex PCR reaction both JH-rev as well as JH-1 rev were used for sequencing. After sequencing forward and reverse sequencing results were aligned. To determine the closest matching germline VH-sequence as well as the mutation status, i.e. the percentage of sequence identity, of the VH-segment determined the IMGT/V-Quest-Database was used. The primers for individual PCRs were as follows: PCR1: VH1, JH, JH-1; PCR2: VH3, JH, JH-1; PCR3: VH4, JH, JH-1; PCR4: VH2, VH3-21, VH5, VH6, JH, JH-1 ^41^.

### Panel sequencing of CLL samples

For gene mutation analysis of CLL candidate genes we designed a customized Illumina™ TruSeq Custom Amplicon (TSCA) panel with two independent primer sets for a redundant coverage of *NOTCH1, SF3B1, ATM, TP53, RPS15, BIRC3, MYD88, FBXW7, POT1, XPO1, NFKBIE, EGR2 and BRAF* ^26^*. For ATM, BIRC3, EGR2, FBXW7, MYD88, NFKBIE, POT1* and *TP53* the full gene was covered. For *BRAF* (exons 11-18), *NOTCH1* (exon 34 +3’UTR), *RPS15* (exons 3-4), *SF3B1* (exons 14-16) and XPO1 (exons 14-17) the most commonly affected exons were covered. The selection of these targets comprises the 11 most frequently mutated genes in CLL identified via unbiased whole exome sequencing of 528 CLL patients ^9^. Library preparation was performed using TruSeq Custom Amplicon Assay Kit v1.5 including extension and ligation steps between custom probes. Samples were indexed, pooled and loaded on an Illumina MiSeq flowcell in 32 sample batches.

The cumulative target size was 41,352 basepairs (bp) covered with 304 amplicons in each panel with an amplicon length up to 250 bp. Adjacent 5 intron bp were included to cover splice site mutations. Input of 250 ng DNA from peripheral blood mononuclear cells was sufficient for libraries according to the Illumina TSCA protocol.

We used a custom bioinformatics pipeline including BWA and Samtools (alignment; ^58^), and Varscan (variant calling and annotation; ^59^). Current databases (COSMIC ^60^, 1000G ^61^, dbSNP145 ^62^, ClinVar ^63^) were taken into consideration to evaluate and report variants above a threshold of 5 % mean variant allele fraction (VAF) as pathogenic/non pathogenic. Only mutations which occurred in at least three patients and an allele frequency of more than 20 % were considered for further analyses.

### In-depth data dependant acquisition mass spectrometry proteomics using HiRIEF

Cell pellets were dissolved in Lysis buffer (4 % SDS, 50 mM HEPES pH 7.6, 1 mM DTT), heated to 95° C and sonicated. The total protein amount was estimated (Bio-Rad DC). Samples were then prepared for mass spectrometry analysis using a modified version of the SP3 protein clean-up and a digestion protocol ^42, 64^, where proteins were digested by LysC and trypsin (sequencing grade modified, Pierce). In brief, up to 250 µg protein from each sample was alkylated with 4 mM Chloroacetamide. Sera-Mag SP3 bead mix (20 µl) was transferred into the protein sample together with 100 % Acetonitrile to a final concentration of 70 %. The mix was incubated under rotation at room temperature for 18 min. The mix was placed on the magnetic rack and the supernatant was discarded, followed by two washes with 70 % ethanol and one with 100 % acetonitrile. The beads-protein mixture was reconstituted in 100 µl LysC buffer (0.5 M Urea, 50 mM HEPES pH: 7.6 and 1:50 enzyme (LysC) to protein ratio) and incubated overnight. Finally, trypsin was added in 1:50 enzyme to protein ratio in 100 µl 50 mM HEPES pH 7.6 and incubated overnight. The peptides were eluted from the mixture after placing the mixture on a magnetic rack, followed by peptide concentration measurement (Bio-Rad DC Assay). The samples were then pH adjusted using TEAB pH 8.5 (100 mM final conc.), 65 µg of peptides from each sample were labelled with isobaric TMT-tags (TMT10plex reagent) according to the manufacturer’s protocol (Thermo Scientific). Each set consisted of 9 individual patient samples and the tenth channel contained the same sample pool in each set, consisting of a mixture of patient samples. Sample pools were used as denominators when calculating TMT-ratios and thus served to link the 8 sets together. The tryptic peptides for each set were separated by immobilized pH gradient - isoelectric focusing (IPG-IEF) on 3–10 strips as described previously ^25^.

Of note, the labelling efficiency was determined by LC-MS/MS before pooling of the samples. For the sample clean-up step, a solid phase extraction (SPE strata-X-C, Phenomenex) was performed and purified samples were dried in a SpeedVac. An aliquot of approximately 10 µg was suspended in LC mobile phase A and 1 µg was injected on the LC-MS/MS system.

Online LC-MS was performed as previously described ^15, 25^ using a Dionex UltiMate™ 3000 RSLCnano System coupled to a Q-Exactive-HF mass spectrometer (Thermo Scientific). Each of the 72 plate wells was dissolved in 20 µl solvent A and 10 µl were injected. Samples were trapped on a C18 guard-desalting column (Acclaim PepMap 100, 75 μm x 2 cm, nanoViper, C18, 5 µm, 100 Å), and separated on a 50 cm long C18 column (Easy spray PepMap RSLC, C18, 2 μm, 100 Å, 75 μm x 50 cm). The nano capillary solvent A was 95 % water, 5 % DMSO, 0.1 % formic acid; and solvent B was 5 % water, 5 % DMSO, 95 % acetonitrile, 0.1 % formic acid. At a constant flow of 0.25 μl min^−1^, the curved gradient went from 6-8 % B up to 40 % B in each fraction in a dynamic range of gradient length (see table below), followed by a steep increase to 100 % B in 5 min. FTMS master scans with 60,000 resolution (and mass range 300-1500 m/z) were followed by data-dependent MS/MS (30 000 resolution) on the top 5 ions using higher energy collision dissociation (HCD) at 30 % normalized collision energy. Precursors were isolated with a 2 m/z window. Automatic gain control (AGC) targets were 1e^6^ for MS1 and 1e^5^ for MS2. Maximum injection times were 100 ms for MS1 and 100 ms for MS2. The entire duty cycle lasted ∼2.5 s. Dynamic exclusion was used with 30 s duration. Precursors with unassigned charge state or charge state 1 were excluded. An underfill ratio of 1 % was used.

Protein and peptide identification and quantification was carried out as previously described ^15, 25^. Briefly, Orbitrap raw MS/MS files were converted to mzML format using msConvert from the ProteoWizard tool suite. Spectra were then searched using MSGF+ (v10072) and Percolator (v2.08), where search results from 8 subsequent fractions were grouped for Percolator target/decoy analysis. All searches were done against the human protein subset of Ensembl 75 in the Galaxy platform. MSGF+ settings included precursor mass tolerance of 10 ppm, fully-tryptic peptides, maximum peptide length of 50 amino acids and a maximum charge of 6. Fixed modifications were TMT-10plex on lysines and peptide N-termini, and carbamidomethylation on cysteine residues; a variable modification was used for oxidation on methionine residues. Quantification of TMT-10plex reporter ions was done using OpenMS project’s IsobaricAnalyzer (v2.0). PSMs found at 1 % FDR (false discovery rate) were used to infer gene identities.

Protein quantification by TMT10plex reporter ions was calculated using TMT PSM ratios to the entire sample set (all 10 TMT-channels) and normalized to the sample median. The median PSM TMT reporter ratio from peptides unique to a gene symbol was used for quantification. Protein false discovery rates were calculated using the picked-FDR method using gene symbols as protein groups and limited to 1% FDR.

Supplementary methods table. HiRIEF fractions gradient length.

| Sample | gradient length |
| --- | --- |
| fraction_01 | 50 min |
| fraction_02 | 50 min |
| fraction_03 | 70 min |
| fraction_04 | 70 min |
| fraction_05 | 90 min |
| fraction_06 | 90 min |
| fraction_07 | 110 min |
| fraction_08 | 110 min |
| fraction_09 | 110 min |
| fraction_10 | 110 min |
| fraction_11 | 110 min |
| fraction_12 | 90 min |
| fraction_13 | 90 min |
| fraction_14 | 90 min |
| fraction_15 | 90 min |
| fraction_16 | 70 min |
| fraction_17 | 50 min |
| fraction_18 | 50 min |
| fraction_19 | 50 min |
| fraction_20 | accumulate in trap column |
| fraction_21 | accumulate in trap column |
| fraction_22 | accumulate in trap column |
| fraction_23 | accumulate in trap column |
| fraction_24 | accumulate in trap column |
| fraction_25 | accumulate in trap column |
| fraction_26 | accumulate in trap column |
| fraction_27 | 50 min |
| fraction_28 | 50 min |
| fraction_29 | 70 min |
| fraction_30 | 50 min |
| fraction_31 | 50 min |
| fraction_32 | accumulate in trap column |
| fraction_33 | accumulate in trap column |
| fraction_34 | accumulate in trap column |
| fraction_35 | accumulate in trap column |
| fraction_36 | 50 min |
| fraction_37 | 50 min |
| fraction_38 | 70 min |
| fraction_39 | 70 min |
| fraction_40 | 50 min |
| fraction_41 | 50 min |
| fraction_42 | 50 min |
| fraction_43 | accumulate in trap column |
| fraction_44 | accumulate in trap column |
| fraction_45 | accumulate in trap column |
| fraction_46 | accumulate in trap column |
| fraction_47 | accumulate in trap column |
| fraction_48 | accumulate in trap column |
| fraction_49 | accumulate in trap column |
| fraction_50 | 50 min |
| fraction_51 | 70 min |
| fraction_52 | 70 min |
| fraction_53 | accumulate in trap column |
| fraction_54 | accumulate in trap column |
| fraction_55 | accumulate in trap column |
| fraction_56 | accumulate in trap column |
| fraction_57 | accumulate in trap column |
| fraction_58 | accumulate in trap column |
| fraction_59 | accumulate in trap column |
| fraction_60 | accumulate in trap column |
| fraction_61 | accumulate in trap column |
| fraction_62 | accumulate in trap column |
| fraction_63 | accumulate in trap column |
| fraction_64 | 50 min |
| fraction_65 | 50 min |
| fraction_66 | 70 min |
| fraction_67 | accumulate in trap column |
| fraction_68 | accumulate in trap column |
| fraction_69 | accumulate in trap column |
| fraction_70 | accumulate in trap column |
| fraction_71 | 50 min |
| fraction_72 | 50 min |

### Mass Spectrometry - DIA-based proteomics

Each sample was dissolved in 200 µl lysis buffer (25 mM HEPES pH 7.6, 4 % SDS, 1 mM DTT), heated at 90° C for 5 min and sonicated for 1 min. The total protein amount was estimated (Bio-Rad DC). Samples were then prepared for mass spectrometry analysis using a modified version of the SP3 protein clean-up and a digestion protocol ^42, 64^, where proteins were digested by LycC and trypsin (sequencing grade modified, Pierce). In brief, 200 µg (or the entire amount if <200 µg protein was available) from each sample was alkylated with 4 mM Chloroacetamide. Sera-Mag SP3 (GE Healthcare products 45152105050250 and 65152105050250, distributed by Thermo Fisher) bead mix (20 µl) was transferred into the protein sample together with 100% Acetonitrile to a final concentration of 70 %. The mix was incubated under rotation at room temperature for 20 min. The mix was placed on the magnetic rack and the supernatant was discarded, followed by two washes with 70 % ethanol and one with 100 % acetonitrile. The beads-protein mixture was reconstituted in 100 µl LycC buffer (0.5 M Urea, 50 mM HEPES pH: 7.6 and 1:50 enzyme (LycC) to protein ratio) and incubated overnight. Finally, trypsin was added in 1:50 enzyme to protein ratio in 100 µl 50 mM HEPES pH 7.6 and incubated overnight. Peptide concentration was measured using Bio-Rad DCC.

50 μg of peptides from each sample were cleaned by SP3 beads. For that, peptides were dried by SpeedVac, and dissolved in 20 µl water. 10 µl beads were added to each tube and mixed by short vortex. 570 µl acetonitrile was added to each sample to reach 95 % ACN. The mixture was incubated for 30 minutes at room temperature. To remove the buffer, the tube was placed on a magnetic rack and incubated for 2 minutes at room temperature. Supernatant was discarded. Magnetic beads were washed by addition of 250 μl of acetonitrile and incubated for 30 seconds on the magnetic stand. Supernatant was discarded and the beads air-dried. Tryptic peptides were detached from the beads by addition of 100 μl of 3 % ACN, 0.1 % FA and transferred to a new tube.

5 μg of peptides from each sample were injected and separated using an Ultimate 3000 RSLCnano system coupled to a Q Exactive HF (Thermo Fischer Scientific, San Jose, CA, USA). Samples were trapped on an Acclaim PepMap nanotrap column (C18, 3 mm, 100 Å, 75 µm x 20 mm, Thermo Scientific), and separated on an Acclaim PepMap RSLC column (C18, 2 µm, 100 Å, 75 µm x 50 cm, Thermo Scientific). Peptides were separated using a gradient of mobile phase A (5 % DMSO, 0.1 % FA) and B (90 % ACN, 5 % DMSO, 0.1 % FA), ranging from 6 % to 30 % B in 180 min with a flow of 0.25 ml/min.

For data independent acquisition (DIA), data was acquired using a variable window strategy. The survey scan was performed at 120,000 resolution from 400-1200 m/z, with a max injection time of 200 ms and target of 1e6 ions. For generation of HCD fragmentation spectra, max ion injection time was set as auto and AGC of 2e5 were used before fragmentation at 28 % normalized collision energy, 30,000 resolution. The sizes of the precursor ion selection windows were optimized to have similar density of precursor m/z. The median size of windows was 18.3 m/z with a range of 15-88 m/z covering the scan range of 400-1200 m/z. Neighbor windows had a 2 m/z overlap.

For protein identification and quantification, all raw files analyzed by Spectronaut using the Direct-DIA option without the use of a spectral library, files were searched against ENSEMBL protein database (GRCh38.98.pep.all.fasta). All parameters were kept as default for protein identification. Briefly, runs were recalibrated using iRT standard peptides in a local and non-linear regression. Precursors, peptides and proteins were filtered with FDR 1 %. The decoy database was created by mutation method. For quantification, only peptides unique to a protein group were used. Protein groups were defined based on gene symbols to obtain a gene symbol centric quantification. Stripped peptide quantification was defined as the top precursor quantity. Protein group quantification was calculated by the median value of the top 3 most abundant peptides. Quantification was performed at the MS2 level based on the peak area. The quantitative values were filtered using the qvalue for each sample. Imputation was not performed at any stage of the quantification data generation.

### Protein-protein correlation

#### Protein complex analysis

Protein core complex information was retrieved from the CORUM website (http://mips.helmholtzmuenchen.de/corum/#download). All complex members were assumed to interact with each other. Protein complex information was converted into a pairwise interaction matrix as previously described ^15^. The distribution of correlations between proteins/genes in known complexes in both the transcriptomics and proteomics data was compared to the distribution of correlations between randomly selected protein/gene pairs.

For the investigation of trisomy 12 related complexes, CORUM complexes which contained genes located on chromosome 12 were kept. The log2-value of the relative median abundance of individual proteins in the trisomy 12 cases compared to non-trisomy 12 cases was calculated and complexes with an overall upregulation and no down regulated proteins were kept.

#### Protein-protein interaction network

In order to identify a protein population with high standard deviation across the samples, we first calculated a modified quantile standard deviation for the log ratios of each full-overlap protein. Therefore, the lowest and highest value was discarded. The distribution of all log quantile standard deviations was then modelled with a mixture of two Gaussian distributions by applying an expectation maximization algorithm (R package mixtools). The converged model with the two underlying distributions was assumed to represent unmodulated and modulated proteins.

Next, we determined the log standard deviation cutoff that optimally separated the modulated from the unmodulated protein population. We probed each percentile as a cutoff for the mixed distributions, calculated the true and false positive rates and averaged the values over ten consecutive runs. The log standard deviation cutoff was selected by optimising the number of true positive minus false positive modulated proteins, eventually rounded to the lower .5 to ensure reproducibility.

Pairwise Pearson correlations between each modulated protein were calculated. In a QQ-plot, we compared our values to a hypothetical normal distribution and roughly set a cutoff where correlation coefficients started to deviate from the diagonal, which was at r = 0.5. All correlations equal or greater than this were translated into edges between protein nodes in an initial protein-protein-interaction network (github code: “ppi_network”). The network was visualized in Gephi 0.9.2. The initial number of nodes (n =1801) were filtered using a Kcore setting of 3 and above and the core network of 1047 nodes was used for further analysis. Modularity clustering of the nodes was carried out with a resolution of 0.8.

Annotation of the modularity clusters was done by first extracting all proteins belonging to a cluster. Next, any protein in the full overlap dataset (n =7313) that had a Pearson correlation above 0.7 to any of the cluster members was included in the target gene set for that cluster. Enrichment using a target-background approach was carried out against the MSigDB categories Hallmark, C1, C2, C5 and C6. The full overlap data set (n =7313) was used as background and the R packages msigdbr and ClusterProfiler were used to calculate enrichments (github code: “enrichment_of_network-msigdb”).

### Analysis of differential splicing

#### RNA level

Due to its short computational time the julia-based tool *whippet* ^65^ was used for visualisation of specific splicing events, as shown in Fig. S6, following the recommended workflow. In brief, an index was created from the human reference genome (GRCh37). Fastq files were quantified using the options for single-end reads. ASB-CLL was compared to all other samples, or *SF3B1* mutated samples were compared to wt samples, by using the *whippet-delta* functions and setting the parameters of *-r* to 20 and *-s* to 3. For visualisation delta percent-spliced-in values were plotted in R.

## Notes

https://www.imbi.uni-heidelberg.de/dietrichlab/CLL_Proteomics/

### Competing Interest Statement

The authors have declared no competing interest.

### Summary of Updates

Author name corrected

## References

1. Hallek, M. Chronic lymphocytic leukemia: 2017 update on diagnosis, risk stratification, and treatment. Am. J. Hematol. 92, 946–965 (2017).

2. Oakes, C. C. et al. DNA methylation dynamics during B cell maturation underlie a continuum of disease phenotypes in chronic lymphocytic leukemia. Nat. Genet. 48, 253–264 (2016).

3. Queirós, A. C. et al. A B-cell epigenetic signature defines three biologic subgroups of chronic lymphocytic leukemia with clinical impact. Leukemia 29, 598–605 (2015).

4. Kulis, M. et al. Whole-genome fingerprint of the DNA methylome during human B cell differentiation. Nat. Genet. 47, 746–756 (2015).

5. Hamblin, T. J., Davis, Z., Gardiner, A., Oscier, D. G. & Stevenson, F. K. Unmutated Ig V(H) genes are associated with a more aggressive form of chronic lymphocytic leukemia. Blood 94, 1848–1854 (1999).

6. Stevenson, F. K., Krysov, S., Davies, A. J., Steele, A. J. & Packham, G. B-cell receptor signaling in chronic lymphocytic leukemia. Blood 118, 4313–4320 (2011).

7. Byrd, J. C. et al. Targeting BTK with ibrutinib in relapsed chronic lymphocytic leukemia. N. Engl. J. Med. 369, 32–42 (2013).

8. Munir, T. et al. Final analysis from RESONATE: Up to six years of follow-up on ibrutinib in patients with previously treated chronic lymphocytic leukemia or small lymphocytic lymphoma. Am. J. Hematol. 94, 1353–1363 (2019).

9. Landau, D. A. et al. Mutations driving CLL and their evolution in progression and relapse. Nature 526, 525–530 (2015).

10. Bosch, F. & Dalla-Favera, R. Chronic lymphocytic leukaemia: from genetics to treatment. Nat. Rev. Clin. Oncol. 16, 684–701 (2019).

11. Puente, X. S. et al. Non-coding recurrent mutations in chronic lymphocytic leukaemia. Nature 526, 519–524 (2015).

12. Wang, L. et al. Transcriptomic Characterization of SF3B1 Mutation Reveals Its Pleiotropic Effects in Chronic Lymphocytic Leukemia. Cancer Cell 30, 750–763 (2016).

13. Döhner, H. et al. Genomic aberrations and survival in chronic lymphocytic leukemia. N. Engl. J. Med. 343, 1910–1916 (2000).

14. Zhang, H. et al. Integrated Proteogenomic Characterization of Human High-Grade Serous Ovarian Cancer. Cell 166, 755–765 (2016).

15. Johansson, H. J. et al. Breast cancer quantitative proteome and proteogenomic landscape. Nat. Commun. 10, 1600 (2019).

16. Akbani, R. et al. A pan-cancer proteomic perspective on The Cancer Genome Atlas. Nat. Commun. 5, 3887 (2014).

17. Yang, M. et al. Proteogenomics and Hi-C reveal transcriptional dysregulation in high hyperdiploid childhood acute lymphoblastic leukemia. Nat. Commun. 10, 1519 (2019).

18. Archer, T. C. et al. Proteomics, Post-translational Modifications, and Integrative Analyses Reveal Molecular Heterogeneity within Medulloblastoma Subgroups. Cancer Cell 34, 396–410.e8 (2018).

19. Sinha, A. et al. The Proteogenomic Landscape of Curable Prostate Cancer. Cancer Cell 35, 414–427.e6 (2019).

20. Clark, D. J. et al. Integrated Proteogenomic Characterization of Clear Cell Renal Cell Carcinoma. Cell 179, 964–983.e31 (2019).

21. Petralia, F. et al. Integrated Proteogenomic Characterization across Major Histological Types of Pediatric Brain Cancer. Cell 183, 1962–1985.e31 (2020).

22. Mayer, R. L. et al. Proteomics and metabolomics identify molecular mechanisms of aging potentially predisposing for chronic lymphocytic leukemia. Mol. Cell. Proteomics 17, 290–303 (2018).

23. Johnston, H. E. et al. Proteomics Profiling of CLL Versus Healthy B-cells Identifies Putative Therapeutic Targets and a Subtype-independent Signature of Spliceosome Dysregulation. Mol. Cell. Proteomics 17, 776–791 (2018).

24. Eagle, G. L. et al. Total proteome analysis identifies migration defects as a major pathogenetic factor in immunoglobulin heavy chain variable region (IGHV)-unmutated chronic lymphocytic leukemia. Mol. Cell. Proteomics 14, 933–945 (2015).

25. Branca, R. M. M. et al. HiRIEF LC-MS enables deep proteome coverage and unbiased proteogenomics. Nat. Methods 11, 59–62 (2014).

26. Tausch, E. et al. Prognostic and predictive impact of genetic markers in patients with CLL treated with obinutuzumab and venetoclax. Blood 135, 2402–2412 (2020).

27. Kienle, D. L. et al. Evidence for distinct pathomechanisms in genetic subgroups of chronic lymphocytic leukemia revealed by quantitative expression analysis of cell cycle, activation, and apoptosis-associated genes. J. Clin. Oncol. 23, 3780–3792 (2005).

28. Dittmer, D. et al. Gain of function mutations in p53. Nat. Genet. 4, 42–46 (1993).

29. Jethwa, A. et al. TRRAP is essential for regulating the accumulation of mutant and wild-type p53 in lymphoma. Blood 131, 2789–2802 (2018).

30. Argelaguet, R. et al. Multi-Omics Factor Analysis-a framework for unsupervised integration of multi-omics data sets. Mol. Syst. Biol. 14, e8124 (2018).

31. Muggen, A. F. et al. Basal Ca(2+) signaling is particularly increased in mutated chronic lymphocytic leukemia. Leukemia 29, 321–328 (2015).

32. Inoue, D. et al. Spliceosomal disruption of the non-canonical BAF complex in cancer. Nature 574, 432–436 (2019).

33. Shen, S. et al. rMATS: robust and flexible detection of differential alternative splicing from replicate RNA-Seq data. Proc. Natl. Acad. Sci. U. S. A. 111, E5593–601 (2014).

34. Hüttmann, A. et al. Gene expression signatures separate B-cell chronic lymphocytic leukaemia prognostic subgroups defined by ZAP-70 and CD38 expression status. Leukemia 20, 1774–1782 (2006).

35. Kipps, T. J. et al. Long-Term Studies Assessing Outcomes of Ibrutinib Therapy in Patients With Del(11q) Chronic Lymphocytic Leukemia. Clin. Lymphoma Myeloma Leuk. 19, 715–722.e6 (2019).

36. Dong, F. et al. Identification of survival-related predictors in hepatocellular carcinoma through integrated genomic, transcriptomic, and proteomic analyses. Biomed. Pharmacother. 114, 108856 (2019).

37. Li, C. et al. Integrated Omics of Metastatic Colorectal Cancer. Cancer Cell 38, 734–747.e9 (2020).

38. Zenz, T. et al. TP53 mutation and survival in chronic lymphocytic leukemia. J. Clin. Oncol. 28, 4473–4479 (2010).

39. Burger, J. A. & Chiorazzi, N. B cell receptor signaling in chronic lymphocytic leukemia. Trends Immunol. 34, 592–601 (2013).

40. Baliakas, P. et al. Tailored approaches grounded on immunogenetic features for refined prognostication in chronic lymphocytic leukemia. Haematologica 104, 360–369 (2019).

41. Szankasi, P. & Bahler, D. W. Clinical laboratory analysis of immunoglobulin heavy chain variable region genes for chronic lymphocytic leukemia prognosis. J. Mol. Diagn. 12, 244–249 (2010).

42. Moggridge, S., Sorensen, P. H., Morin, G. B. & Hughes, C. S. Extending the Compatibility of the SP3 Paramagnetic Bead Processing Approach for Proteomics. J. Proteome Res. 17, 1730–1740 (2018).

43. Boekel, J. et al. Multi-omic data analysis using Galaxy. Nat. Biotechnol. 33, 137–139 (2015).

44. Dobin, A. et al. STAR: ultrafast universal RNA-seq aligner. Bioinformatics 29, 15–21 (2013).

45. Anders, S., Pyl, P. T. & Huber, W. HTSeq—a Python framework to work with high-throughput sequencing data. Bioinformatics 31, 166–169 (2015).

46. Love, M. I., Huber, W. & Anders, S. Moderated estimation of fold change and dispersion for RNA-seq data with DESeq2. Genome Biol. 15, 550 (2014).

47. Dietrich, S. et al. Drug-perturbation-based stratification of blood cancer. J. Clin. Invest. 128, 427–445 (2018).

48. Tissino, E. et al. CD49d promotes disease progression in chronic lymphocytic leukemia: new insights from CD49d bimodal expression. Blood 135, 1244–1254 (2020).

49. Ritchie, M. E. et al. limma powers differential expression analyses for RNA-sequencing and microarray studies. Nucleic Acids Res. 43, e47 (2015).

50. Zhu, Y. et al. DEqMS: a method for accurate variance estimation in differential protein expression analysis. Mol. Cell. Proteomics (2020) doi:10.1074/mcp.TIR119.001646.

51. Korotkevich, G. et al. Fast gene set enrichment analysis. Cold Spring Harbor Laboratory 060012 (2021) doi:10.1101/060012.

52. Subramanian, A. et al. Gene set enrichment analysis: A knowledge-based approach for interpreting genome-wide expression profiles. Proc. Natl. Acad. Sci. U. S. A. 102, 15545–15550 (2005).

53. Wilkerson, M. D. & Hayes, D. N. ConsensusClusterPlus: a class discovery tool with confidence assessments and item tracking. Bioinformatics 26, 1572–1573 (2010).

54. Zhu, Y. et al. SpliceVista, a tool for splice variant identification and visualization in shotgun proteomics data. Mol. Cell. Proteomics 13, 1552–1562 (2014).

55. Kanehisa, M., Sato, Y., Furumichi, M., Morishima, K. & Tanabe, M. New approach for understanding genome variations in KEGG. Nucleic Acids Res. 47, D590–D595 (2019).

56. The Gene Ontology Consortium. The Gene Ontology Resource: 20 years and still GOing strong. Nucleic Acids Res. 47, D330–D338 (2019).

57. Liberzon, A. et al. Molecular signatures database (MSigDB) 3.0. Bioinformatics 27, 1739–1740 (2011).

58. Li, H. A statistical framework for SNP calling, mutation discovery, association mapping and population genetical parameter estimation from sequencing data. Bioinformatics 27, 2987–2993 (2011).

59. Koboldt, D. C. et al. VarScan 2: somatic mutation and copy number alteration discovery in cancer by exome sequencing. Genome Res. 22, 568–576 (2012).

60. Tate, J. G. et al. COSMIC: the Catalogue Of Somatic Mutations In Cancer. Nucleic Acids Res. 47, D941–D947 (2019).

61. Voight, B. F. et al. A global reference for human genetic variation. Nature 526, 68–74 (2015).

62. Kitts, A., Phan, L., Ward, M. & Holmes, J. B. The Database of Short Genetic Variation (dbSNP). in The NCBI Handbook [Internet]. 2nd edition (National Center for Biotechnology Information (US), 2014).

63. Landrum, M. J. et al. ClinVar: improving access to variant interpretations and supporting evidence. Nucleic Acids Res. 46, D1062–D1067 (2018).

64. Hughes, C. S. et al. Ultrasensitive proteome analysis using paramagnetic bead technology. Mol. Syst. Biol. 10, 757 (2014).

65. Sterne-Weiler, T., Weatheritt, R. J., Best, A. J., Ha, K. C. H. & Blencowe, B. J. Efficient and Accurate Quantitative Profiling of Alternative Splicing Patterns of Any Complexity on a Laptop. Mol. Cell 72, 187–200.e6 (2018).

