## Supplementary material for "Proteogenomics refines the molecular classification of chronic lymphocytic leukemia": SI5 Consensus cluster proteomics

consensus matrix legend

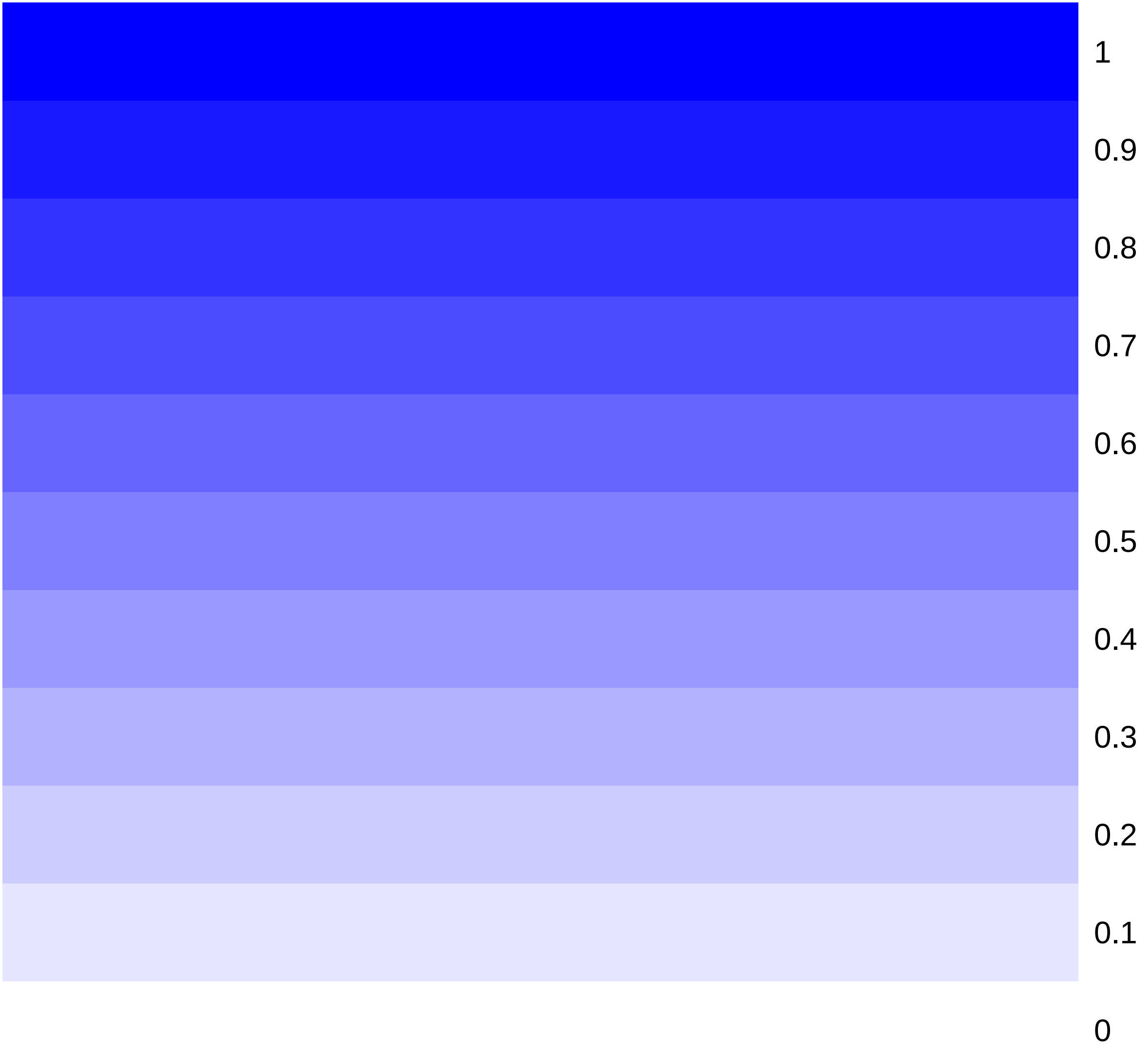

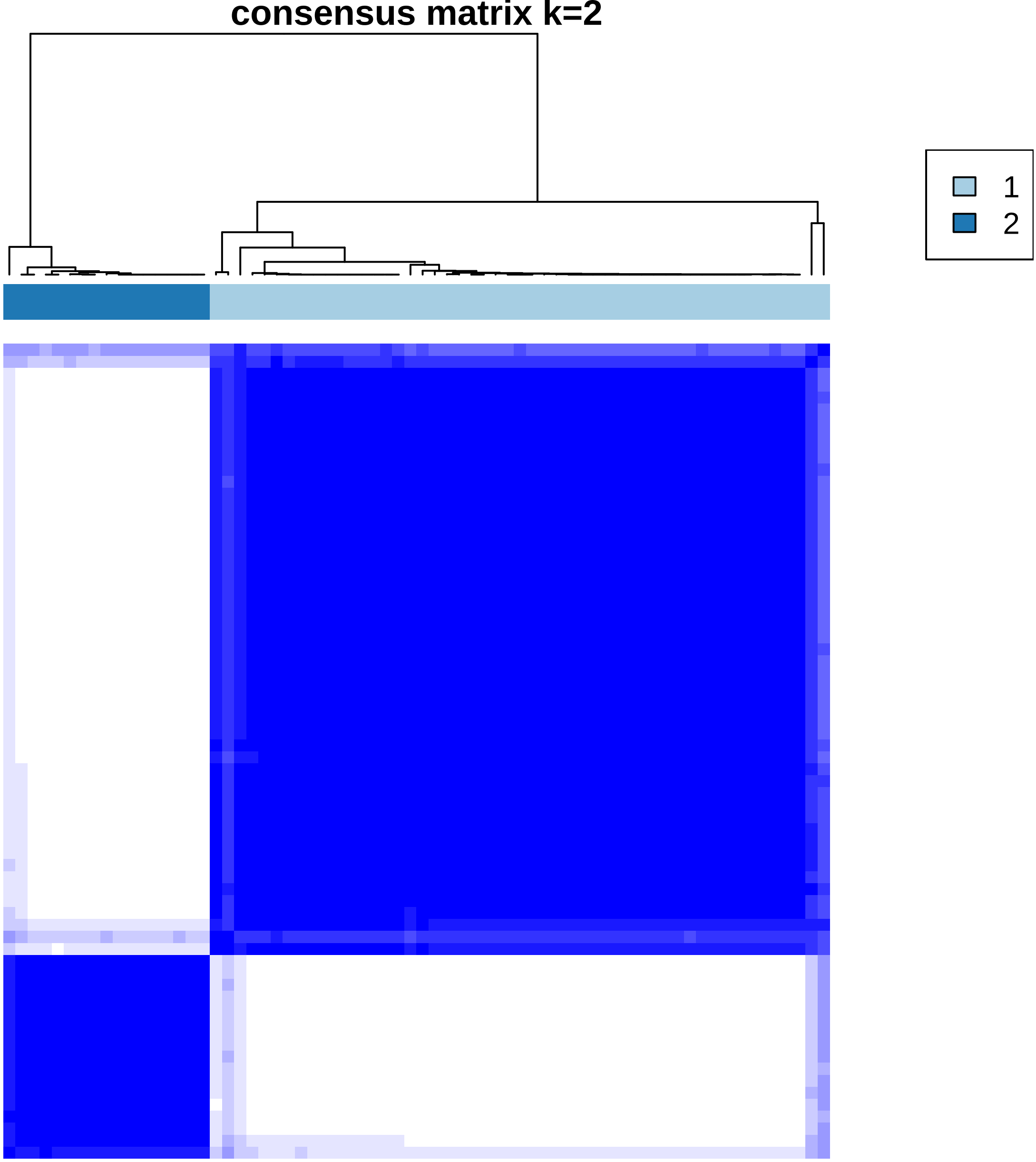
consensus matrix k=2
1
2

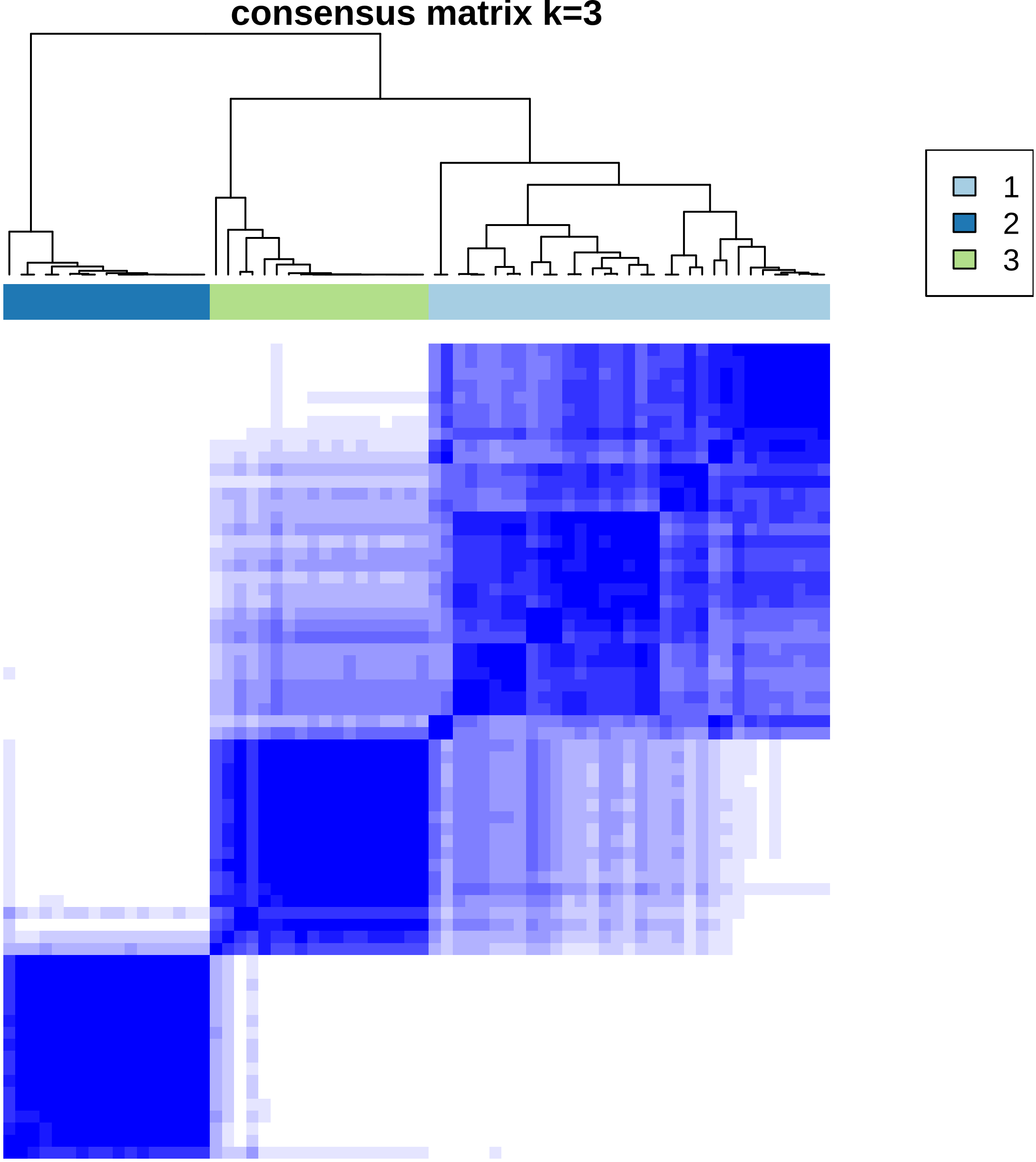
consensus matrix k=3
1
2
3

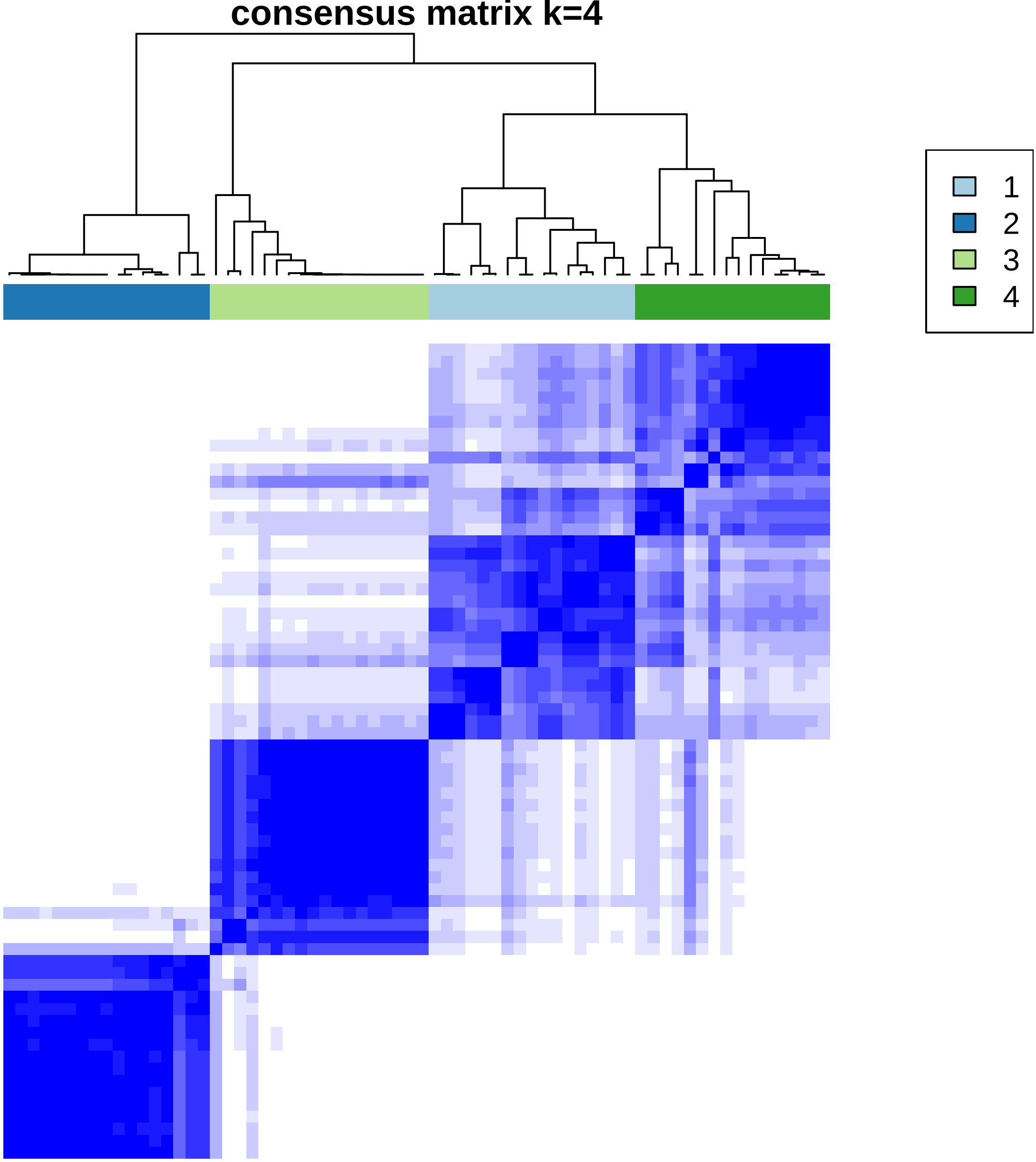
consensus matrix k=4
1
2
3
4

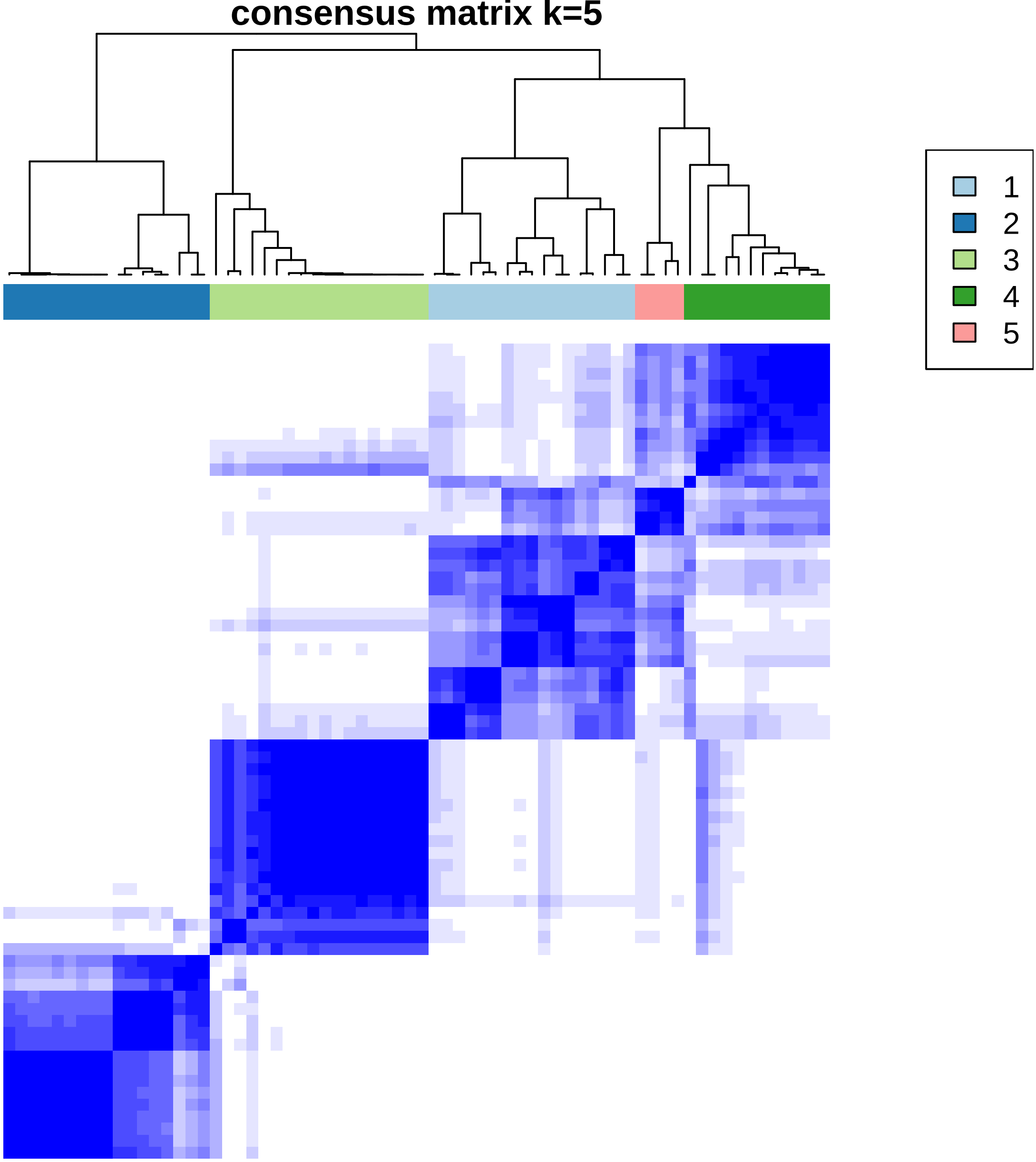
consensus matrix k=5
1
2
3
4
5

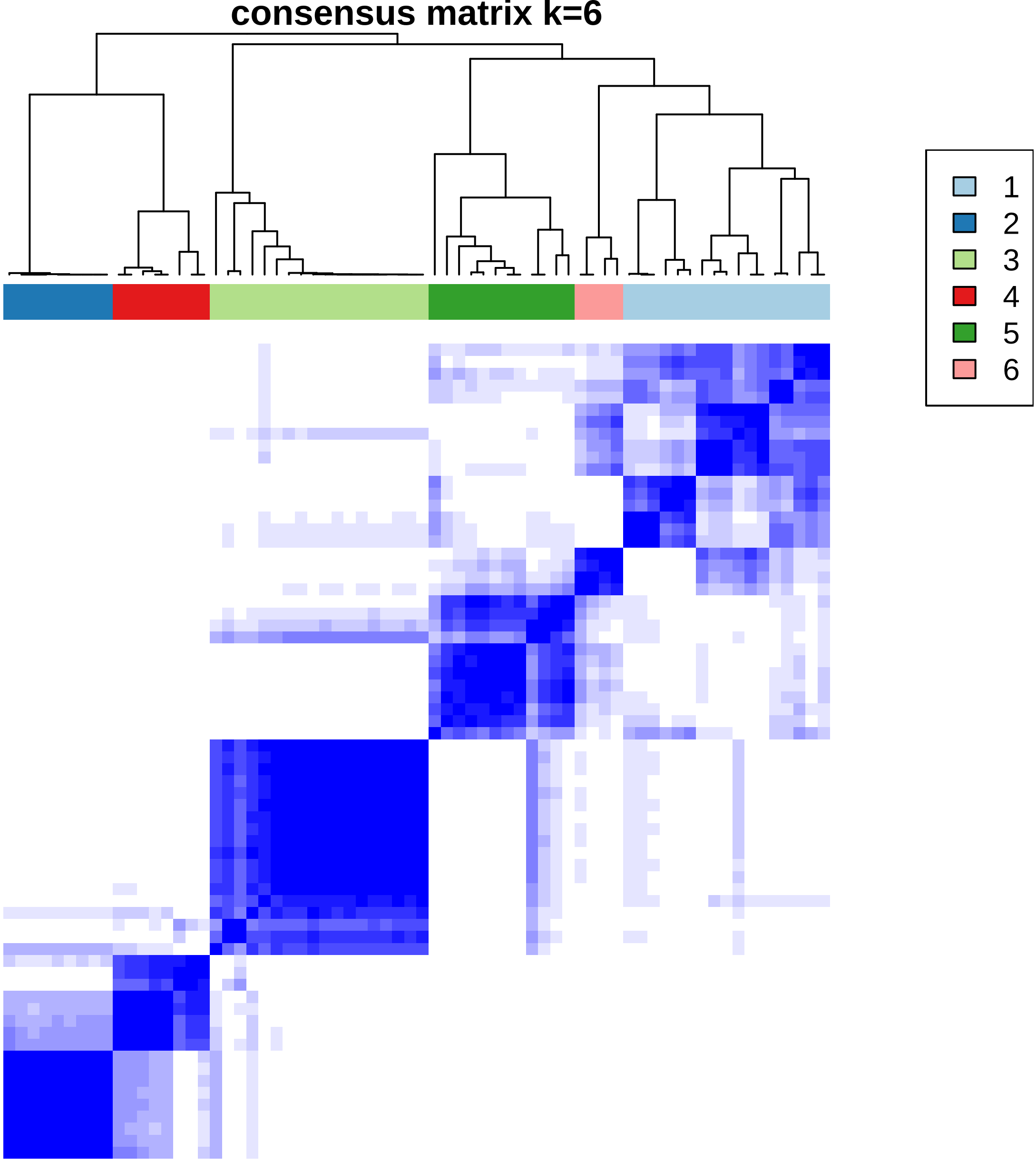
consensus matrix k=6
1
2
3
4
5
6

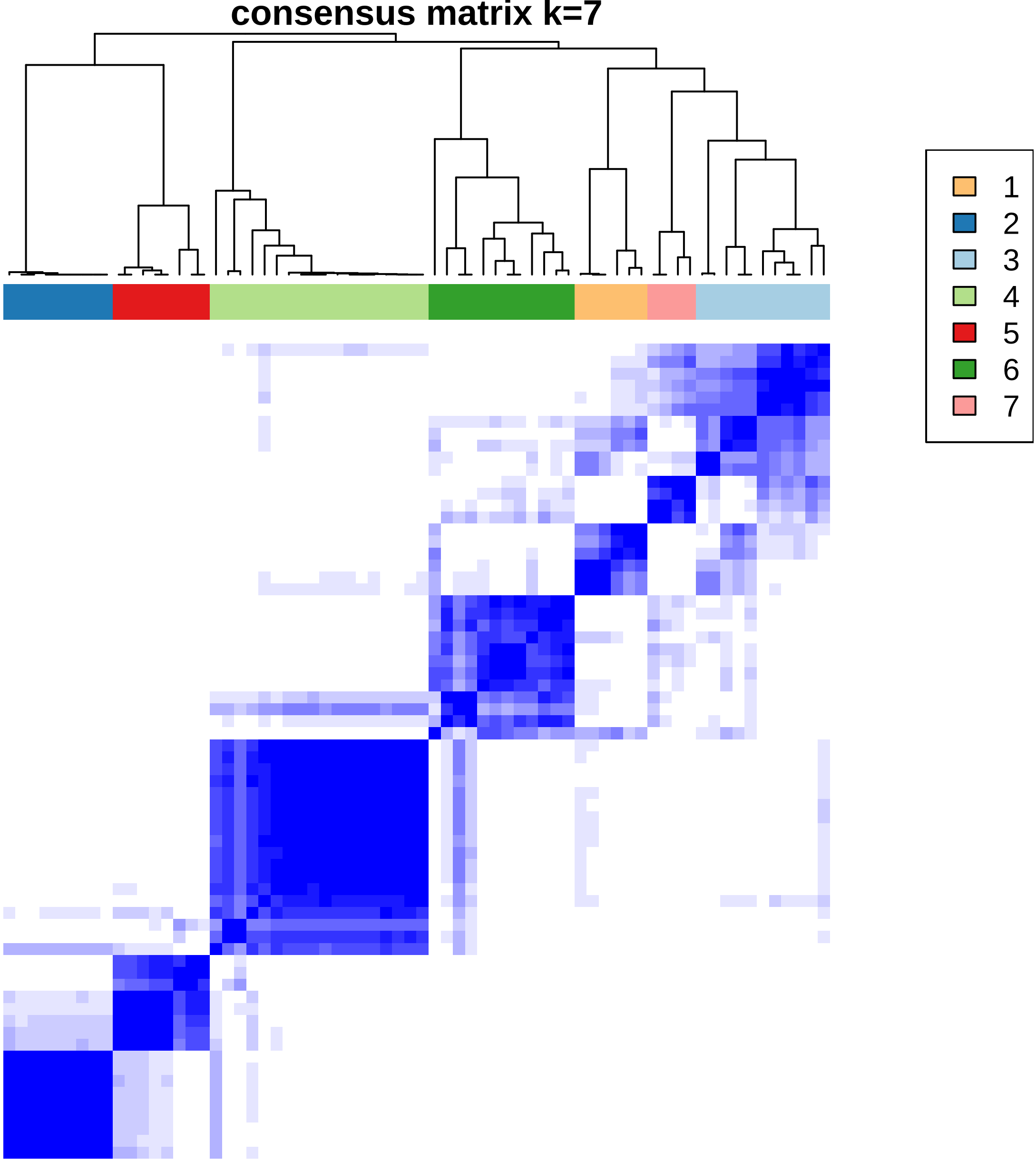
consensus matrix k=7
1
2
3
4
5
6
7

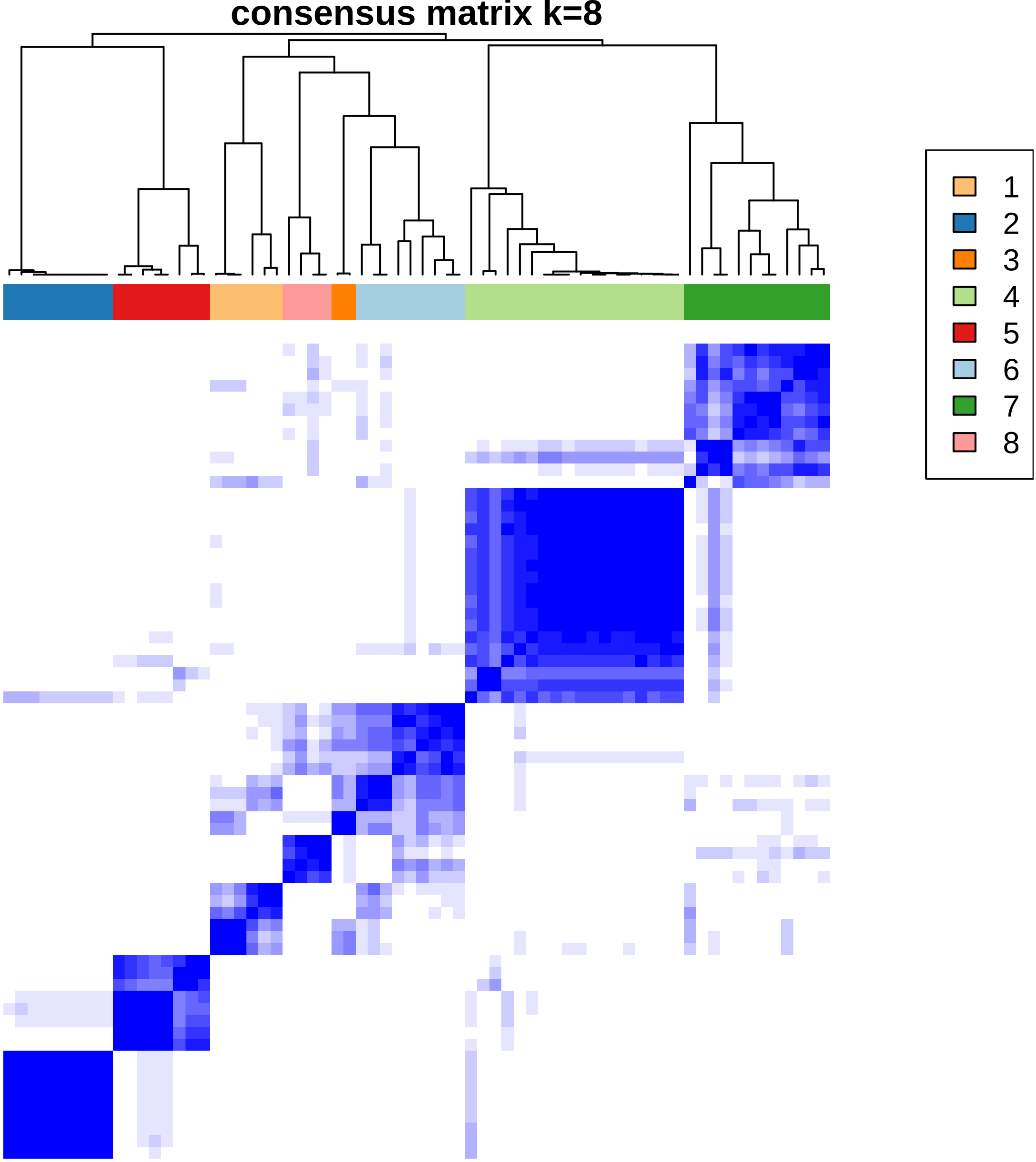
consensus matrix k=8
1
2
3
4
5
6
7
8

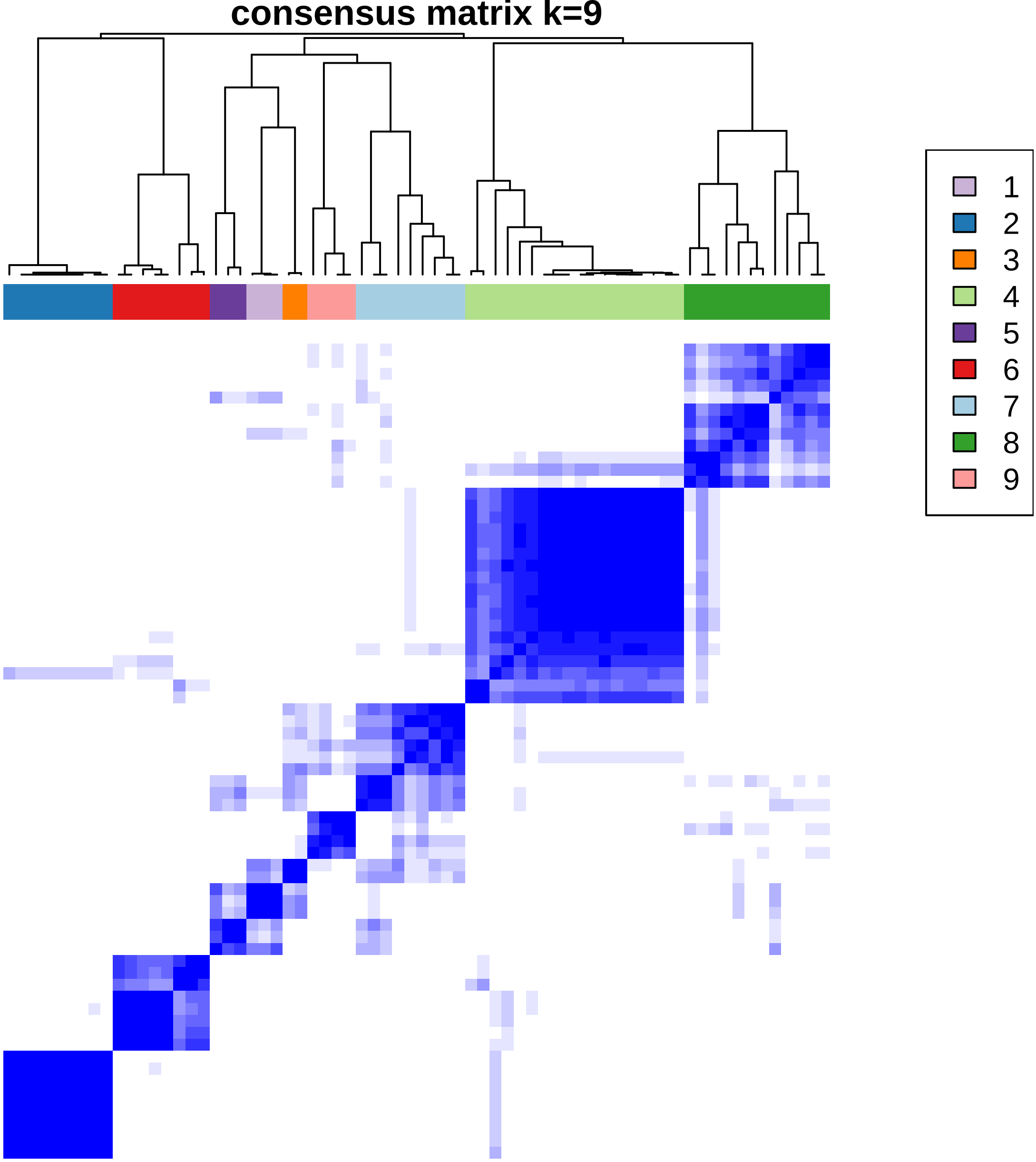
consensus matrix k=9
1
2
3
4
5
6
7
8
9

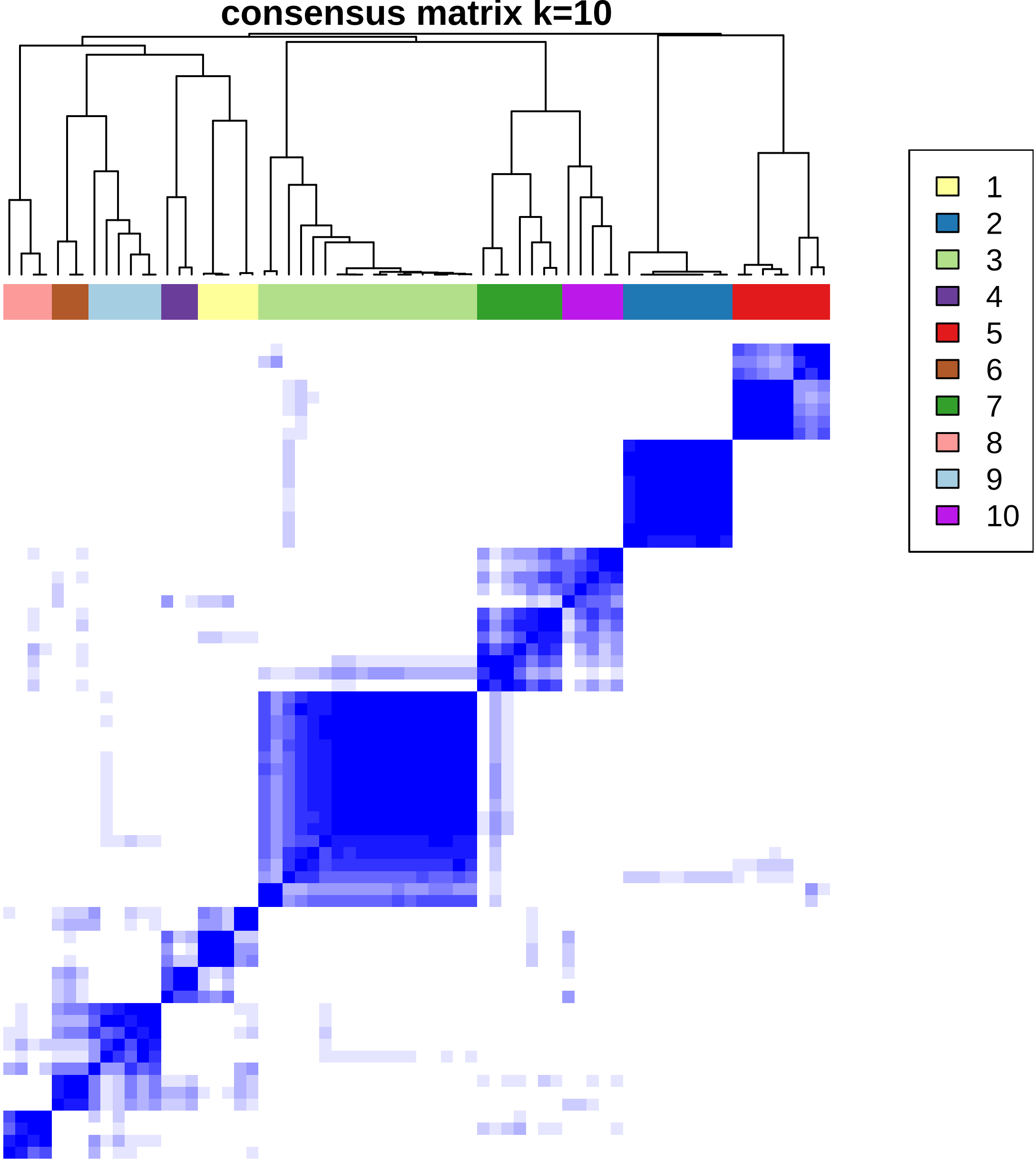
consensus matrix k=10
1
2
3
4
5
6
7
8
9
10

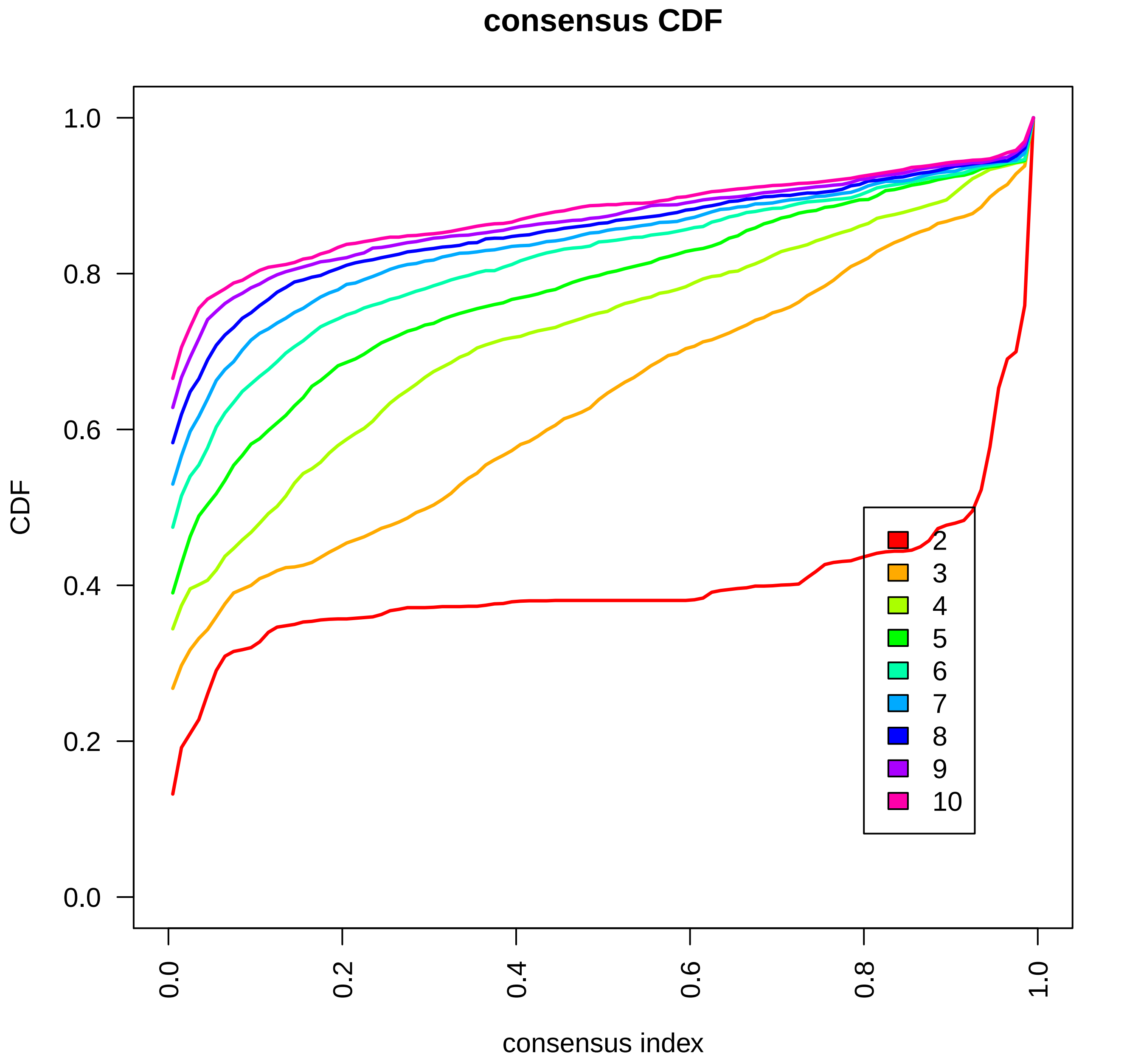
consensus CDF
CDF
consensus index
1.0
0.8
0.6
0.4
0.2
0.0
0.0
0.2
0.4
0.6
0.8
1.0
2
3
4
5
6
7
8
9
10

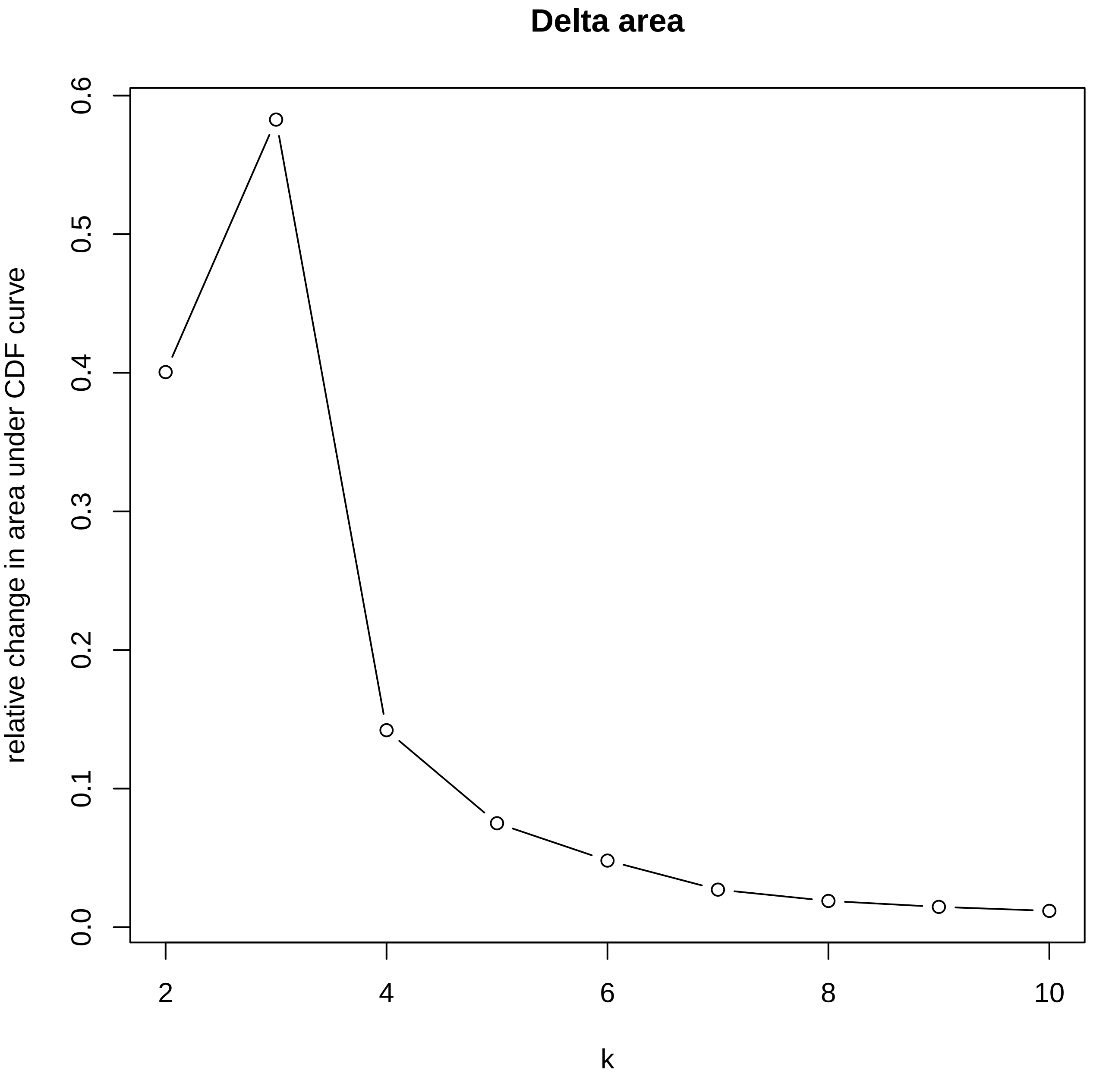

Delta area
relative change in area under CDF curve
0.0
0.1
0.2
0.3
0.4
0.5
0.6
2
4
6
8
10
k

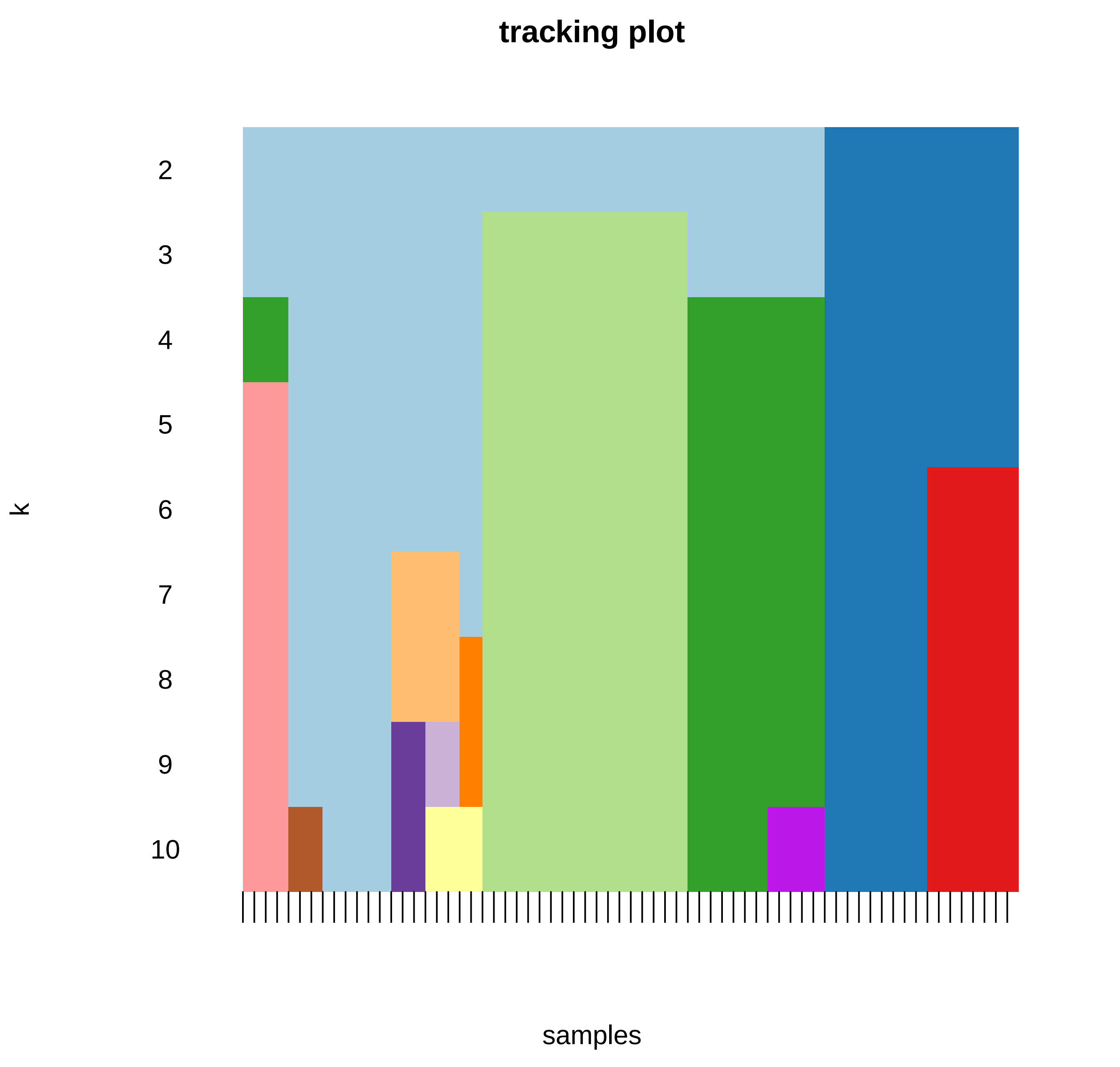

tracking plot
k
2
3
4
5
6
7
8
9
10
samples
